# Cryo-EM structures of the Plant Augmin reveal its intertwined coiled-coil assembly, antiparallel dimerization and NEDD1 binding mechanisms

**DOI:** 10.1101/2025.02.25.640204

**Authors:** Md Ashaduzzaman, Aryan Taheri, Yuh-Ru Julie Lee, Yuqi Tang, Fei Guo, Stephen D. Fried, Bo Liu, Jawdat Al-Bassam

## Abstract

Microtubule (MT) branch nucleation is fundamental for building parallel MT networks in eukaryotic cells. In plants and metazoans, MT branch nucleation requires Augmin and NEDD1 proteins which bind along MTs and then recruit and activate the gamma-tubulin ring complex (γ-TuRC). Augmin is a fork-shaped assembly composed of eight coiled-coil subunits, while NEDD1 is a WD40 β-propellor protein that bridges across MTs, Augmin, and γ-TuRC during MT branch nucleation. Here, we reconstitute hetero-tetrameric and hetero-octameric Arabidopsis thaliana Augmin assemblies, resolve their subunit interactions using crosslinking mass spectrometry and determine 3.7 to 7.3-Å cryo-EM structures for the V-junction and extended regions of Augmin. These structures allowed us to generate a complete de novo plant Augmin model that reveals the long-range multi coiled-coil interfaces that stabilize its 40-nm hetero-octameric fork-shaped organization. We discovered the dual calponin homology (CH) domain forming its MT binding site at the end of its V-junction undertake open and closed conformations. We determined a 12-Å dimeric Augmin cryo-EM structure revealing Augmin undergoes anti-parallel dimerization through two conserved surfaces along Augmin’s extended region. We reconstituted the NEDD1 WD40 β-propellor with Augmin revealing it directly binds on top its V-junction and enhances Augmin dimerization. Our studies suggest that cooperativity between the Augmin dual CH domains and NEDD1 WD40 binding site may regulate Augmin V-junction dual binding to MT lattices. This unique V-shaped dual binding and organization anchors Augmins along MTs generating a platform to recruit γ-TuRC and activate branched MT nucleation.

## Introduction

Microtubule (MT) nucleation is essential for organizing the cytoskeletal networks^1,2^. Across eukaryotes, MT nucleation requires the highly conserved cone-shaped γ-tubulin ring complex (γ-TuRC), which nucleates nascent MTs by templating their tube-like, thirteen protofilament organization^3,4^. The γ-TuRCs nucleate MTs either from MT organizing centers, or as branches alongside polymerized MTs^1,2^. In animal cells, the γ-TuRCs localize to centrosomes and nucleate most MTs during interphase, leading to a centralized MT cellular MT network with polymerizing dynamic MT plus-ends extending to the cell periphery, while MT minus ends are anchored to γ-TuRC in the centrosome^1,2^. In contrast, in plant cells, which lack centrosomes, γ-TuRCs localize along existing MTs and nucleate MT branches to form a near parallel peripheral MT network^5,6^. In both plant and animal mitotic cells, γ-TuRCs are recruited to bind along mitotic spindle MTs and produce parallel MTs extending towards chromosomes in the mitotic spindle^7^. Augmin is required for centrosome-independent γ-TuRC activated MT branch nucleation in mitosis leading parallel MTs to form bipolar mitotic spindles. The activities of Augmin and <u>N</u>eural precursor cell <u>e</u>xpressed, <u>d</u>evelopmentally <u>d</u>ownregulated 1 (NEDD1) are necessary for aligning and segregating chromosomes during cell division, and the defects in its eight subunits lead to short and thin mitotic spindles^7,8^. *D. Melanogaster* RNAi screens for mitotic phenotypes identified a subset of the eight Augmin subunits as essential for γ-tubulin association with spindle MTs and producing parallel MT arrays in mitosis ^9^. In contrast to the primarily mitotic functions of Augmin in animals, plant Augmin promotes γ-TuRC mediated MT-branch nucleation in interphase and mitosis^5–7^.

Recombinant γ-TuRCs are weak MT nucleators, as other conserved factors are needed to recruit, anchor and activate them at centrosomes or along dynamic MTs ^10–12^. Augmin is among the most conserved γ-TuRC associated factors across plants and animals^13^. The metazoan Augmin complex consists of eight unique coiled-coil subunits, termed HAUS 1,2,3,4,5,6,7,8^14^. Eight equivalent plant Augmin subunits, named AUG 1,2,3,4,5,6,7,8, were identified in *Arabidopsis thaliana*, revealing their essential function for MT-branch nucleation during both interphase and mitosis^15–17^.

Plant genomes, however, include eight unique AUG8 paralogs, suggesting a diversity of Augmin functions in mediating MT branch nucleation in plant cells^7,16^. In interphase, branched MTs emerge on average at 40° (incident angle) from the polymerized MTs, whereas in mitosis, branched MTs emerge at 10°, leading to a mostly parallel MT array suggesting a diversity of Augmins with unique mitotic and interphase specified by unique AUG8 adaptors^7,16^.

In addition to Augmin, other MT associated proteins (MAP) regulate Augmin MT association. TPX2 has been shown to recruit Augmin to MTs in *Xenopus laveis* extracts and is conserved in both plants and animals^18^; however, its deletion is totally dispensable in plants^19^. In contrast, NEDD1, also termed GCP-WD, is a highly conserved MAP across plants and animals, and its defects critically impact MT nucleation function in both systems^18,20^. NEDD1 consists of a WD40 containing β-propellor domain and a C-terminal helical coiled-coil^20,21^. Human mutations in NEDD1 or Augmin are linked to neurological disorders and are dysregulated in neural precursor cells^22–24^. Similarly, defects in NEDD1 and Augmin are lethal in plants suggesting critical roles and mechanisms in MT nucleation^7,25^. Live cell imaging in *Drosophila* cells during anaphase show that Augmins first bind to MTs followed by γ-TuRC recruitment to nucleate daughter dynamic MTs^26^. A recent study suggested NEDD1 recruits Augmin to bind to MTs, and then recruit γ-TuRC to activate MT branch nucleation^20^. Furthermore, these studies suggest that Augmins oligomerize upon binding MTs prior to recruiting γ-TuRC^20^. However, the nature of the oligomerization remains completely unknown. The structural mechanisms of Augmin and NEDD1 in regulating γ-TuRC activation along dynamic MTs have mostly remained poorly understood, in part, due to lack of reconstitution studies of these systems.

Augmin assemblies are hetero-octamers composed of 30-nm extended region that end with a 10-nm wide V-shaped junction (V-junction), resembling the shape of a “tuning fork”. Biochemical studies suggest the Augmin V-junction binds MTs via its conserved dual-headed calponin homology (CH) MT binding domain in HAUS 6,7 (AUG 6,7) and positively charged disordered N-terminal region of Haus8 (AUG8), all of which reside at the tip of the long arm of the V-junction^14^. However, it is not clear how the second end of the V-junction stabilizes Augmin binding to MTs. The Augmin dual CH-domains have been compared to the well-studied NDC80/Nuf2 kinetochore complex, which contains a similar dual MT binding CH-domains^27,28^.

The conformation of the dual CH-domains upon MT binding remains unknown. It also remains unknown how Augmins anchor along MTs via their V-junctions, and recruit γ-TuRC via their extended region. Recent reports of multiple low-resolution Augmin cryo-EM structures, in combination with AlphaFold 2/ColabFold models, have led to structural models for the eight coiled-coil assembly, suggesting a general view of the hetero-octameric organization^29–31^. However, among the clearest of these cryo-EM maps, low-resolution α-helical density is observed, while most of the coiled-coil assembly interactions were inferred by placing AlphaFold2 models into the low-resolution cryo-EM density maps. Difficulties in studying the structures of Augmin structures stem from their extremely elongated shape and flexibility, hindering crystallographic or cryo-EM structure determination studies.

Our study presents a comprehensive structural and biochemical analysis of the Augmin assembly. By reconstituting recombinant *Arabidopsis thaliana* Augmin hetero-octameric and hetero-tetrameric assemblies, we utilized crosslinking mass spectrometry and single particle Cryo-EM to determine structures for different regions of the Augmin particle leading to a near complete *de novo* Augmin model. Our structural analysis reveals insights into the dynamic flexibility and states of Augmins such as the conformation of AUG6,7 dual CH-domains at the tip of the V-junction, which adopt both splayed and packed states. Furthermore, we observed that Augmins undergo anti-parallel dimerization, mediated by conserved interfaces along their extended regions. This organization was visualized in a 12-Å cryo-EM structure of the Augmin dimer assembly, highlighting the spatial arrangement and transitions of the extended domain. Our findings also demonstrate that NEDD1 WD40 β-propellor binding to Augmins requires their V-junction region, and it enhances Augmin dimerization. The 12-Å Cryo-EM structure of the Augmin-NEDD1 β-propeller revealed its binding site on top of the V-junction. Our results lead us to a structural model wherein Augmin binding to MTs is stabilized by AUG6,7,8 and the NEDD1 β-propeller, positioned on different ends of the V-junction to anchor Augmin along multiple MT-protofilaments. This arrangement likely creates a platform for anchoring γ-TuRC, thereby facilitating MT branch nucleation. Our work significantly advances the understanding of Augmin’s role in MT dynamics, providing detailed molecular insights into its interactions and structural organization.

## Results

### Biochemical reconstitution and characterization of Plant Augmin assemblies

To purify *Arabidopsis thaliana* Augmin assemblies, coding regions for AUG1,2,3,4,5,6,7,8 subunits were assembled into polycistronic bacterial expression vectors (**Figure S1**). The AUG1, AUG3, AUG4, and AUG5 subunits consist mostly of highly conserved α-helices, while AUG2, AUG6, AUG7, and AUG8 were relatively less conserved (**Figure 1A**). The MT binding region of AUG8 is highly divergent and predicted to be disordered or unstructured and was thus excluded from expression (**Figure 1A**). To overcome many rare Leu and Arg plant codons, codon optimized AUG2,7,8 sequences were used for bacterial expression. We generated three polycistronic co-expression vectors consisting of either eight subunits (AUG1,2,3,4,5,6,7,8) or four subunits (AUG1,3,4,5 or AUG2,6,7,8) (**Figure S1**). Polycistronic expression led to soluble assemblies of AUG1,2,3,4,5,6,7,8 and AUG1,3,4,5 (**Figure S1**). However, AUG2,6,7,8 assembly were unstable and could not be purified, suggesting that AUG2,6,7,8 require the AUG1,3,4,5 for their solubility and can only be studied as a part of the full Augmin (AUG1,2,3,4,5,6,7,8). Mass spectrometry confirms each Augmin subunits is purified in the assembly and that the AUG6 C-terminal region is prone to degradation with multiple polypeptide bands (**Figure S1E**; **Figure 1A**). Mass spectrometry also identified several contaminating β-barrel containing proteins, suggesting these copurify with Augmin through multiple steps (Supplementary list file). We measured the masses of these Augmin assemblies using size exclusion chromatography with multi-angle light scattering (SEC-MALS) and mass photometry (MP) revealing AUG1,3,4,5 are hetero-tetramers of 250-280-kDa mass, and the AUG1,2,3,4,5,6,7,8 assemblies form a hetero-octamers of ~380-420 kDa mass (**Figure 1B, D; Figure S1D, H**). The measured masses suggest a stoichiometry of one Augmin subunit per Augmin assembly, consistent with human Augmin^14^. Although the plant AUG1,2,3,4,5,6,7,8 hetero-octamers are stable, changes in ionic strength led to their destabilization through a cascade of AUG2,6,7,8 dissociation likely caused by the degradation or dissociation of AUG6, resulting a mixture of hetero-octamers (AUG1,2,3,4,5,6,7,8) and hetero-tetramers (AUG1,3,4,5) over time (see **Figure S7F**).

**Figure 1:**
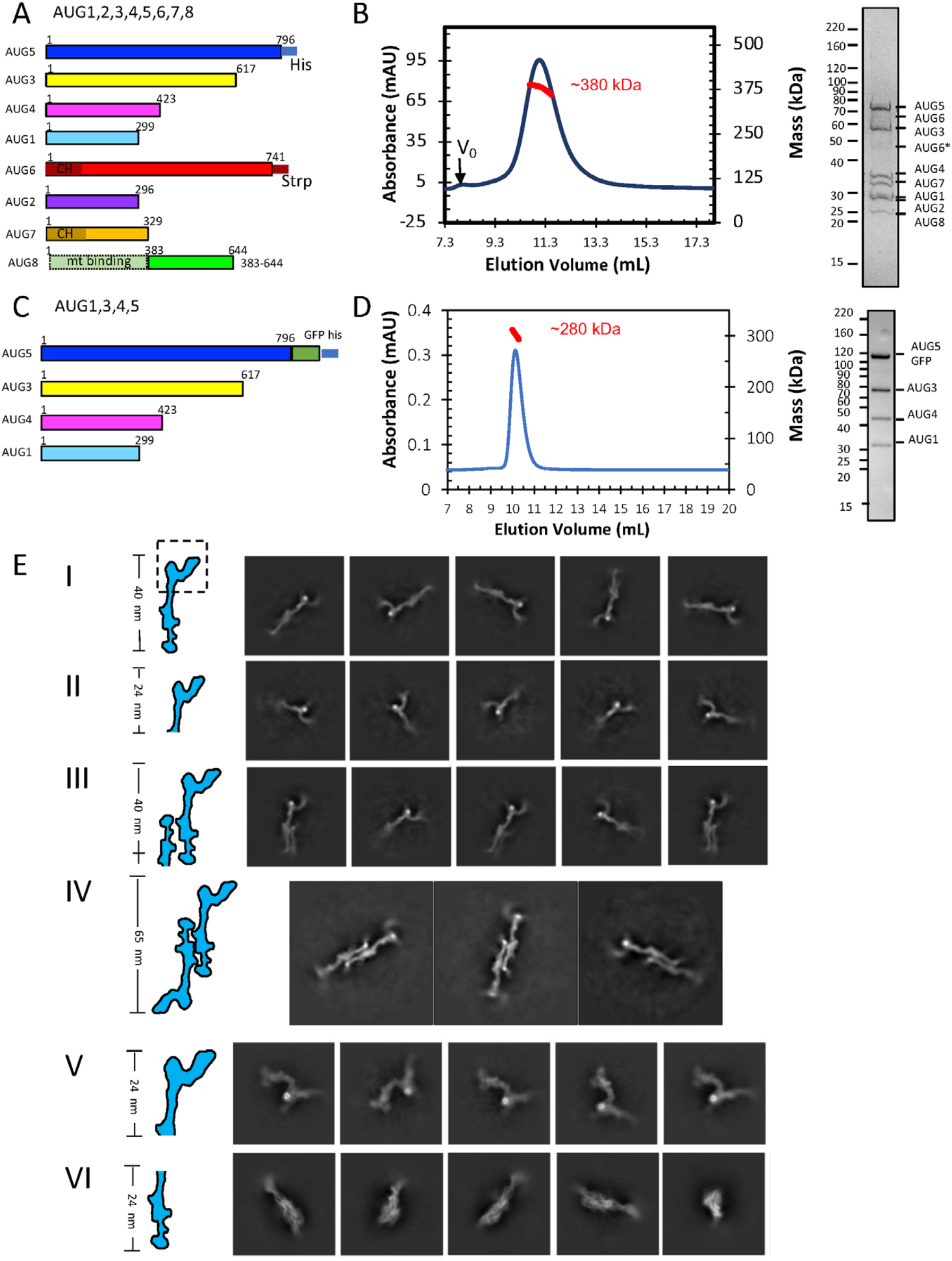
Biochemical and Structural characterization of *A. thaliana* Augmin assemblies. A) Schematic representation of plant Augmin hetero-octamer (AUG1,2,3,4,5,6,7,8) subunit organization, showing domain boundaries and purification tags. Note: AUG8 construct (residues 383-644) excludes the N-terminal MT binding domain (See **Figure S1**). B) Biochemical validation of hetero-octameric Augmin complex. Left: SEC-MALS analysis confirming monodisperse assembly with predicted octameric mass. Right: SDS-PAGE of purified complex showing all eight subunits (See **Figure S1**). C) Schematic of minimal Augmin hetero-tetramer (AUG1,3,4,5) subunit organization, including domain boundaries and purification tags. D) Biochemical validation of hetero-tetrameric complex. Left: SEC-MALS analysis demonstrating monodisperse assembly with tetrameric mass. Right: SDS-PAGE confirming presence of four subunits. E) Cryo-EM 2D-class averages revealing Augmin complex architecture. I: Monomeric AUG1,2,3,4,5,6,7,8: 40 nm tuning fork structure. II: Monomeric AUG1,2,3,4,5,6,7,8: V-junction focus with extended region. III: Dimeric AUG1,2,3,4,5,6,7,8: Two extended regions, single V-junction. IV: Dimeric AUG1,2,3,4,5,6,7,8: Two extended regions, two V-junctions. V: Focused V-junction from AUG1,2,3,4,5,6,7,8: 24 nm V-junction and stem regions. VI: Monomeric AUG1,3,4,5: 23 nm extended region.

### Cryo-EM imaging of Augmin reveal dynamic architecture and higher order oligomers

We collected cryo-EM datasets for hetero-octameric (AUG1,2,3,4,5,6,7,8) and Hetero-tetrameric (AUG1,3,4,5) Augmin assemblies (**Figure S2; Figure 1E**). 2D-class averages show that the AUG1,2,3,4,5,6,7,8 particles have a characteristic 40-nm tuning-fork shape, similar to human Augmin (**Figure 1E, panel I**)^29–31^. 2D-class averages also reveal a 30-nm extended region with a wide center, a narrow leg-like extension, and a foot-like density at one end, which closely resembles the shape of the AUG1,3,4,5 particles (see below). The other end forms a V-junction shape, leading to a second arm with a globular end (**Figure 1E, panel II**). The V-junction shows a high electron density spot, suggesting it is bound to a globular mass, which maybe potentially associated with nucleic acids (**Figure 1E**). In many cases, the 2D-class averages show the crescent-shaped V-junction without the 23-nm extended region, suggesting this region may be either broken off or out of focus in those images (**Figure 1E, panel II**). 2D-classes of 45-nm particles reveal dual extended region densities with a single V-junction (**Figure 1E, panel III**). Finally, 2D-classes of 60-nm particles display dimeric, C2-symmetric Augmin particles with dual anti-parallel tuning forks, indicating that Augmin dimers form in an anti-parallel manner via their respective central and foot regions (**Figure 1E, panel IV**). These diverse types of 2D-class averages suggest that these AUG1,2,3,4,5,6,7,8 particles have a propensity for dimerization, with some particles potentially losing the V-junction, which aligns with the biochemical interpretation of Augmin dissociating into sub-complexes. However, nearly half of the AUG1,2,3,4,5,6,7,8 particles are missing the extended regions (**Figure E, panel II**).

The 2D-class averages for the AUG1,3,4,5 assemblies show ~24-nm elongated filament-like particles with clear secondary structural elements. These particles have a wide profile near their center, a narrow leg-like density connected to a foot-like domain at one end, and an elongated short extension at the other end (**Figure 1E, panel VI**). The 2D-classes of AUG1,3,4,5 particles exhibit similar features to the lower extended region of the AUG1,2,3,4,5,6,7,8 particles (**compare Figure 1E, panels I and VI**). These particles are visible from multiple angles, including end-on views (**Figure 1E, panel VI**; **Figure S2**). These observations suggest that the foot region is mobile and flexible in the cryo-EM images, despite the more ordered structure of the extended region.

### Single particle cryo-EM structures of full Augmin and its different regions

2D-class averages of the hetero-octameric Augmin particles (AUG1,2,3,4,5,6,7,8) show a mixture of mostly side and top views (**Figure 1E, panels I and II**). As a result of the low signal to noise due to the large box sizes and extensive flexibility, we were only able to determine a consensus 9-Å Augmin structure (**Figure S2**). This reconstruction shows characteristic features of previously seen for the hetero-octameric human Augmins (**Figure S2, lower left**)^29–31^. Heterogeneity analysis allowed us to determine a low-resolution reconstruction for the Augmin particles with a second extended section, termed Augmin 1.5, revealing an incomplete Augmin dimer particle (**Figure S2, lower middle**). We also isolated the Augmin dimer particles and applied C2 symmetry leading to a 12-Å resolution reconstruction of the central core dimer interface, termed the Augmin dimer. This structure reveals in more detail how the extended regions transition in dimerization (**Figure S2, lower right; see below**).

To determine the single particle structures of the AugminV-junction-stem region with greater clarity, we combined all particles with the Augmin V-junction by recentering and re-extracting this region from the AUG1,2,3,4,5,6,7,8 datasets, and processing them as described in the scheme presented in Figure S3. The flexibility analyses revealed variable rotation of up to 201 in the long arm of the Augmin V-junction (**Figure S3; Figure S7A-C**). We determined two reconstructions for this V-junction-stem region: 1) A 7.3-Å resolution reconstruction representing a state in which the dimeric CH domains are tightly packed, which we termed the “closed state” (**Figure 2A, B, left and middle panel; Figure S3, lower left**). 2) A 10-Å resolution reconstruction representing a state in which the CH domains are splayed apart, which we termed the “open state” (**Figure 2A, right panel**). In the open state, the bow density rotates downward by ~15° with respect to the base of the V-junction. In the closed state, the V-junction density rotates upward by 15° compared to the open states. Vectorial comparisons of the residue-to-residue movements for models of the two states show that AUG2,6,7,8 moves upward while the AUG3,5 in the base also moves upward (**Figure S7C**). The 7.3-Å cryo-EM map of the closed state shows clear helical connectivity in the base of the V-junction, and that the closed dual-pronged packed CH domains are visible at the end of the V-junction (**Figure 2B, left and middle panel; Figure S5D-I**). In contrast, the 10-Å cryo-EM map of the open state shows the rotation of the V-junction and the opening of the dual-pronged CH-domains (**Figure 2B, right panel; Figure S6;** see below). These structures reveal the transitions of the V-junction and the globular head domain of AUG6,7 suggesting that their dual CH-domains undergo an open and close transition (**Figure S6H-J**). We determined a 3.7-Å cryo-EM structure of the hetero-tetrameric Augmin (AUG1,3,4,5) using the scheme presented in **Figure S4**. Our biochemical reconstitution of this minimal Augmin assembly led to a high resolution cryo-EM structure that reveals clear helical density and side chain interactions, previously not observed in the metazoan Augmin structures (Figure 2C; Figure S5J-L)29-31. The maps reveal the detailed folding organization of AUG1,3,4,5 subunits in the extended region of Augmin. The end of the AUG1,3,4,5 particle was flexible, and we thus utilized a combination of 3D-variability, flexibility and local refinement to obtain a 6-Å reconstruction for this region (**Figure S4, Figure S5M**). The previous Augmin structures have not resolved the terminal extended region, likely due to its flexibility, coupled with its consistently inaccurate prediction by AlphaFold2 models^29–31^. The helical features of this tripod-shaped terminal domain are clear, allowing for the accurate placement of and morphing of Alpha Fold models (**Figure 2D**; **Figure S5B-C, J-M**).

**Figure 2:**
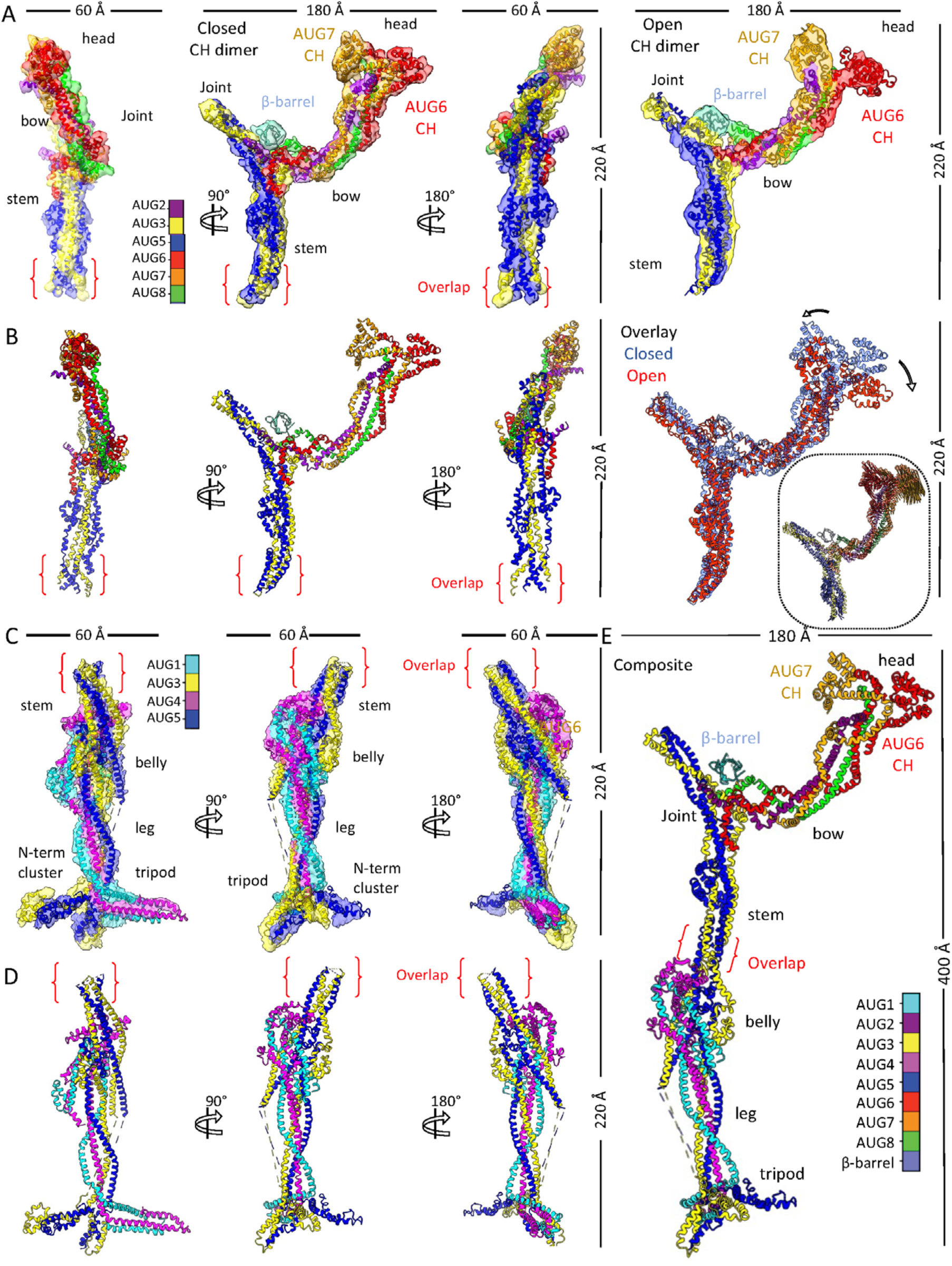
Single particle Cryo-EM structures and models of Augmin assemblies. A) Cryo-EM reconstructions of the AUG1,2,3,4,5,6,7,8 V-junction and stem. Left: 7.3-Å structure with segmented model of AUG1,2,3,4,5,6,7,8 (closed-state AUG6,7 CH-domain dimer). Right: 10 Å structure with open-state AUG6,7 CH-domain dimer (See **Figures S2-S4**). B) Model analysis. Left: AUG1,2,3,4,5,6,7,8 subunit organization in V-junction and stem. Right: Conformational comparison of closed (red) and open (blue) states. Inset: Vector representation of conformational transition. C) 3.7 Å reconstruction of AUG1,3,4,5, extended region with segmented model (See **Figures S2-S5**). D) Detailed subunit organization of AUG1,3,4,5 extended region model. Composite model of complete AUG1,2,3,4,5,6,7,8 complex derived from combined maps and models (See **Figures S5A-C**)

Using the overlap between the cryo-EM density maps of 7.3 to 3.7 Å resolution of the V-junction stem and the extended region and their fit into the consensus 10-Å Augmin particle cryo-EM structure, we are able to fully map helical densities across the 40-nm Augmin hetero-octamer particle (**Figure 2B, D, E; Figure S5A-B; video S1**). A globular density, representing a bright spot residing on top of the V-junction, was identified, which likely corresponds to the β-barrel domains of bacterial proteins co-purified with Augmin. The β-barrel was placed in this density and thus excluded from our model building and interpretation (**Figure 1E, panel V**; **Figure S5B, G**). These β-barrel proteins likely occupy a highly active binding site in Augmin, substituting a missing a binding partner for a protein in the bacterial expression system (**see below**) Our single particle cryo-EM structures lead to full density maps of all structured regions of the Augmin particle and allowed us to build all regions for the Augmin hetero-octamer (**video S1**).

### Complete *de novo* model of Augmin hetero-octamer reveals novel coiled-coil interfaces

Using our overlapping cryo-EM maps for the Augmin V-junction-stem and the extended regional, we were able to generate a *de novo* model of the Augmin hetero-octamer (**Figure 2E; Figure S5B-C; video S1**). The extended region is a 30 nm long assembly with a wide multi-helical region in its center, termed the “belly”, connected to a narrow four-helical bundle below, termed the “leg”, which terminates into a three-prong multi-helical density, termed the “tripod” (**Figure 2E; video S1**). On the other end of the belly region, a four helical bundle curved structure, we termed the “stem”, extends towards the V-junction leading to a nexus point that we term the “joint” (**Figure 2E; video S1**). Above the joint, a short extension extends upwards, in line with the direction of the stem. On the other side, a long four helical bundle that we termed the “bow”, extends in the other direction from the joint which ends with a globular region we termed the “head”, composed of the AUG6,7 CH domain dimer (**Figure 2E; video S1**).

Our Augmin model explains the multi-subunit coiled-coil helical interactions that stabilize the hetero-octamer and allow for flexibility at its terminal regions (**Figure S7**). The 30-nm extended region is composed of the hetero-tetrameric AUG1,3,4,5 assembly, while the V-junction are represented by AUG2,6,7,8 C-terminal regions assembled onto a platform composed of AUG3,5 folding back on themselves (**Figure 3A-B; Figure S5; Figure S7, Figure S8**). The belly region contains the AUG1,4 which binds the central N- and C-terminal regions of AUG3,5 (**Figure 3B**). The AUG1,3,4,5 C-termini form four helical bundles that supertwist together in the leg region and then fold their C-termini together in the tripod region **(Figure 2D; Figure 3B**). In the tripod, the AUG1,3,4,5 diverge into two subdomains where AUG1,4 fold into a larger spoke while the AUG3,5 C-termini form a second shorter spoke (**Figure 2D; Figure 3B**). The final and wider spoke in the tripod consists of the AUG3,5 N-terminal bundle, which extends from AUG3,5 N-terminal helices (**Figure 3B; Figure S5M**). The tripod region was not predicted accurately by AlphaFold (**Figure S8C**). From the top end of the belly region, the stem of the V-junction emerges from two sets of helices of the AUG3 and AUG5 with opposite topologies (**Figure 2E; Figure 3B; Figure S8C**). This region is a platform where AUG3,5 subunits “foldback zone” and assemble with AUG2,6,7,8 subunits to form the V-junction (**Figure 3B, top panels**). In the AUG1,3,4,5 assembly, the stem is not ordered, likely due to the destabilization of the AUG3,5 foldback zone in the absence of AUG2,6,7,8 (**Figure S7F; Figure S8C**). The AUG2,6,7,8 C-termini form four helical bundles that can be followed into the V-junction arm, including the AUG6,7 N-terminal dual CH domains (**Figure 3B**). Our model differs from previous AlphaFold models in that the helical regions of AUG3,5 foldback zone undergoes more extensive folding with AUG6 and AUG2 C-termini (**Figure 2A-B; Figure 3B; Figure S8A-B**).

**Figure 3:**
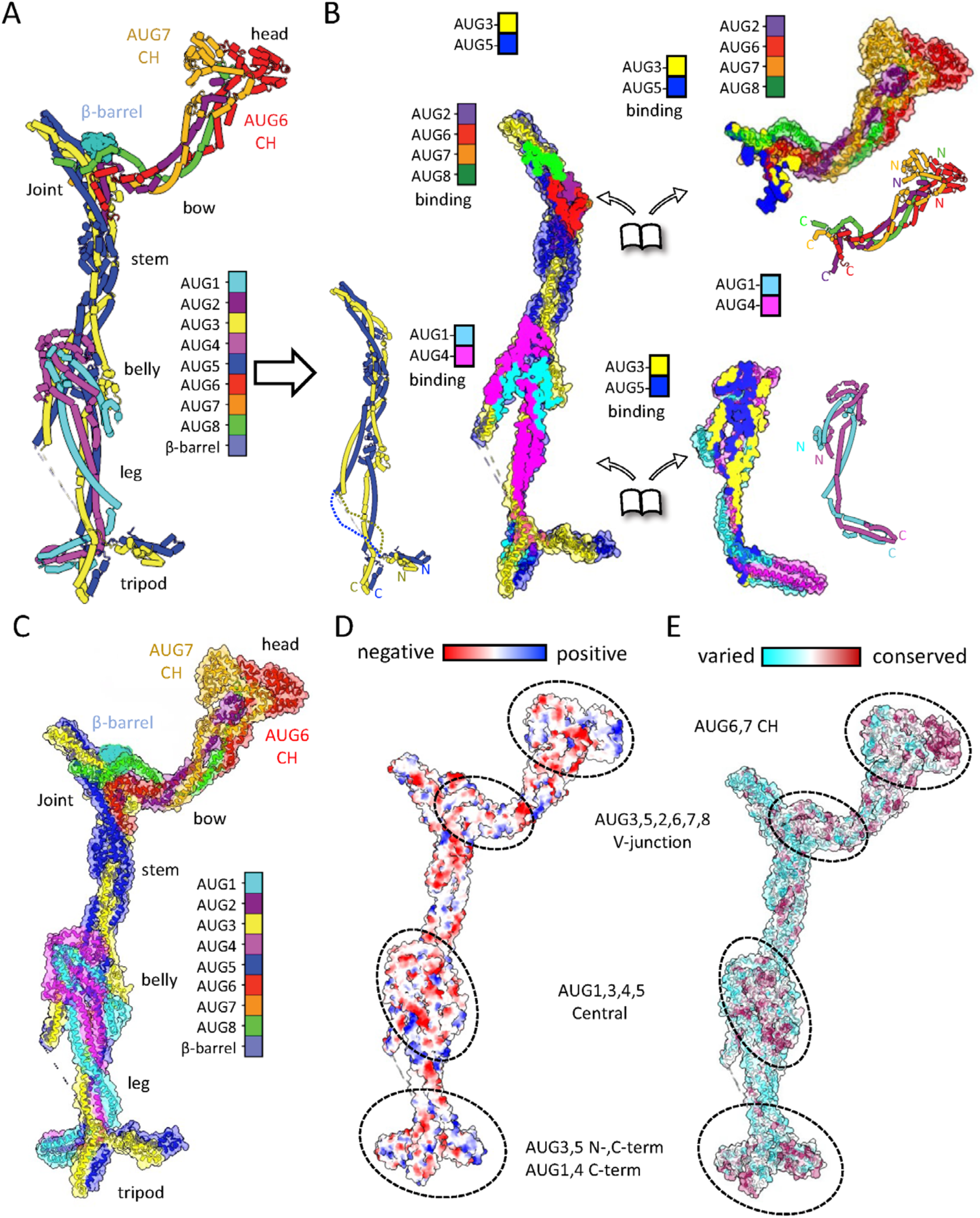
Architectural coiled-coil organization and conservation of the Augmin assembly. A) Ribbon diagram highlighting individual subunit folds and organization. B) Subcomplex interaction analysis. Left: AUG3,5 heterodimer with interaction footprints. Right: AUG1,4 and AUG2,6,7,8 subcomplexes with corresponding interfaces. Insets: Topological organization of each subunit. C) Complete Augmin hetero-octamer assembly showing structural domains. D) Electrostatic surface potential map highlighting functional regions (**Figure S9A-C**). E) Surface conservation analysis across plant, animal, and insect (**Figure S9F-H**).

### Augmin hetero-octameric structure stabilized by long and short range coiled-coil interactions

The structures reveal the full complex topology of all eight Augmin subunits (**Figure 3B**). The AUG3 and AUG5 are heterodimeric parallel long coiled-coil that form the backbone of the Augmin structure (**Figure 3B, left panel**). AUG1,4 subunits fold into the lower half of the extended region, and they stabilize their N- and C-terminal regions by folding in parallel with AUG3,5 N-term cluster in the leg while anti-parallel folding into against themselves into the belly region with both N and C-terminal central helices of AUG3,5, giving this region its girth with an eight helical bundle (**Figure 3B, lower left**). At the bottom of the extended region of Augmin, the tripod consists of a co-folded AUG1,4 C-terminal bundle as the longest spoke and AUG3,5 C-terminal bundle, which forms a second shorter spoke, and the AUG3,5 N-terminal cluster forming the third wider spoke of the tripod (Figure S5M; Figure 3B, lower panel). In the V-junction of Augmin, the AUG2 and AUG6 C-termini intertwine within the AUG3,5 foldback, forming a six helical bundle leading the stem to widen just below the V-junction **(Figure 2A-B; Figure 3B, top right**). The AUG2,6,7,8 C-termini stabilize the AUG3,5 helices in the foldback zone at the top V-junction on while their N-terminal domains towards the long end of the V-junction to form the head globular region (**Figure 3B, top left**). The two sets of N- and C-termini of AUG3,5 fold onto AUG1,4 foldback zone leading to the belly region in the center (**Figure 3B, lower panel**). The Augmin model reveals a mixture of parallel and anti-parallel intertwined coiled coils stabilized by short and long-range foldback interactions leading to its conserved organization (**Figure S8B, D; Figure 3B**).

We resolved two states of the AUG6,7 CH-domains in the head region: an open and closed state of the AUG6,7 CH domains (**Figure 2B, left panel; Figure S6G-J**). In the closed state, the CH domains are aligned laterally tightly against each other (**Figure 2A, left panel; Figure S6H**), and in the open state they are splayed apart (**Figure 2A, right panel; Figure S6G**). These transitions are coupled with a lateral rotation of the AUG2,6,7,8 helical bundle (**Figure 2B, right panel**). Taken together, the Augmin V-junction undergoes transitions continuously stretched throughout the structure where stabilization in one part would likely stabilize subsequent parts (**Figure S7**). Due to flexibility in this region, we were unable to determine a high-resolution density map of the V-junction (**Figure S7**).

### Four critical conserved surfaces can be identified on the Augmin structure

To understand the sequence conservation and charge distribution, we aligned sequences for each of the hetero-octameric subunits (AUG1,2,3,4,5,6,7,8) across two hundred orthologs from plants, animals, and insects. We plotted sequence conservation and compared those to a charge distribution plot along the surface of the Augmin model (**Figure S9, Figure 3D-E**). We observe extensive conservation in the coiled-coil helical interactions between subunits within various regions of the structure suggesting conserved long and short-range assembly interactions (**Figure S9A-C; Figure 3E**). Conservation analysis on the surface of the Augmin hetero-octamer reveals four conserved regions with unique charge distributions, likely indicating critical zones of protein-protein interactions (**Figure S9A-C, Figure 3D-E**). 1) A large surface on the backside of the belly region involving interfaces with AUG3,5 and the AUG1,4 N-terminal helical regions. 2) the AUG3,5 C-termini, and the AUG1,4 C-termini in the tripod (**Figure 3D-E**). 3) the top of the V-junction interfaces composed of the AUG3,5 central foldback zone interfacing with AUG2,6,7,8 C-terminal helical bundle (**Figure 3D-E**); and 4) AUG6,7 dual CH domain dimer in the head region, which mediates MT binding (**Figure 3D-E**). The third region is bound by the density which we assigned to the β-barrel density, which likely forms the binding site for NEDD1-β-propellor (**see below**)

**Augmin undergoes antiparallel dimerization via the tripod to belly regions interfaces**

2D-class averages of full Augmin hetero-octameric (AUG1,2,3,4,5,6,7,8) particles show side views of a “head-to-tail” Augmin homodimers bound to each other along their long axis (**Figure 1E, third panel; Figure S2, left panels**). 2D-class averages for the 1.5 Augmin particle show an identical “head-to-tail” dimer of extended regions but are missing a V-junction and stem regions from one of the two assemblies (**Figure 1E, panel IV**). We determined a 12-Å C2 symmetric anti-parallel cryo-EM single particle structure followed by extended region model placement and de novo model refinement (**Figure S2, left panels; Figure 4**). In this structure, the V-junctions and part of the stem are 30 nm away from the center and were plagued with higher flexibility and were thus excluded from our structures (**Figure S2, left panels**). Despite the 12 Å resolution of the central region structure, we were able to place the Augmin core models into each Augmin assembly in the head-tail dimer structure and understand the conformational transitions they undertake upon dimerization (**Figure 4B-C; Figure S10**).

**Figure 4:**
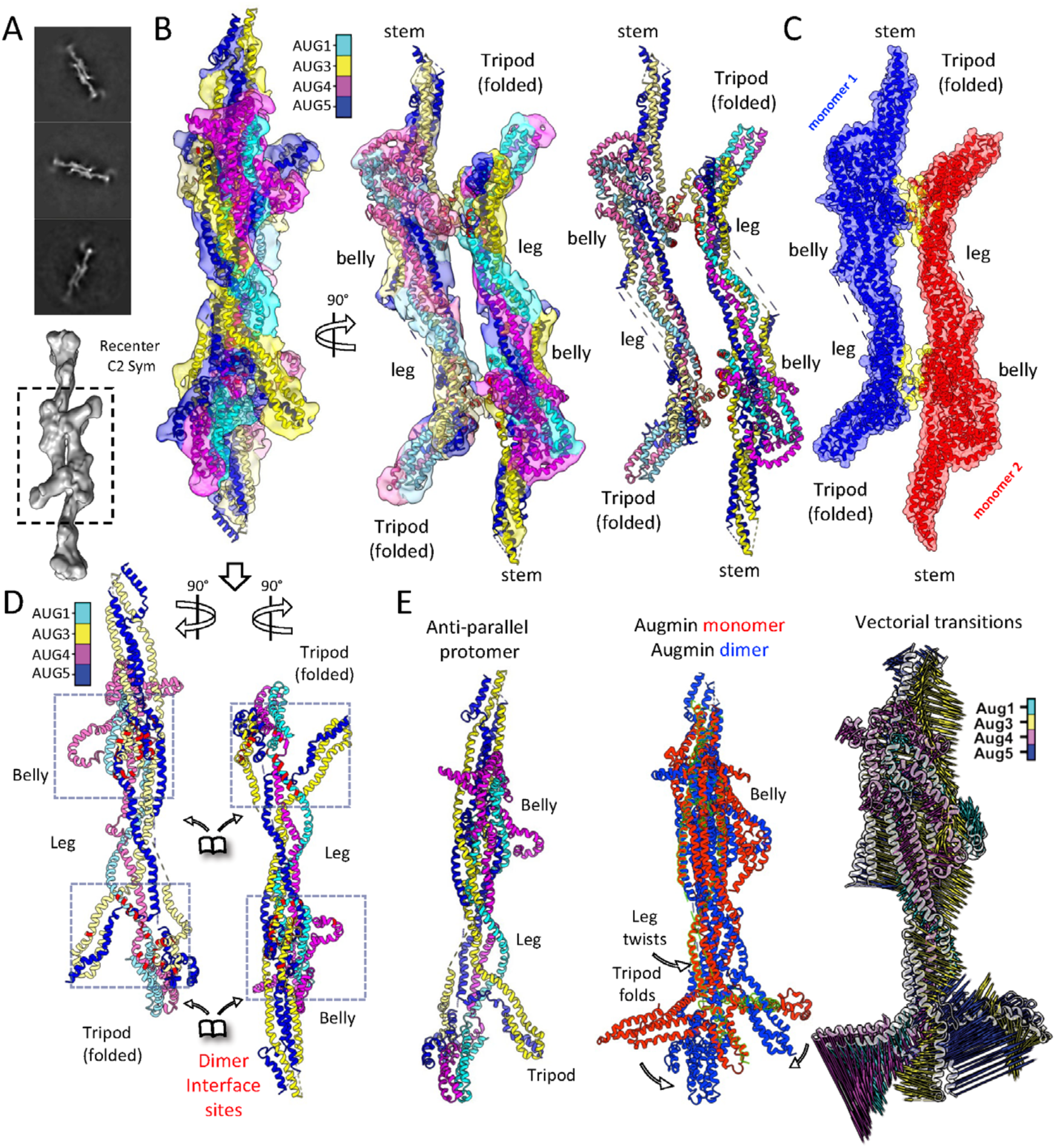
Cryo-EM structure and analysis of Augmin antiparallel dimer assembly. A) Top: 2D class averages showing antiparallel organization of extended domains. Bottom: Low-resolution reconstruction with focused refinement of dimerization interface. B) Detailed structural analysis of extended region dimer. Left: Side view of segmented reconstruction with colored subunit organization. Middle: 90° rotated view showing subunit arrangement. Right: Isolated model highlighting subunit organization at dimer interface. C) Dimer interface analysis showing protomer organization. Red/blue: individual protomers. Yellow: Interface region between protomers. D) Conformational changes in dimer formation. Splayed view of protomers showing AUG1,3,4,5 organization. Interface zones highlighted in red at foot and belly regions. Arrows indicate domain movements during dimerization. E) Monomer-to-dimer transition analysis. Left: Single protomer structure. Middle: Overlay of monomeric and dimeric states. Right: Vector representation of conformational changes in tripod, belly, and leg regions.

Dimerization is mediated by one face of the belly region and the back side of a rearranged tripod region (**Figure 4B-D; Figure S10**). In the Augmin dimer state, the tripod region undergoes a 20° rotational re-arrangement that repositions its three lobes to be facing outwards away from the dimer interface (**Figure 4E**). The AUG3,5 N-terminal cluster lies close to the dimerization site on the back of the AUG1,4 C-terminal lobe and binds to the belly domain of the second Augmin particle, which is antiparallelly oriented (**Figure 4D**). The belly region and the leg region both twist in their folds, accommodating the rearrangement in the dimeric Augmin state. The surfaces of the tripod and the belly regions mediating the head-to-tail Augmin dimerization were two of the four highly conserved regions on Augmin as noted earlier (**Figure 3C-E**). The Augmin dimer structure reveals the conformational transitions in the extended region during anti-parallel dimerization.

### Crosslinking mass spectrometry validate subunit interactions within the Augmin complex

To understand the multi-subunit coiled-coil interactions within the Augmin hetero-octamers, we carried out crosslinking mass spectrometry (XL-MS) for both hetero-tetrameric (AUG1,3,4,5) and hetero-octameric (AUG1,2,3,4,5,6,7,8) assemblies (**Figure 5; Figure S11; see Materials and Methods**). The XL-MS datasets revealed both intra-subunit and inter-subunit peptide crosslinks (**Figure 5; Figure S11A**). When we mapped all identified crosslinks onto the cryo-EM models for the Augmin hetero-tetramer (AUG1,3,4,5) and the Augmin hetero-octamer (AUG1,2,3,4,5,6,7,8), we found that 50% (105 out of 209) and 70% (80 out of 115) of these crosslinks had Cα–Cα distances below 30 Å. These data suggest that the hetero-tetrameric (AUG1,3,4,5) assembly is more dynamic, particularly in the tripod and stem regions due to the flexibility of the tripod and the lower stability of AUG3,5 foldback zone—a region not observed in the cryo-EM map (**Figure 5A; Figure S11B**). In contrast the hetero-octameric (AUG1,2,3,4,5,6,7,8) model explains the majority of the crosslinks, suggesting it is a more stable assembly, consistent with both our cryo-EM structures and biochemical studies (**Figure 5B; Figure S11C**). When we assessed how many crosslinks in the experiments on hetero-octameric (AUG1,2,3,4,5,6,7,8) Augmin can be explained when crosslinks are additionally allowed to span across assemblies in the model of the anti-parallel dimer, we found 75% (44 out of 59) of the crosslinks now have Cα–Cα distances below 30 Å, suggesting that some of the Augmin assemblies are in the anti-parallel dimeric state (**Figure S11C**).

**Figure 5:**
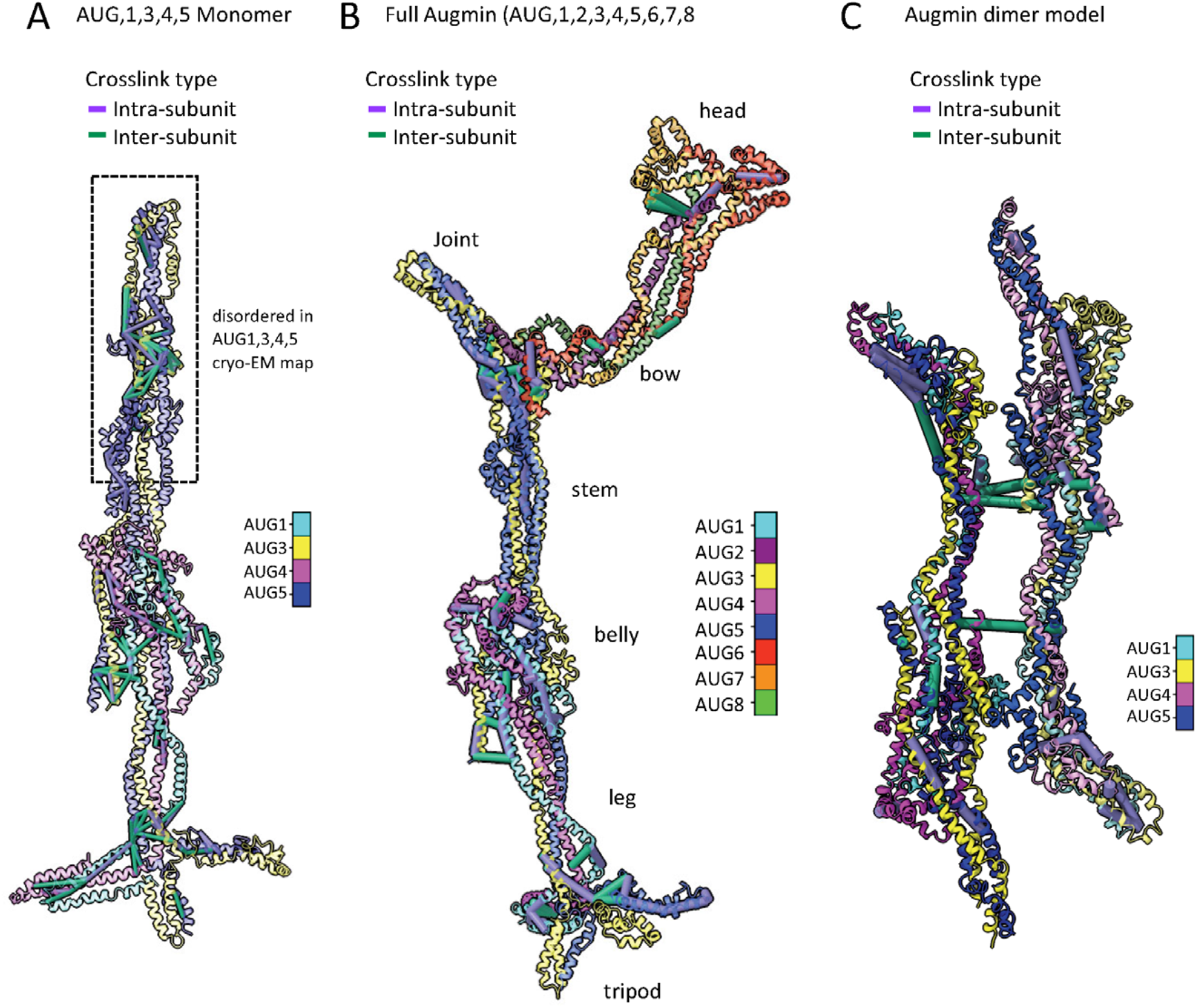
XLMS validates features of Augmin hetero-octamer and Antiparallel Augmin dimer. A) XLMS analysis of hetero-tetrameric (AUG1,3,4,5) Augmin with mapped crosslinks on the cryo-EM generated model. This model of the flexible fold-back zone in AUG3,5, which is absent in the cryo-EM map, but is modeled from (AUG1,2,3,4,5,6,7,8) structure (See **Figure S11**). B) Comprehensive XLMS analysis of hetero-octameric (AUG1,2,3,4,5,6,7,8) Augmin model. Validated interactions throughout complex. Emphasis on joint and tripod zones. C) Crosslink mapping of AUG1,2,3,4,5,6,7,8 anti-parallel dimer assembly validating interface regions. For more details see **Figure S11**

Plotting the AUG1,3,4,5 XL-MS residue pairs identified on the AUG1,3,4,5 model reveals extensive close-range inter and intra-subunit interfaces that are consistent with the folding of the AUG1,3,4,5 subunits in the belly region, leg region, and most crucially the tripod region (**Figure 5A; Figure S11D-E**). In the tripod region, intra-subunit and inter-subunit interfaces can be observed in all three legs corresponding to the AUG1,4 and AUG3,5 C-termini and AUG3,5 N-termini (**Figure 5A; Figure S11D-E**). We observe extensive crosslinks in the AUG3,5 central foldback zone, even though this region was not resolved in the AUG1,3,4,5 cryo-EM structure (**Figure 5A; Figure S8C; Figure S11D-E**) suggesting this region is folded, but too flexible in the absence of AUG2,6,7,8 to be resolved in cryo-EM structures (**Figure 2E, Figure S8C; Figure S11D-E**). There were also many longer distance crosslinks between the stem and other parts of the structure, which may suggest the stem undergoes larger scale movements. The AUG1,2,3,4,5,6,7,8 XL-MS data were plotted on the full Augmin model (**Figure 5B; Figure S11F-H**). We observe 20 inter and intra-subunit crosslinks in the region forming the V-junction (**Figure 5B; Figure S11F-H**). Six AUG2,6,7,8 crosslinks are observed in the bow region and six more AUG6,7 crosslinks are observed in the dual CH domains (**Figure 5B; Figure S11F-H**). At least 4 unique residue pairs representing the head-to-tail Augmin antiparallel dimer cross-subunit crosslinks (**Figure 5C; Figure S11C**). In addition, we identified the crosslinks that differ in fitting the Augmin hetero-octamer anti-parallel dimer model which don’t fit the monomer model to be between the tripod and leg regions (Figure S11G). The tripod and leg regions undergo conformational change and become closer in the Augmin anti-parallel dimer model (Figure 4; Figure 5C). In summary, plots of our detailed XLMS crosslinks on the *de novo* Augmin model are fully consistent with the critical and distinguishing features of our structural model and conformational transitions: such as the folding of AUG1,4 and AUG3,5 in the tripod region, Antiparallel coiled-coil folding in the belly region between the N-termini of AUG1,4 and AUG3,5 central regions (**Figure 5C; Figure S11D-E**). Thus, these XLMS data validate the new Augmin *de-novo* model and support antiparallel dimerization of the extended region via the conserved interfaces in the belly and tripod regions (**Figure 3E; Figure 5B-C**).

### NEDD1 WD40-β-propeller binds Augmin assemblies that contain AUG2,6,7,8 subunits

We next reconstituted the NEDD1 WD40 β-propellor with the hetero-tetrameric (AUG1,3,4,5) and hetero-octameric (AUG1,2,3,4,5,6,7,8) Augmin assemblies to define the location of the NEDD1 Augmin binding site (Figure 6A-B). Full-length *A. thaliana* NEDD1 was insoluble in bacteria or insect cells. In contrast, the conserved N-terminal NEDD1-WD40 β-propeller (residues 1 to 315) is soluble when expressed in insect cells and purified as monodisperse protein (**Figure 6A-B; Figure S12A-B**). The NEDD1-WD40 β-propellor was previously observed bind MTs and synergize with Augmin^32^. We first used sucrose density gradients to explore NEDD1 WD40 β-propellor binding to AUG1,3,4,5 and AUG1,2,3,4,5,6,7,8 (**Figure 6B; Figure S12C-H**). The AUG1,2,3,4,5,6,7,8 bound NEDD1-WD40 β-propellor and they both co-elute at high molecular weight with Augmin subunits, with excess NEDD1-WD40 eluting at lower molecular weight (**Figure S12-F-H**). In contrast, NEDD1-WD40 β-propellor did not co-elute with AUG1,3,4,5 (**Figure S12C-E**). Thus, the NEDD1 β-propellor binding site is missing in the AUG1,3,4,5, but is present in AUG1,2,3,4,5,6,7,8 (**Figure 6B; Figure S12C-E**). We validated the AUG1,2,3,4,5,6,7,8 interaction with NEDD1 WD40-β-propellor using size exclusion chromatography (**Figure 6C-D**). Mass photometry on sucrose density purified NEDD1-WD40 β-propellor AUG1,2,3,4,5,6,7,8 compared AUG1,2,3,4,5,6,7,8 to suggest similar mass (450 vs 460 KDa); however, NEDD1-WD40 β-propellor binding induces Augmin dimerization (987 kDa), compared to its absence where Augmins behave mostly as monomers in solution (**Figure 12G-H**). These data are consistent with location of the NEDD1-WD40 β-propellor binding site to be only present in AUG1,2,3,4,5,6,7,8, and is missing in AUG1,3,4,5 suggesting it lies in the V-junction, which is composed of AUG2,6,7,8 (**Figure 6A-D; Figure S12A-F**). NEDD1 binding also mildly enhances the dimerization of Augmin particles.

**Figure 6:**
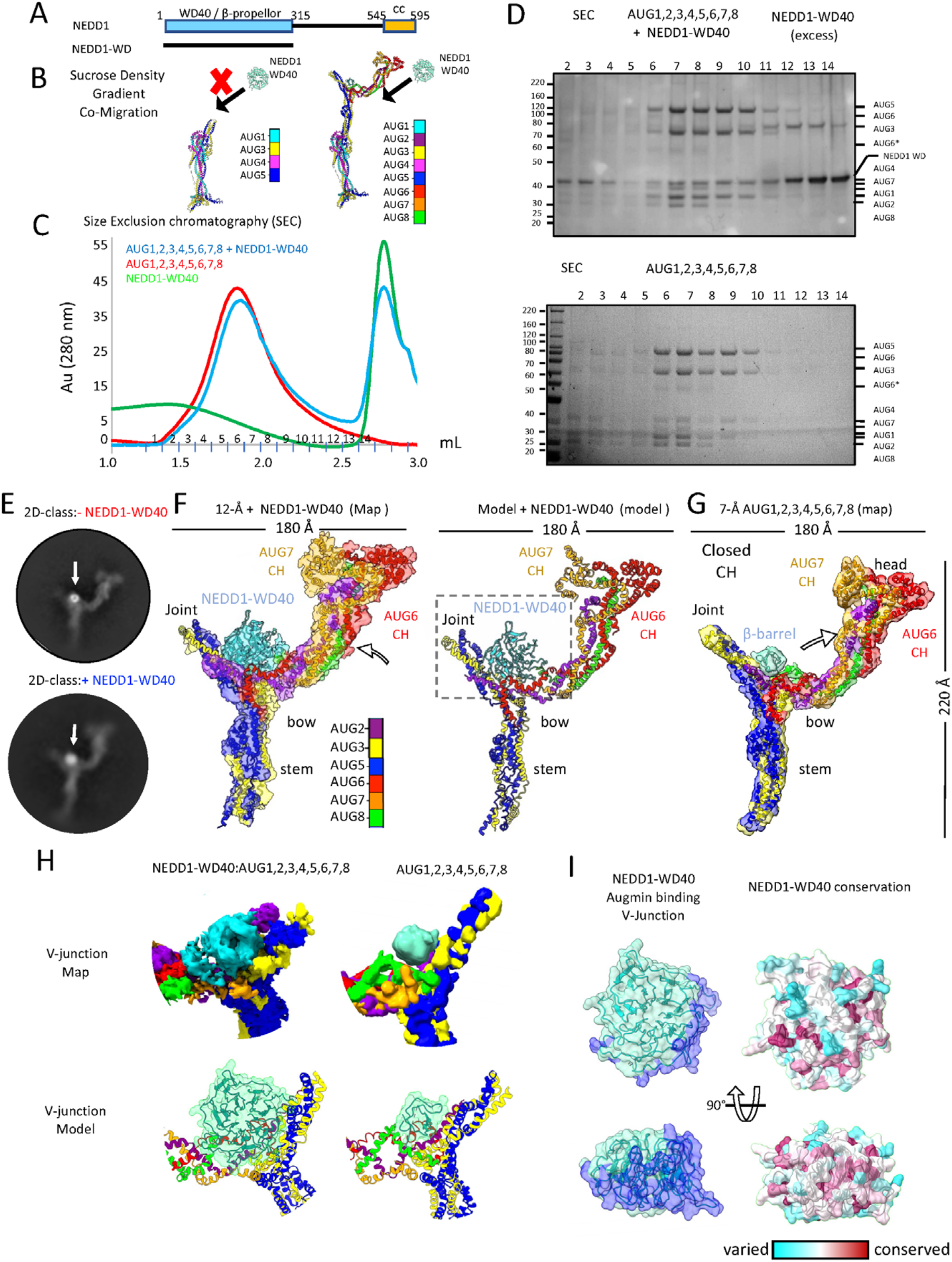
Biochemical reconstitution and structure NEDD1 WD40 β-propeller-Augmin complex. A) NEDD1 domain organization. N-terminal WD40 β-propeller domain. C-terminal helical domain (Note: Only WD40 β-propeller domain successfully purified). B) Complex formation analysis: AUG1,2,3,4,5,6,7,8 binds NEDD1-WD40 β-propellor and while AUG1,3,4,5 shows no binding (See **Figure S12C-H**). C) SEC analysis of complex formation. Red: AUG1,2,3,4,5,6,7,8 alone. Blue: AUG1,2,3,4,5,6,7,8 with NEDD1-WD40 β-propellor. Green: NEDD1-WD40 alone. D) Biochemical validation. SDS-PAGE showing co-migration of NEDD1-WD40 with AUG1,2,3,4,5,6,7,8 and comparison with AUG1,2,3,4,5,6,7,8 alone. E) Cryo-EM 2D-class average comparison NEDD1-WD40 β-propellor binding to Augmin. Top: AUG1,2,3,4,5,6,7,8 V-junction-stem with β-barrel density. Bottom: AUG1,2,3,4,5,6,7,8-NEDD1-WD40 β-propellor complex. F) Structural characterization of the complex. Left: 12 Å Cryo-EM segmented reconstruction with fitted models. Right: Ribbon representation of complex. G) 7.3-Å model showing β-barrel density bound to V-junction stem. Note the marked conformational change. H) Comparative analysis of V-junction. Top: Segmented maps of complex vs. AUG1,2,3,4,5,6,7,8 alone. Bottom: Atomic models of NEDD1-WD40 and β-barrel regions. I) NEDD1-WD40-β propellor interface analysis. Right: Surface views of Augmin interaction site. Left: Conservation analysis across species.

### Cryo-EM structure of Augmin-NEDD1-WD40 β-propellor reveals its binding onto the V-junction

To determine the NEDD1-WD40-β-propellor binding site on Augmin, we collected cryo-EM data for complexes of AUG1,2,3,4,5,6,7,8-NEDD1-WD40-β-propellor. Initial 2D-class averages show a 50% bigger mass on top of the V-junction where we the unknown barrel shaped density binds Augmin. These 2D classes also shows increased proportions of Augmin dimers and no 1.5 Augmin classes. We then determined a 12-Å cryo-EM structure revealing a larger mass, with circular shape and central hole, attached to the base of the V-junction (Figure 6E-G). This density is flipped by 90° and has a wider and circular “donut” like shape. Although this map is of lower resolution, we observe conformational change in the V-junction region, tilting upwards, upon placement and morphing of our previously determined Augmin model. We were able to place the AlphaFold predicted NEDD1-WD40 β-propellor model into the new density on top of the V-junction, revealing that the NEDD1 β-propellor binds via its narrow side interface orientation on top of the Augmin V-junction (**Figure 6F, H, I**). We plotted the conservation of plant and animal NEDD1 orthologs on the fitted NEDD1 WD40-β-propellor Alpha Fold model (**Figure 6I**). The conservation plot shows that the most conserved surface residues in NEDD1-WD40 β-propellor overlaps extensively on interface between the fitted NEDD1 Augmin V-junction model (**Figure 6I**). Furthermore, our low resolution cryo-EM structure and resulting model for the NEDD1-β-propellor-Augmin V-junction is fully consistent with our biochemical reconstitution studies (**Figure 6B-D**). The proximity of the NEDD1-WD40 β-propellor binding site at the base of the V-junction to AUG6,7 MT binding CH-domain dimer at the tip of the V-junction suggests that NEDD1-WD40 β-propellor MT binding may stabilize the conformation of Augmin’s V-junction to the MT lattice. We propose that the NEDD1-WD40 β-propellor binds both Augmin and MTs coupled with AUG6,7 CH-domain dimer MT binding may lead to extensive contact with multiple MT protofilaments (**model described below**; **Figure 7A**).

**Figure 7:**
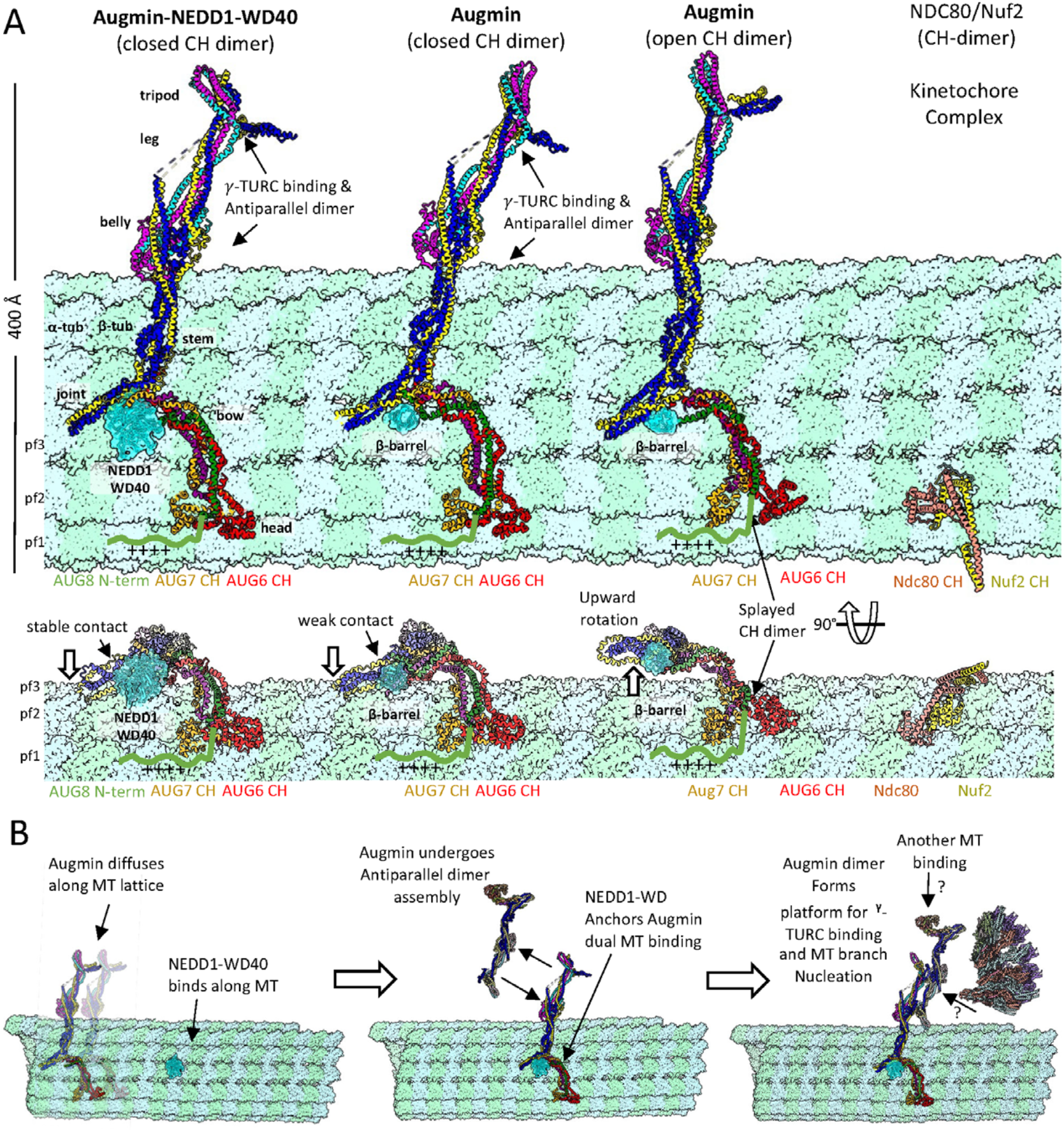
Mechanistic model of Augmin-NEDD1-MT interactions and MT branch nucleation. A) MT lattice binding modes. Left: NEDD1-WD40 complex with closed AUG6,7 CH-domains. Center-left: β-barrel density impact on MT binding. Center-right: Open CH-domain configuration. Right: NDC80/Nuf2 CH-domain reference structure views shown in a side and a 90° rotated orientations. B) Proposed mechanism for Augmin function. Left: Transition from diffuse to NEDD1-WD40-bound state. Middle: Formation of anchored complex and dimerization. Right: γ-TuRC recruitment and MT branch nucleation.

## Discussion

### Reconstitution and cryo-EM structures of plant Augmin provide new insights into its architecture and function

We have biochemically reconstituted and determined single particle cryo-EM structures of *A. thaliana* hetero-octameric (AUG1,2,3,4,5,6,7,8) and hetero-tetrameric (AUG1,3,4,5) Augmin assemblies (**Figure 1–2; video S1**). Our ability to reconstitute the multiple types of assemblies, apply single particle recentering, flexibility and heterogeneous refinement strategies allowed us to achieve medium to high resolution for the majority of the Augmin structure, not achieved in past studies^29–31^. Due to the improved resolution in these Augmin structures, we were able to produce a composite cryo-EM structure from our different maps of Augmin that allowed us to produce a *de novo* model of the plant Augmin hetero-octamer (**video S1**). This plant Augmin hetero-octamer model is supported by extensive XLMS studies that resolved a detailed interactions among subunits (**Figure 5)**. Our Plant Augmin model differs extensively from the metazoan AlphaFold or ColabFold derived models placed into the lower resolution cryo-EM structures (**Figure S14**)^29–31^. We also resolved two states, open and closed, for the AUG6,7 CH-domain dimer in the head of the Augmin V-junction region associated with changes in the V-junction arm (**Figure 2; video S1**).

### Biochemical and structural studies reveal roles of Augmin regions in dimerization and NEDD1 binding

Our biochemical and structural studies uncovered Augmin’s propensity for dimerization (**Figure 4**). Biochemical reconstitution of NEDD1 WD40 β-propellor with Augmin reveal its binding site on the Augmin V-junction and its ability to enhances Augmin dimerization. Our cryo-EM data reveals that Augmin particles form anti-parallel dimers mediated via two binding sites on the extended section in belly and tripod regions. We show the two dimerization interfaces are highly conserved across Augmin orthologs in plants and animals (**Figure S9; Figure 3E**).

Our cryo-EM structure of the Augmin-NEDD1-WD40 β-propellor reveal the NEDD1 binding site resides along the AUG2,6,7,8 on top of the V-junction, but likely induces conformational changes in the Augmin extended region promoting dimerization. These structures and biochemical studies a provide a new view of the long-distance cooperativity within Augmin between its NEDD1/MT binding sites at V-junction and its γ-TuRC binding site, likely residing along its extended region. The V-junction binding to NEDD1 WD40 β-propellor and MTs likely promotes the extended region dimerization, generating a platform for γ-TuRC binding (model described below).

### A two-site model for Augmin V-junction and NEDD1 binding along MT lattices

Our Augmin structural and biochemical studies allow us to develop an Augmin MT binding site relation to NEDD1-WD40 β-propellor and its structural organization as shown in **Figure 7**. Recent total internal reflection fluorescence reconstitution studies of human Augmin and 1-TuRC in MT branch nucleation by Zhang et al dissected the relationship of between NEDD1 and Augmin in binding the MT lattice^20^. The initial binding of Augmin to MTs is dynamic and diffusive. However, Augmin signal increases by two to three-fold leading to static Augmin binding along MTs, suggesting Augmin oligomerization may impact high affinity MT binding^20^. After this transition, Augmin(s) efficiently recruit γ-TuRC to activate branch MT nucleation. Augmin binding to MTs is stabilized by NEDD1-WD40 β-propellor binding to MT lattices. Our Augmin structures provide a potential explanation for this process. We compared our Augmin structures with conformation of AUG6,7 CH domain dimer in open and closed states and Augmin-NEDD1 WD40 β-propellor state, all modeled from our structures, by docking them onto the MT lattice, as shown in **Figure 7A**. We overlaid the Augmin via the dual AUG6,7 Ch-domain onto the NDC80/Nuf2 kinetochore dual CH domain dimer in their MT bound states (**Figure 7A, right**)^27^. In our docking, the Augmin V-junction lays along MTs bridging across multiple protofilaments (**Figure 7A**). Due to the conformational changes in the V-junction of each of the states of Augmin (Video S1), the footprint of each of the V-junction regions of closed, open and NEDD1-WD40 β-propellor bound states are compared. Comparison of the size of MT footprint of each of the Augmin v-junction in of the three states on interaction with the MT binding sites suggests that Augmin AUG6,7 CH-domains dimer in the closed state likely binds with a higher affinity along MTs compared to the open state (**Figure 7A, middle and right panels**). The binding of NEDD1-WD40 β-propellor to the V-junction forms a second crucial MT binding site that likely stabilizes Augmin V-junction interaction with multiple MT protofilaments (**Figure 7A, left panel**) With this stabilized dual binding interface, NEDD1-WD40 β-propellor bound Augmin becomes more tightly bound. We believe this organization may explain the “force bearing” properties of Augmin in stabilizing MT branch nucleation sites. This binding arrangement rationalizes the V-junction shape of the interface which leads the 30-nm Augmin extended region to become propped above the MT surface (**Figure 7A, video S1**).

### Augmin dimerization may anchoring of the γ-TuRC on MTs and activate branch nucleation

Our MT binding model presents several testable hypotheses regarding the roles of Augmin, NEDD1, and γ-TuRC domains in assembling the MT branch junction machinery (**Figure 7B**). Augmin extended regions likely dimerize, as visualized in our structure (**Figure 4**), creating larger interfaces to bind and activate γ-TuRC complexes, anchoring them more efficiently along MTs to promote MT branch nucleation. NEDD1’s helical coiled-coil C-terminal domain may facilitate Augmin dimerization by forming tetrameric oligomers which may γ-TuRC binding by interacting with Mozart and the GCP6 N-terminal region, as described in a recent structural study^33^. Augmin oligomeric assemblies, promoted by NEDD1, likely form a key platform for γ-TuRC binding. Our newly resolved Plant Augmin structures highlight the complexity of the multi-helical interactions stabilizing the Augmin hetero-octamer. Many of these interactions are highly conserved across plants, animals, and insects, contributing to the well-conserved Augmin assembly overall shape. The multi-helical coiled-coil interactions of AUG1–8 make Augmins flexible, potentially allowing communication between their V-junction and extended regions in response to biochemical of mitotic phosphorylation cues. However, Augmin oligomerization appears to serve an as-yet unknown but critical function in forming the platform for γ-TuRC binding and MT branch nucleation (**Figure 7B**). This behavior is reminiscent of other regulatory complexes, such as the crosslinker PRC1 or COPII coatomer proteins, which also undergo head-to-tail dimerization^34,35^.

## Conclusion

We have determined cryo-EM structures leading to a full *de novo* model for plant Augmin complex coiled-coil assembly that is verified using XLMS. We found that Augmin forms antiparallel dimers through conserved interfaces in its extended region, and that NEDD1 WD40 β-propellor domains directly bind the Augmin V-junction, close to its AUG6,7,8 MT binding site. Altogether we present a model in which NEDD1 promotes Augmin’s MT binding and oligomerization through downstream conformational effects that result in the recruitment of the γ-TuRC complex in MT branch formation.

## Acknowledgement

We would like to thank Dr. Ian Humphreys and Dr. David Baker (University of Washington, Seattle) for initial help with AlphaFold2 predictions of hetero-tetrameric and hetero-octameric Augmin assemblies. We like to thank Dr Hernando Sosa (Albert Einstein College of Medicine) for providing a PDB vectorial comparison script. We like to thank Dr Richard McKenney, Dr Gant Luxton (Molecular Cellular Biology UC-Davis) for the critical reading of this manuscript. Cryo-EM data were collected at the UC-Davis cryo-EM facility with support by the Molecular Cellular Biology department and College of Biological sciences. We thank Dr Camille Scott at the UC-Davis High Performance Computing Facility for computational support for cryo-EM structure determination. BL and JAB thank the College of biological Sciences and Departments of Plant Biology and Molecular Cellular Biology at UC-Davis for the generous support of this inter-departmental collaboration. JAB acknowledges grant support from the National Institutes of Health (GM110283). BL acknowledges grant support from the National Science foundation (NSF/BSF-2416267 and MCB-2148207). SDF acknowledges support from the National institutes of health (1DP2GM140926), National Science foundation (MCB-2045844) Camille Dreyfus foundation and the Sloan Foundation. Cryo-EM maps and models presented here will be available in the Electron microscopy database (EMDB) with the EMBD-IDs: EMD-49225, EMD-49224, EMD-49227, EMD-49182, EMD-49183, EMD-49230 and corresponding models are available at the Protein data bank (PDB) with the PDB-ID 9NBB,9NBA,9NBD,9NA8,9NA9 and 9NBI

## Author contribution

MA purified Augmin assemblies, carried out all biochemical studies, prepared XLMS samples, determined and refined all structures, built and refined all models, supervised the project and co-wrote and co-revised manuscript. AT purified Augmin assemblies, prepared cryo-EM grids, collected cryo-EM data, and determined initial structures. YRJL prepared bacterial polycistronic constructs for Augmin reconstitution and expression. YT prepared and carried out XLMS experiment. FG assisted cryo-EM data collection and cryo-EM grid preparation. SDF provided grant support, supervised and advised XLMS studies. BL provided grant support, supervised and advised Augmin expression construct preparation and data interpretation. JAB conceived the project, provided grant support, supervised and advised all co-authors, carried out biochemical experiments, prepared figures, wrote and revised manuscript.

## Methods

### Cloning, Protein expression and purification

*A. thaliana* (At) Augmin AUG1,2,3,4,5,6,7,8 subunits and other plant orthologs were aligned with metazoan and insect counterparts, revealing high conservation The C-terminal region (residues 338-644) of AUG8 contains a highly conserved α-helical domain, while its divergent, disordered N-terminal domain containing the MT binding region was excluded.

AUG2,6,7,8 sequences contained multiple plant specific rare Arg codons absent in bacteria, necessitating codon optimization. Each AUG1,2,3,4,5,6,7,8 subunit ORF was cloned into pET3a vectors and assembled into polycistronic co-expression constructs (**Figure S1A, S1F**) with C-terminal 8Xhis tags on AUG5 (His) and AUG6-StrepII tag (Strep) (**Figure S1A**). AUG1,3,4,5 subunits were assembled into a polycistronic vector with AUG5 C-terminal GFP and His tags (**Figure S1F**). The *At* NEDD1-WD40 β-propellor (residues 1-315) domain was cloned into pFastBac with strepII tag.

Augmin Polycistronic constructs were transformed into SoluBL21 (AMSBio) cells on ampicillin plates. Hetero-octameric (AUG,1,2,3,4,5,6,7,8) Augmin was overexpressed from overnight cultures diluted 1:200 into 2xYT media with 200 µg/mL ampicillin. Cells were induced with 0.5 mM IPTG at OD^600^ 0.6, grown at 18° C for 12-14 h, and harvested by centrifugation at 3,000 rpm for 25 min. Pellets were resuspended in lysis buffer (50 mM HEPES pH 7.5, 200 mM KCl, 1 mM MgCl^2^, 1 mM EGTA, 12 mM ß-mercaptoethanol, 10 % (v/v) glycerol) with protease inhibitors (5 µg/mL leupeptin, pepstatin aprotinin, 0.1 mM PMSF, 5 µg/mL benzamidine, EDTA-free mini tablets (Sigma Aldrich). DNaseI was added and cells lysed by microfluidization. Clarified lysates (18,000 rpm, 20 min, 4° C) were loaded onto recycled Ni-IDA columns (Macherey Nagel). Augmin was eluted with 200 mM Imidazole after washing. Assembly was verified by SDS-PAGE. Eluates were loaded onto Hi5-QFF columns (cytiva), and flow-through was further purified on Superose 6 16/60 columns in 50 mM HEPES pH 7.5, 200 mM KCl, 1 mM EGTA, 1 mM MgCL^2^, 10% (v/v) glycerol. The hetero-tetrameric (AUG1,3,4,5) Augmin was purified similarly but eluted from Q-FF with ~400 mM KCl and gel filtered on Superdex 200 16/60 (cytiva) in 50 mM HEPES pH 7.5, 300 mM KCl, 1 mM MgCl2, 1 mM EGTA, 5% (v/v) glycerol. Aliquots (3 mg/mL) supplemented with 15% (v/v) glycerol were snap-frozen.

*At* NEDD1-WD40 β-propellor expressed in Hi5 cells (1:20 virus inoculum) 27 °C for 60-70 h. Cells (≥90% viability) were lysed in 50 mM HEPES pH 7.5, 250 mM KCl, 1 mM MgCl2, 1 mM EGTA, 5% (v/v) glycerol with protease inhibitors. NEDD1-WD4040 was purified on regenerated Strep-Tactin columns (IBA) with 100 mM *D*-biotin elution and gel filtered on Superdex 200 16/60 (**Figure S12**) in lysis buffer. Aliquots (1 mg/mL) with 20% (v/v) glycerol were snap-frozen.

Purified full-length Augmin subunits were analyzed by LC-MS/MS (Taplin Facility). Gel bands were excised from 12% SDS-PAGE, digested, and identified peptides manually mapped.

### Mass Photometry and Multi-angle light scattering experiments

Mass Photometry experiments were performed using a Refyen 1.0 mass photometer. Data were analyzed in Discover MP software using mass calibration provided by Refyen. Samples (**Figures S1D, S1H, S12G, and S12H**) were crosslinked with a ~50-100-fold molar excess of glutaraldehyde, then diluted to 5-10 ng/mL in 12 µL 20 mM HEPES pH 7.0, 100 mM KCl. Non-crosslinked samples were analyzed without dilution in 50 mM HEPES pH 7.5, 150-300 mM KCl, 1 mM MgCl2, 1 mM EGTA.

SEC-MALS was performed using an HPLC system equipped with a Superdex 200 Increase 10/300 column inline with a Wyatt Mini-DAWN TREOS multi-angle light scattering detector and Optilab reflective index detector (Wyatt Technologies). AUG1,2,3,4,5,6,7,8 (200 µL, 1 mg/mL; **Figure S1B**) or AUG1,3,4,5 (200 µL, 1 mg/mL; **Figure S1G**) were injected and eluted in 50 mM HEPES pH 7.5, 200 mM KCl, 1 mM MgCl2, 1 mM EGTA. Molecular masses were determine using ASTRA software (Wyatt Technologies).

### Augmin/Nedd1-WD40 β-propellor binding experiments

Reconstitutions of hetero-octameric (AUG1,2,3,4,5,6,7,8) with NEDD1 was evaluated was by size exclusion chromatography on a Superose 6 5/150 column (cytiva) in 50 mM HEPES pH 7.5, 150 mM KCl, 1 mM MGCl2, 1 mM EGTA, 5% (v/v) glycerol with 100 µL injections. AUG1,2,3,4,5,6,7,8 at 1 mg/mL was incubated with and without a 10-fold molar excess of WD40-NEDD1. Sucrose density gradient experiments were performed with AUG1,2,3,4,5,6,7,8 (~1 mg/mL) or AUG1,3,4,5 (~2 mg/mL) in the presence or absence of a 5-fold molar excess of NEDD1-WD40 on 10-40 % sucrose gradients in 50 mM HEPES pH 7.5, 150 mM KCl, 1 mM EGTA, 1 mM MgCl2 and fractions were evaluated using SDS PAGE (**Figure S12C-H**)

### Cryo-EM sample preparation and data collection

The hetero-octameric (AUG1,2,3,4,5,6,7,8) and Hetero-tetrameric (AUG1,3,4,5) Augmin samples were purified by size exclusion chromatography on Superdex 200 10/300 columns (cytiva), concentrated to 1 mg/mL, and crosslinked with 200 nM BS3 (ThermoFisher) on ice for 2 h, followed by quenching with 10 µM Tris-HCl pH 8. AUG1,2,3,4,5,6,7,8-NEDD1-WD40-β-propellor complexes were purified on a Superdex 200 16/60 column, concentrated to 3 mg/mL, crosslinked with 1 μM BS3 for 1 h, and quenched with 1 mM Tris-HCl pH 7.0. Buffers are described in the Supplementary Information.

Cryo-EM grids were prepared using a Leica EM GP2 with sensor blotting at 20 °C and 95% humidity, 5-10 s pre-blot, 5-8 s blot time, and 1.5-1.8 mm extra push. R 1.2/1.3 300 mesh grids (Quantifoil) were used (Cu for AUG1,2,3,4,5,6,7,8 and AUG1,3,4,5; Au for AUG1,2,3,4,5,6,7,8-NEDD1-WD40). Some AUG1,2,3,4,5,6,7,8 grids included 0.001% NP-40.

Grids were screened on a Glacios microscope (Thermo Fisher) with a K3 direct electron detector (Gatan) at 200 kV using SerialEM^36^ at 11,000x and 45,000x magnifications using low dose conditions. High-resolution data were collected at 45,000x (0.44 Å/pixel) in Super-resolution mode) with SerialEM^36^ and beam-image shift^37^, at −0.6 to −1.8 µm defocus and 60 e-/Å^2^ total dose over 75 frames

### Cryo-EM Single Particle Image processing

#### full Augmin hetero-octamer (AUG1,2,3,4,5,6,7,8) structure

(***Figure S2***,*black arrows):* 5717 movies of hetero-octameric (AUG1,2,3,4,5,6,7,8) Augmin were motion-corrected (MotionCor2^38^, 2x binned) and CTF-estimated (CTFfind^39^) in RELION^40^. Particles (20 million) were picked by LoG-based template-free picking (50-450 Å diameter) and extracted (200 pixels, 3.52 Å/pixel). After 2D classification in cryoSPARC^41^, 714,446 particles were re-extracted in RELION^40^ (300 pixels, 1.76 Å/pixel). Further 2D- and 3D-classification. Using *ab initio* model generation followed by refinement yielded a 9.6 Å map from 180,160 particles, 3DFlex^42^ refinement in cryoSPARC^41^ followed by 2D-classification and homogeneous refinement produced a 9.1 Å map based on Fourier shell correlation (FSC). Local resolution was calculated in PHENIX^43^.

#### The Augmin V-junction-stem structure (closed state)

(***Figure S3***): Particles were re-centered on the V-junction and re-extracted (160 pixels, 1.76 Å/pixel). 2D-classification selected 440,976 particles, which were 3D-refined, CTF-refined, and polished^44^ in RELION^40^. Duplicate-removed particles (223,355) were re-extracted (200 pixels, 1.76 Å/pixel), and *ab initio* model was generated and refined in cryoSPARC^41^. Heterogeneous refinement with three *ab initio* models yielded 107,694 particles in the closed state. Further 2D-classification and 3DFlex^42^ refinement, followed by re-extraction, 2D- and 3D-refinement, produced a 7.3 Å closed V-junction map from 75,828 particles. A second dataset of hetero-octameric (AUG1,2,3,4,5,6,7,8) Augmin 11,256 movies was processed similarly, and particles from both datasets (200,853) were merged in cryoSPARC^41^. 2D, heterogeneous, non-uniform^45^, and local refinement, combined with 3D variability analysis^46^ and re-extraction (256 pixels, 1.76 Å/pixel), yielded a final 7.3 Å map from 18,243 particles (3DFlex^42^ refinement, DeepEMhancer^47^ sharpening).

#### The Augmin V-junction-stem (open state)

(***Figure S3***, *grey arrows):* 94,659 particles from heterogeneous refinement of the closed state were 3D refined, 2D classified, and re-extracted (200 pixels, 1.76 Å/pixel) to select 67,132 particles. Homogeneous, non-uniform^45^, and 3DFlex^42^ refinement, followed by 3D variability analysis^46^ and individual frame reconstruction, selected 37,048 particles, 3DFlex^42^ refinement, CTF refinement, and non-uniform refinement produced a 12 Å open V-junction map, after sharpened using DeepEMhancer^47^.

#### Antiparallel Augmin dimer

(***Figure S2***, *blue arrows):* 6,578 particles from 2D classification were used to generate an initial dimer map in cryoSPARC^41^, which was refined in RELION^40^. Particles were re-centered on the AUG1,3,4,5 overlap region and re-extracted (384 pixels, 0.88 Å/pixel). Homogeneous and non-uniform refinement with C2 symmetry, followed by 3DFlex^42^ refinement, yielded an 8.1 Å map (B-factor sharpening).

#### Augmin 1.5-mer structure

(***Figure S2***, *gray arrows):* 81,112 particles showing a second leg were separated by heterogeneous refinement, re-extracted (350 pixels, 1.76 Å/pixel), and 2D classified to select 11,903 particles. Ab initio reconstruction followed by homogeneous and non-uniform refinement produced a 12.4 Å 1.5-mer map.

#### The hetero-tetrameric (AUG1,3,4,5) Augmin extended region structure

(***Figure S4***): 10,830 movies were motion-corrected (MotionCor^38^, 2x binned), and CTF-estimated (GCTF^48^). 15.5 million particles were picked by reference-free LoG picking (80-350 Å diameter) and extracted (100 pixels, 3.52 Å/pixel). 2D classification in cryoSPARC^41^ selected 638,316 particles, which were re-extracted (200 pixels, 1.76 Å/pixel) and 3D refined in RELION^40^, followed by CTF refinement and polishing of 387,892 particles. Iterative 3D refinement, 2D/3D classification, and re-extraction (500 pixel, 0.88 Å/pixel) yielded a 4.1 Å map from 247,267 particles. 3D classification selected 144,287 particles, which were re-extracted (512 pixels, 0.88 Å/pixel) and refined in RELION^40^ and cryoSPARC^41^ to produce a 3.9 Å map. Re-extraction (400 pixels, 0.88 Å/pixel), 2D classification, and iterative refinement in cryoSPARC^41^ yielded a 3.5 Å map from 72,002 particles (3D variability analysis^46^, 3DFlex^42^ refinement, DeepEMhancer^47^ sharpening). Focused local refinement of the tripod with a tight mask produced a 5.9 Å map.

#### The Augmin V-junction-NEDD1-WD40-β-propellor

(***Figure S13***): 4,583 movies from two datasets were motion-corrected (motioncorr2, 2x binned), CTF-estimated (CTFFind3), and blob-picked (50-600 Å diameter) in cryoSPARC^41^. 2D classification of 10 million initial particles (200 pixels, 3.52 Å/pixel) selected 15,844 particles used for template picking, yielding 510,466 particles. 2D classification selected 39,064 particles, which were used to generate an ab initio model and 3D refined. Particles were re-extracted (250 pixels, 1.76 Å/pixel), and 2D classification selected 27,025 particles. 3D refinement and 3DFlex^42^ refinement, followed by re-extraction (200 pixels, 1.76 Å/pixel) and homogeneous, non-uniform, and local refinement, produced a 12 Å map from 26,681 particles (DeepEMhancer^47^ sharpening).

### Model building and refinement

Model interpretation was limited to lower than reported resolutions due to resolution estimate inflation likely caused by map flexibility and low signal-to-noise of elongated particles, a trend observed in all published Augmin reconstructions^29–31^. Interpretation was based on visual map features rather than nominal resolutions. The extended region (AUG1,3,4,5) maps ranging from 3.5-4 Å were interpreted to 4 Å. The AUG1,3,4,5 tripod density and closed V-junction CH-domain maps at 5.9 Å were interpreted to 7.3 Å for the full Augmin model. Open CH-domain, 1.5-mer, and dimer maps were interpreted at 12 Å, 15 Å, and 10 Å, respectively.

Initial models were built by rigid body fitting AlphaFold3 predictions (**Figure S8**) for AUG1,3,4,5 and AUG1,2,3,4,5,6,7,8 into density using UCSF ChimeraX^49^ and Coot^50^. The AUG3,5 fold-back zone and AUG2,6,7,8 regions were built into the 7.3 Å V-junction map by flexible fitting and manual adjustment based on AlphaFold3 secondary structure predictions. The open CH-domain conformation (**Figure S6C**) was modeled by rigid body fitting into distinct CH-domain-like densities.

The 3.7 Å AUG1,3,4,5 map was used for *de novo* modeling of the interacting helical regions in the belly and legs, starting from the AlphaFold prediction. AUG3,5 N-terminal and C-terminal helical bundles and AUG1,4 C-termini were built into the 6 Å map using secondary structure element length, connectivity, and interactions as guides.

A full *de novo* AUG1,2,3,4,5,6,7,8 model was assembled by merging the above V-junction-stem and extended region models and fitting into the 8.1 Å consensus map using a 20 Å, 4-helix bundle in the central AUG3,5 (**Figure S5A-B**). The dimer model was built by rigid body fitting two copies of AUG1,2,3,4,5,6,7,8 into the dimer map. The AUG1,2,3,4,5,6,7,8-NEDD1-WD40 model (**Figure S14B**) was built by fitting the closed V-junction model and placing NEDD1-WD40 into the junction density. All models were subjected to real-space refinement in PHENIX^43^ (**Table 1**).

**Table 1:** Cryo-EM data collection, processing and model building.

| Parameters | Augmin<br>V-junc<br>(closed) | Augmin<br>V-junc<br>(open) | Augmin<br>full | Augmin<br>dimer | Augmin<br>extended<br>body | Augmin<br>extended<br>(Tripod) | Augmin<br>V-junc/<br>NEDD1-WD40 |
| --- | --- | --- | --- | --- | --- | --- | --- |
| Magnification | 45000x |  |  |  |  |  |  |
| Voltage (kV) | 200 |  |  |  |  |  |  |
| Electron exposure (e Å<sup>-2</sup>) | 60 |  |  |  |  |  |  |
| Defocus range (μm) | -0.6 to -1.8 |  |  |  |  |  |  |
| Pixel size (Å) | 0.44 | 0.44 | 0.44 | 0.44 | 0.44 | 0.44 | 0.44 |
| Symmetry imposed | *C1* | *C1* | *C1* | *C2* | *C1* | *C1* | *C1* |
| No. of final particle images | 18243 | 37048 | 123579 | 6578 | 72005 | 72005 | 26681 |
| Map resolution FSC threshold (Å) | 5.9 | 8.6 | 9 | 9.5 | 3.5 | 5.9 | 11.8 |
| Initial model used | AlphaFold multimer and AlphaFold2 prediction |  |  |  |  |  |  |
| Model resolution FSC threshold (Å) | 0.143 |  |  |  |  |  |  |
| PDB ID | 9NBB | 9NBA | - | 9NBD | 9NA8 | 9NA9 | 9NBI |
| EMDB code | EMD-49225 | EMD-49224 | - | EMD-49227 | EMD-49182 | EMD-49183 | EMD-49230 |
| Model resolution (Å) | 8 | 12 | - | 12 | 5 | 7 | 15 |
| Model composition |  |  |  |  |  |  |  |
| Chains | 6 | 6 | 8 | 8 | 4 | 4 | 7 |
| Non-hydrogen atoms | 11941 | 10976 | - | 22570 | 7364 | 4172 | 14095 |
| Protein | 1500 | 1375 | - | 2842 | 935 | 513 | 1785 |
| Nucleic acid | 0 | 0 | - | 0 | 0 | 0 | 0 |
| Ligand | 0 | 0 | - | 0 | 0 | 0 | 0 |
| R.M.S. deviations |  |  |  |  |  |  |  |
| Bonds (Å) | 0.003 | 0.003 | - | 0.003 | 0.003 | 0.003 | 0.003 |
| Angles (°) | 0.654 | 0.745 | - | 0.771 | 0.783 | 0.897 | 0.698 |
| MolProbity score | 2.27 | 2.58 | - | 2.53 | 2.08 | 1.99 | 2.43 |
| Clash score | 24.58 | 36.07 | - | 34.69 | 14.51 | 24.08 | 30.08 |
| Poor rotamers (%) | 0 | 0 | - | 0 | 0 | 0.22 | 0 |
| Ramachandran plot |  |  |  |  |  |  |  |
| Favored (%) | 94.35 | 90.17 | - | 91.53 | 95.67 | 97.41 | 92.66 |
| Allowed (%) | 5.65 | 9.76 | - | 8.40 | 4.33 | 2.59 | 7.34 |
| Disallowed (%) | 0 | 0.07 | - | 0.07 | 0 | 0 | 0 |

### Crosslinking mass spectrometry (XLMS) of Augmin assemblies

AUG1,2,3,4,5,6,7,8 and AUG1,3,4,5 samples were crosslinked with 0.5-2 mM BS3 at 4°C overnight, denatured (8 M urea), reduced (5 mM DTT), alkylated (15 mM IAA), quenched with DTT, digested with LysC and trypsin, desalted (Sep-Pak C18), and vacuum-dried. Desalted peptides were fractionated via Superdex 30 10/300 GL gel filtration, then dried and stored at −80°C. AUG1,3,4,5 fractions were analyzed using an UltiMate3000 UHPLC system coupled to a Q-Exactive HF-X Orbitrap. AUG1,2,3,4,5,6,7,8 fractions were analyzed with a Vanquish Neo UHPLC coupled to Orbitrap Ascend. Peptides were loaded onto PepMap 100 C18 column with Solvent A (0.1% FA in water) and solvent B (0.1% FA in ACN). AUG1,3,4,5 peptides were separated using PepMap RSLC C18 column with gradients from 5% to 90% Solvent B over 130 min; AUG1,2,3,4,5,6,7,8 peptides were separated with gradients from 10.4% to 76% Solvent B. For LC-MS/MS data acquisition, the Q-Exactive HF-X performed MS1 scans at 120,000 resolution (350-1500 m/z), AGC target of 3 × 10^6^, and 50 ms max IT. The top 10 precursors (z = 3-8) were isolated (1.4 m/z window) and fragmented using stepped NCE (30±6). MS2 scans were at 60,000 resolution (200-2000 m/z), AGC target of 8 × 10^3^, and 150 ms max IT. Dynamic exclusion was set to 45 s and in-source CID at 10 eV. For the Orbitrap Ascend, MS1 scans were at 240,000 resolution (380-2000 m/z), normalized AGC target of 150%, and 100 ms max IT The top 20 precursors (z = 4-7preferred than 3) were isolated (1.4 m/z window) and fragmented (NCE 30±6). MS2 scans were at 60,000 resolution (150-2000 m/z), normalized AGC target of 750%, and 250 ms max IT. Dynamic exclusion was set to 30 s and in-source CID at 10 eV. RAW files were processed with the xiSEARCH pipeline and searched using xiSEARCH 1.8.6 (MS1 tolerance: 6 ppm; MS2: 10 ppm; up to 4 miscleavages). Modifications included fixed Cys carbamidomethylation (+57.02 Da) and variable Met oxidation (+15.99 Da), with preferred crosslinks at Lys/N-term and lower priority at Ser, Thr, Tyr. FDR filtering (5%) was done using xiFDR 2.3.2. Figures generated in ChimeraX^48^ with XMAS^56^.

**Video S1: Summary of Cryo-EM structures, segmented maps and models of Plant Augmin** The video shows the hetero-octameric (AUG1,2,3,4,5,6,7,7,8) Augmin 10-Å cryo-EM structure, followed by the placement of the raw 7.3-Å V-junction stem region of AUG1,2,3,4,5,6,7,8 cryo-EM structure (blue) and the extended region of the Augmin (AUG1,3,4,5) hetero-tetramer 3.7-Å cryo-EM structure (red) into the 10-Å full Augmin. The segmented maps and fitted models are shown for 7.3-Å V-junction stem in the closed state, 3.7-Å cryo-EM structure of the extended region, 10-Å cryo-EM structure of the V-junction stem in the open state, and 12-Å cryo-EM structure of the NEDD1-WD40-β-propellor bound V-junction stem. The overlap 20-Å zone between the extended region and the V-junction stem maps are marked and allow composites to be generated to build *de novo* models. We next show in succession the full models of the full hetero-octameric (AUG,1,2,3,4,5,6,7,8) Augmin in the closed state, then open state, and the NEDD1-WD40-β-propellor bound state. Finally, the three full Augmin models are overlaid with the Augmin closed state in blue, the Augmin open state in green and Augmin-NEDD1-WD40-β-propellor bound state in red are all compared, to compare transitions of the V-junction.

