## Supplementary Figures S1-S14 for "Cryo-EM structures of the Plant Augmin reveal its intertwined coiled-coil assembly, antiparallel dimerization and NEDD1 binding mechanisms"

Figure S1

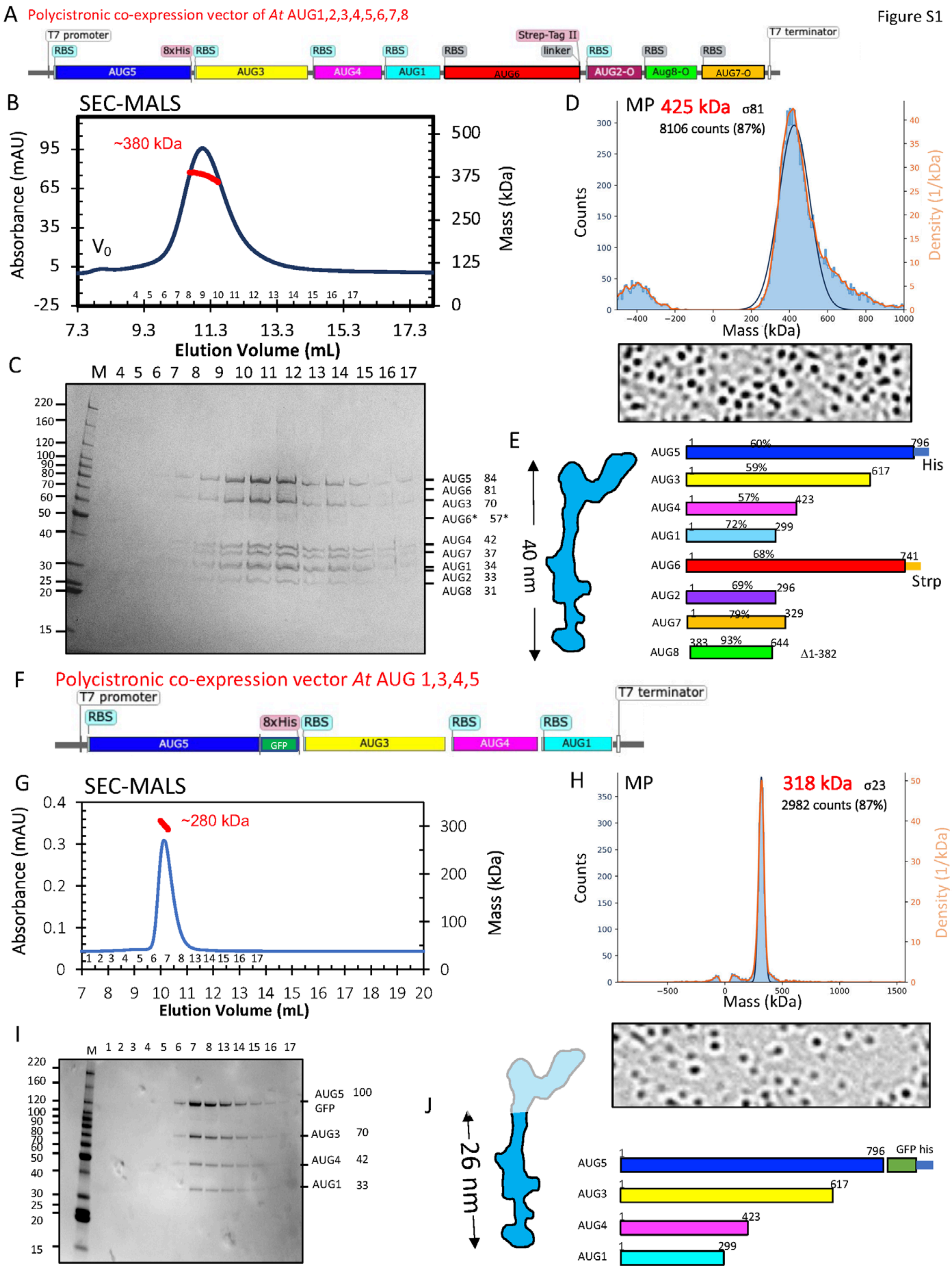

A Polycistronic co-expression vector of At AUG1,2,3,4,5,6,7,8
T7 promoter
RBS
8xHis
AUG5
AUG3
AUG4
AUG1
AUG6
Strep-Tag II
linker
AUG2-O
Aug8-O
AUG7-O
T7 terminator
B SEC-MALS
~380 kDa
V0
Absorbance (mAU)
Mass (kDa)
Elution Volume (mL)
C
AUG5 84
AUG6 81
AUG3 70
AUG6* 57*
AUG4 42
AUG7 37
AUG1 34
AUG2 33
AUG8 31
D MP 425 kDa σ81
8106 counts (87%)
Counts
Density (1/kDa)
Mass (kDa)
E
40 nm
60%
796
His
59%
617
57%
423
72%
299
68%
741
Strp
69%
296
79%
329
383
93%
644
Δ1-382
F Polycistronic co-expression vector At AUG 1,3,4,5
GFP
G SEC-MALS
~280 kDa
H MP 318 kDa σ23
2982 counts (87%)
I
AUG5 100
GFP
AUG3 70
AUG4 42
AUG1 33
J
26 nm
GFP his

**Figure S1**: **Biochemical reconstitution and characterization of *Arabidopsis thaliana* Augmin**

A) Organization of the AUG1,2,3,4,5,6,7,8 polycistronic vector for expressing assemblies in bacteria showing the T7 promoter, ribosomal binding site (RBS) and open reading frame (ORF) order for each of the eight subunits
B) Size exclusion chromatography (SEC) with multi-angle light scattering (SEC-MALS) for AUG1,2,3,4,5,6,7,8 hetero-octameric assemblies revealing their overall mass to be 380 kDa
C) SDS-PAGE of SEC-fractions revealing the eight AUG subunits of traces shown in B.
D) Top, Mass photometry measured for masses purified AUG1,2,3,4,5,6,7,8 fitted with Gaussian distribution revealing masses of 425 KDa. Bottom, example raw mass photometry images of AUG1,2,3,4,5,6,7,8 assemblies.
E) Left the overall organization of the 40 nm hetero-octameric full Augmin particle, Right, linear scheme for all *AT* AUG subunits in the assembly as shown in **Figure 1**. Percentage values above indicate coverage regions of polypeptide in mass spectrometry of purified assemblies.
F) Organization of the AUG1,3,4,5 polycistronic vector for expressing assemblies in bacteria showing the T7 promoter, ribosomal binding site (RBS) and open reading frame (ORF) order for each of the four subunits with AUG5-containing a C-terminal Green Fluorescent Protein (GFP) and His tag.
G) Size exclusion chromatography with multi-angle light scattering (SEC-MALS) for hetero-tetrameric (AUG1,3,4,5) Augmin assemblies revealing their overall mass to be 280 kDa
H) SDS-PAGE of SEC-fractions revealing the four AUG subunits of traces shown in G.
I) Top, Mass photometry measured for masses purified AUG1,3,4,5 assemblies fitted with Gaussian distribution revealing masses of 318 KDa. Bottom, example raw mass photometry images of AUG1,3,4,5 assemblies.
J) Left the overall organization of the 30 nm hetero-tetrameric Augmin particle, Right, linear scheme for all *AT* AUG subunits in the assembly as shown in **Figure 1**.

Cryo-EM processing of the Aug1,2,3,4,5,6,7,8

Figure S2

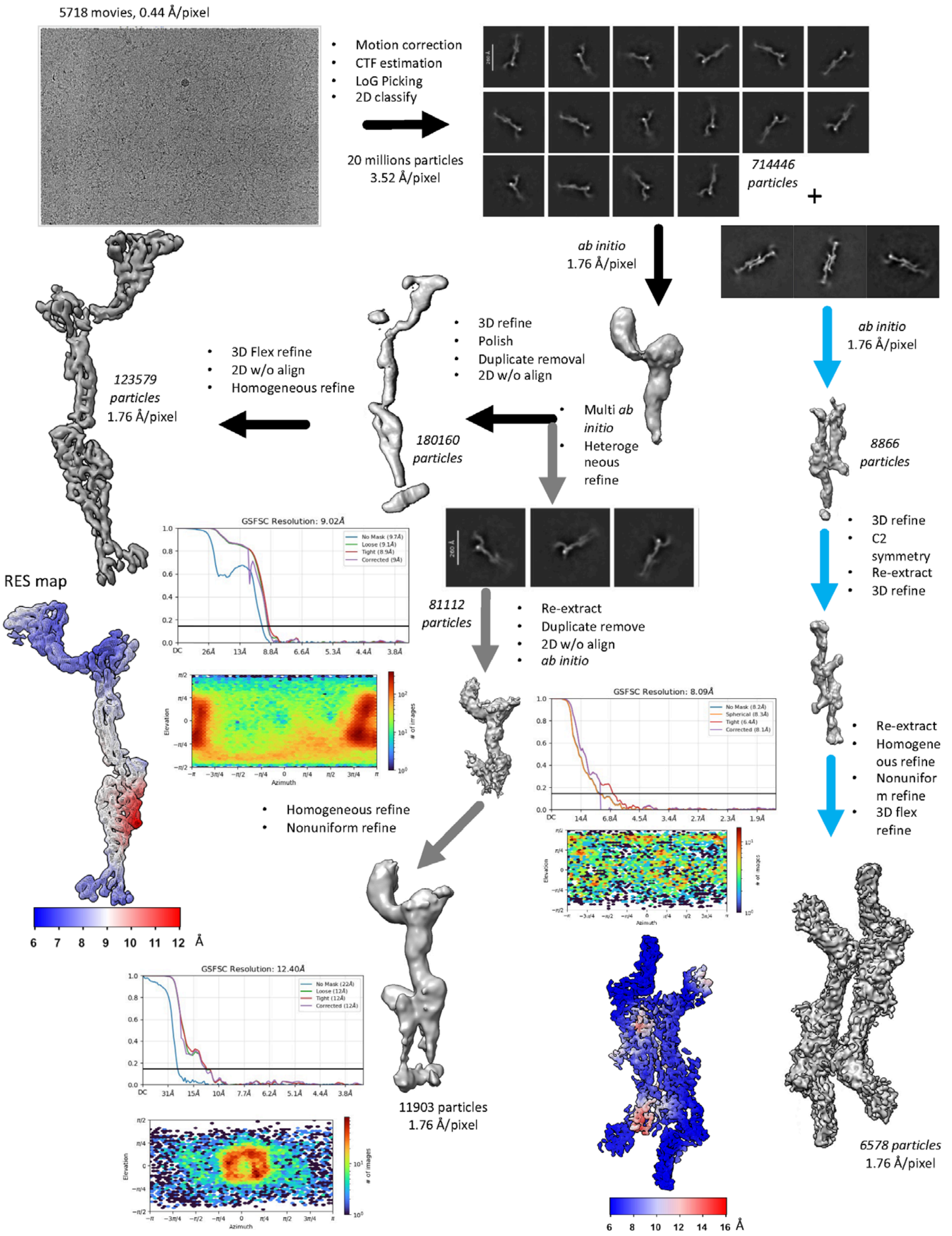

**Figure S2**: **Cryo-EM processing scheme for the full hetero-octameric (AUG1,2,3,4,5,6,7,8) Augmin leading to 10Å cryo-EM structure.**

Top to bottom, hetero-octameric (AUG 1,2,3,4,5,6,7,8) Augmin cryo-EM dataset (example image shown on left) was collected, and raw images were pre-processed using motioncorr2, CTFFind3 then used pick and identify coordinates for particle images, which were processed using a combination of RELION 3-4.0 and CryoSPARC 3.0-4.1. Multiple cycles of 2D classification identified three types of assemblies: top left 2D class averages, full Augmin assemblies, center middle, 2D-class averages, Augmin with additional extended regions (Augmin 1.5), and left 2D class averages, Augmin C2 symmetric dimers. Cycles of auto-3D refinement, flex-refine polishing, homogeneous refine in CryoSPARC 3.1 led to three reconstructions. These reconstruction are shown with their CryoSPARC angular distributions, Fourier shell correlation (FSC) curves and local resolution colored map (Res map). Center left, 10Å-reconstruction of full Augmin assembly. Lower center, a 12 Å reconstruction of Augmin 1.5 with additional extended domain. Lower right, 12Å C2 Augmin dimer reconstruction is shown. Final processing statistics are described in Table I.

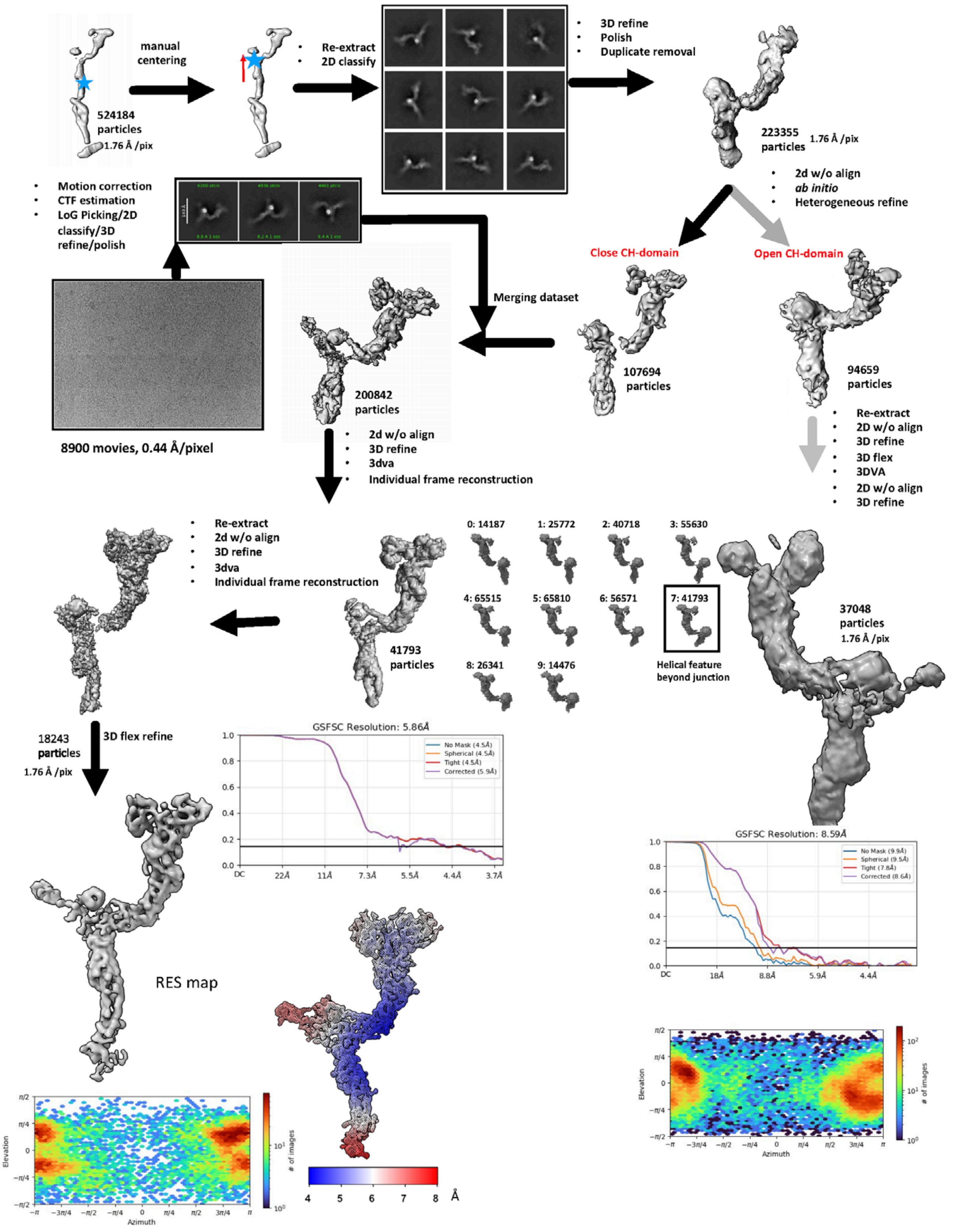
manual centering
524184 particles
1.76 Å /pix
Re-extract
2D classify
3D refine
Polish
Duplicate removal
223355 particles
1.76 Å /pix
2d w/o align
ab initio
Heterogeneous refine
Motion correction
CTF estimation
LoG Picking/2D classify/3D refine/polish
Close CH-domain
Open CH-domain
Merging dataset
107694 particles
94659 particles
200842 particles
8900 movies, 0.44 Å/pixel
2d w/o align
3D refine
3dva
Individual frame reconstruction
Re-extract
2D w/o align
3D refine
3D flex
3DVA
2D w/o align
3D refine
Re-extract
2d w/o align
3D refine
3dva
Individual frame reconstruction
0: 14187
1: 25772
2: 40718
3: 55630
4: 65515
5: 65810
6: 56571
7: 41793
8: 26341
9: 14476
Helical feature beyond junction
41793 particles
37048 particles
1.76 Å /pix
18243 particles
1.76 Å /pix
3D flex refine
GSFSC Resolution: 5.86Å
No Mask (4.5Å)
Spherical (4.5Å)
Tight (4.5Å)
Corrected (5.9Å)
GSFSC Resolution: 8.59Å
No Mask (9.9Å)
Spherical (9.5Å)
Tight (7.8Å)
Corrected (8.6Å)
RES map
4 5 6 7 8 Å
Elevation
Azimuth
### of images

**Figure S3**: **Cryo-EM processing scheme for the hetero-octameric (AUG1,2,3,4,5,6,7,8) Augmin V-junction stem to 7.3 to 12Å cryo-EM structure in the open and closed state.**

Top to bottom, the hetero-octameric (AUG 1,2,3,4,5,6,7,8) V-junction stem particles in **Figure S2** were re-centered and reextracted as shown in top left centering around the V-junction steam region, leading to refined class averages shown on top center. A second cryo-EM dataset for AUG1,2,3,4,5,6,7,8 assemblies (example image shown on left) was collected, and raw images were pre-processed using motioncor2, CTFFind3 then used pick and identify coordinates for particle images, which were processed using a CryoSPARC 4 leading to 2D class averages shown middle left center. Multiple cycles of 3D refinement, heterogenous refine identified two types of V-junction stem assemblies: closed dual CH-dimer Head domain Left, open dual CH-dimer head domain. Particles from both datasets were merged for each of these two. Center middle, 3D classification led to a single class with improved features for the closed state. The dual CH-dimer closed state particles were re-extracted at smaller pixel size 3D auto-refined, 3D-variability refined (3DVA) then flex refined leading to a reconstruction at 7Å resolution. Particles for the open state were auto-refined and leading to a 10Å reconstruction of the Augmin V-junction-stem at in the open CH-dimer state.

Figure S4

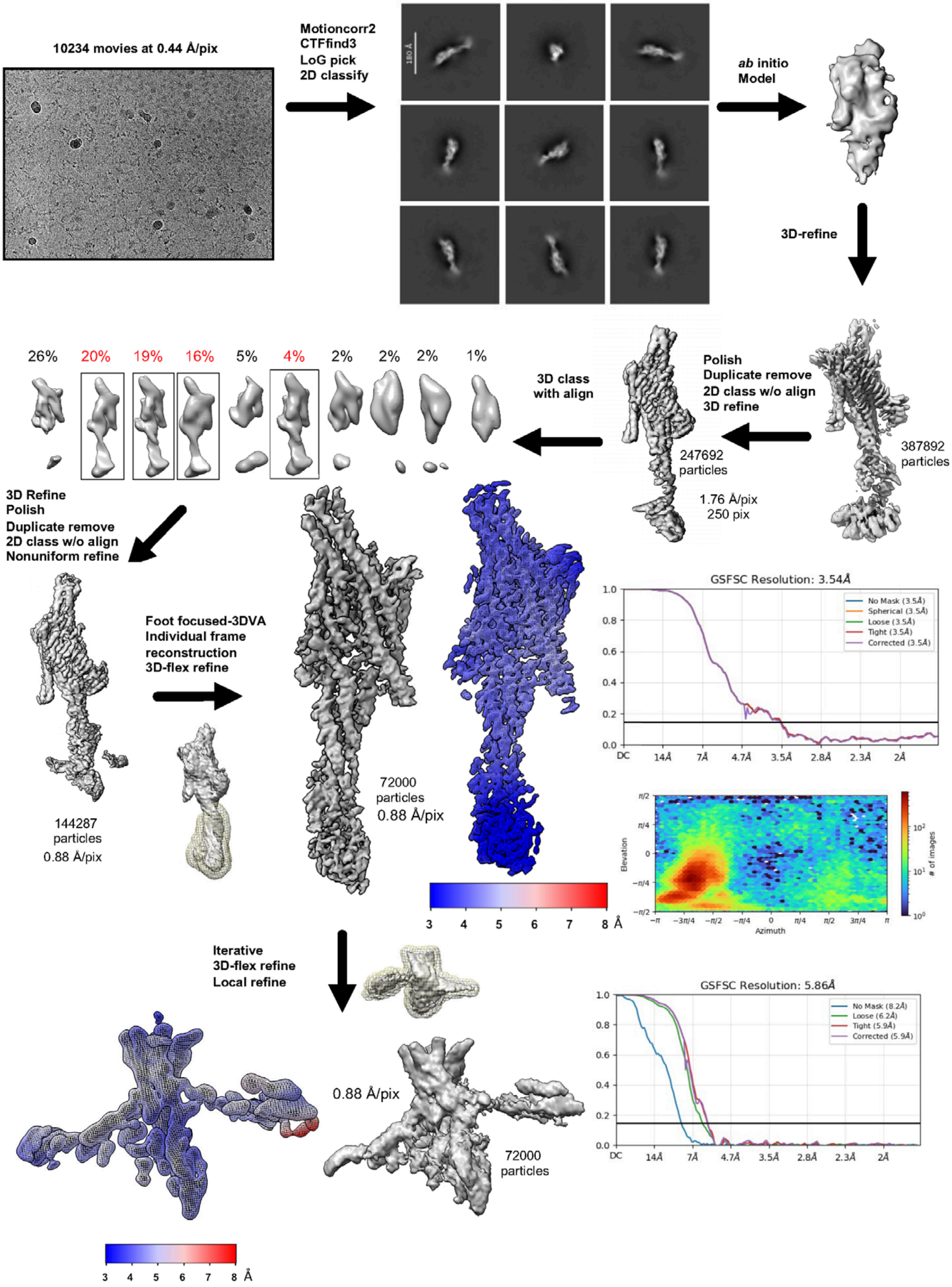

10234 movies at 0.44 Å/pix
Motioncorr2
CTFfind3
LoG pick
2D classify
180 Å
ab initio
Model
3D-refine
Polish
Duplicate remove
2D class w/o align
3D refine
387892
particles
247692
particles
1.76 Å/pix
250 pix
3D class
with align
26%
20%
19%
16%
5%
4%
2%
2%
2%
1%
3D Refine
Polish
Duplicate remove
2D class w/o align
Nonuniform refine
144287
particles
0.88 Å/pix
Foot focused-3DVA
Individual frame
reconstruction
3D-flex refine
72000
particles
0.88 Å/pix
3 4 5 6 7 8 Å
GSFSC Resolution: 3.54Å
No Mask (3.5Å)
Spherical (3.5Å)
Loose (3.5Å)
Tight (3.5Å)
Corrected (3.5Å)
DC 14Å 7Å 4.7Å 3.5Å 2.8Å 2.3Å 2Å
Elevation
Azimuth
### of images
Iterative
3D-flex refine
Local refine
0.88 Å/pix
72000
particles
GSFSC Resolution: 5.86Å
No Mask (8.2Å)
Loose (6.2Å)
Tight (5.9Å)
Corrected (5.9Å)
DC 14Å 7Å 4.7Å 3.5Å 2.8Å 2.3Å 2Å
3 4 5 6 7 8 Å

**Figure S4: Cryo-EM processing scheme for the hetero-tetrameric (AUG1,3,4,5) Augmin leading to a 3.7-6.0 A resolution map for the extended region**

Top to bottom, cryo-EM data movies for hetero-tetrameric (AUG 1,3,4,5) Augmin particles (example image on top left) was collected then pre-processed, using motioncorr2, CTFFind3 then used pick and identify coordinates for particle images, as described, leading to 2D-class averages shown on top center. Class averages were used to generate ab initio model top right, which was refined and then further polished and duplicate removed. This is followed by 3D classification with alignment leading to four classes which were combined, then followed the step shown in center left leading to a refined structure at 3.7Å resolution for the extended region showing clear helical and side chain density shown in center. The lower section of the extended region was 3D-flex refined leading to a 6Å structure for the tripod region.

Figure S5

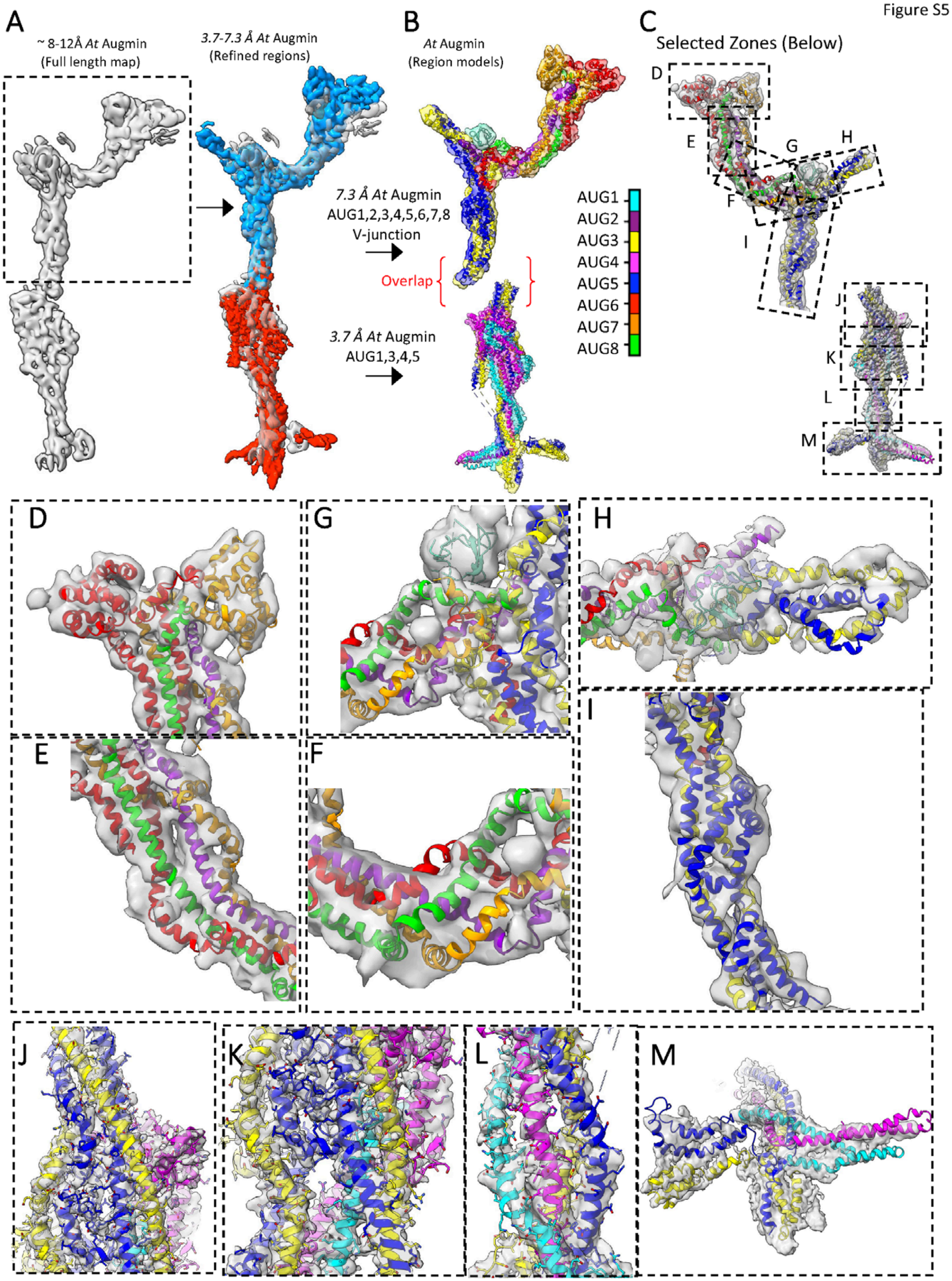
A
~ 8-12Å *At* Augmin
(Full length map)
*3.7-7.3 Å At* Augmin
(Refined regions)
B
*At* Augmin
(Region models)
*7.3 Å At* Augmin
AUG1,2,3,4,5,6,7,8
V-junction
Overlap
*3.7 Å At* Augmin
AUG1,3,4,5
AUG1
AUG2
AUG3
AUG4
AUG5
AUG6
AUG7
AUG8
C
Selected Zones (Below)
D
E
F
G
H
I
J
K
L
M

**Figure S5**: **Composite maps used to generate a full *de novo* Augmin hetero-octamer model**

A) Right, 10-ÅCryo-EM map for Full Augmin hetero-octamer (AUG1,2,3,4,5,6,7,8) generated as presented in **Figure S2**. Left, aligned cryo-EM map (transparent white) as shown on left, overlaid with 7.3Å map for the V-junction stem (blue) generated as described in **Figure S3**, and the 3.7Å AUG1,3,4,5 Augmin extended region map generated as described in **Figure S4**.

B) Segmented V-junction-stem map fitted with modeled AUG1,2,3,4,5,7,8 regions is shown on the top, with the head region of AUG6,7 CH-domain dimer in the closed state. The extended region map is shown on the bottom. The overlap region composed of four helices from AUG3,5 is marked by red parentheses. Subunit color guide shown in on right.

C) A Guide to close up views of the modeled regions shown in B (rotated by 180° from C). The V-junction and stem map zones marked with D-I boxes, representing different regions. The extended region is shown below with zones marked with J-M boxes representing different regions

D-I) Close-up views for different regions of V-junction stem model to map fits with guide to subunit colors shown in B

J-M) Close-up views for different regions of the extended region model to map fits with guide to subunit colors shown in B.

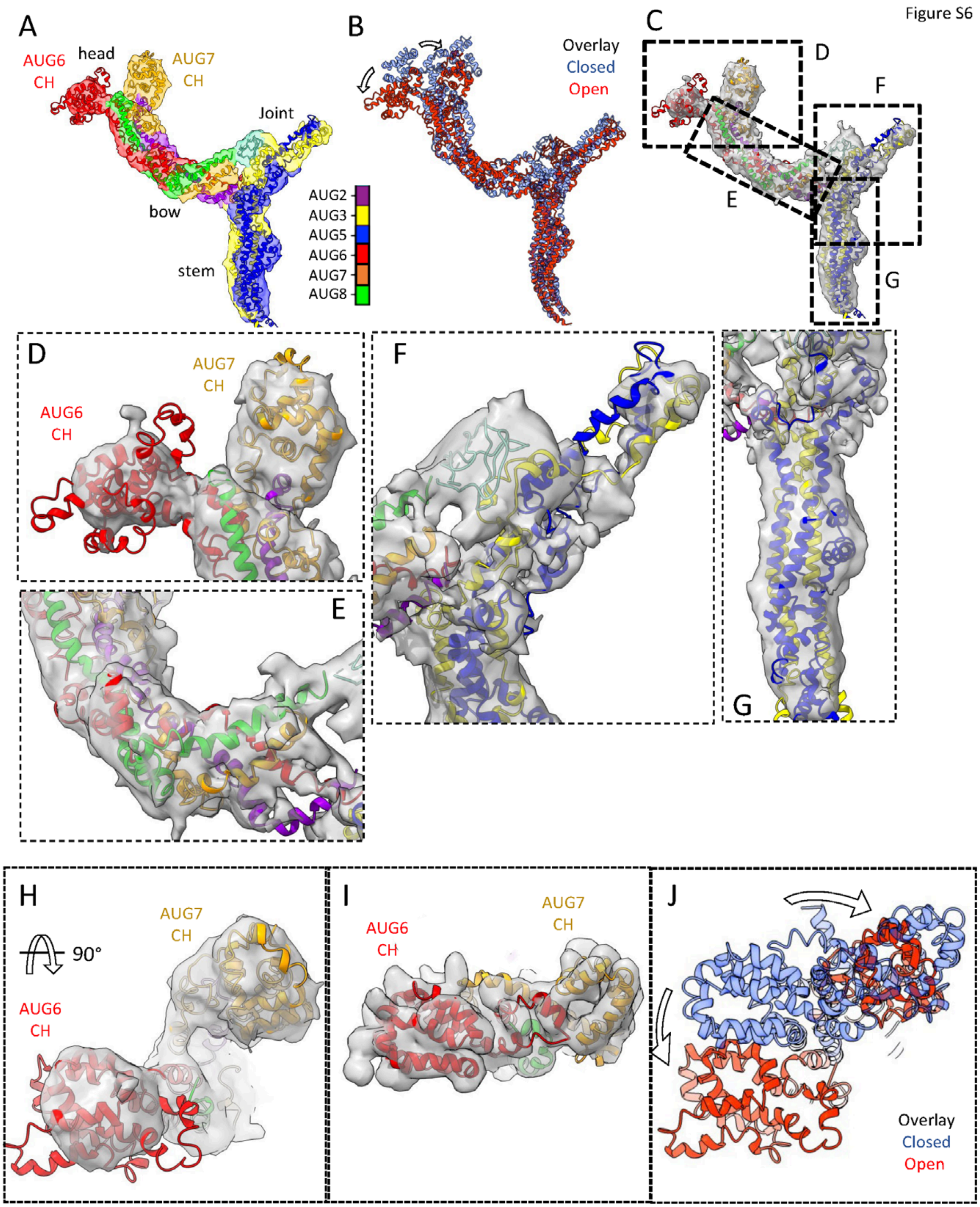
Figure S6
A
AUG6
CH
head
AUG7
CH
Joint
bow
stem
AUG2
AUG3
AUG5
AUG6
AUG7
AUG8
B
Overlay
Closed
Open
C
D
F
E
G
D
AUG7
CH
AUG6
CH
E
F
G
H
AUG7
CH
90°
AUG6
CH
I
AUG6
CH
AUG7
CH
J
Overlay
Closed
Open

**Figure S6**: **Map to model view for the Augmin V-junction-stem in AUG6,7 CH dimer open state**

A) Segmented map fitted with modeled AUG1,2,3,4,5,7,8 regions into the V-junction-stem map (top) with the head region AUG6,7 CH-domain dimer in the open state. Subunit color guide shown in on right.
B) Overlay of the open state V-junction and closed state V-junction as shown in **Figure 2B**
C) A Guide to close up views of the modeled regions shown in A. The V-junction and stem map zones marked with D-I boxes.

D-G) Close-up views for different regions of V-junction stem model to map fits with guide to subunit colors shown in A.

H) Top end view of the head domain AUG6,7 CH-domain dimer in the open state

I) Top end view of the head domain AUG6,7 CH-domain dimer in the closed state

J) Top end view of the overlay of the ch-AUG6,7 dimer in the open and closed state shown in red and blue, respectively, showing the conformational transition of the CH-domains, suggesting AUG7 CH-domain moves more extensively than AUG6 Ch in the opening transition.

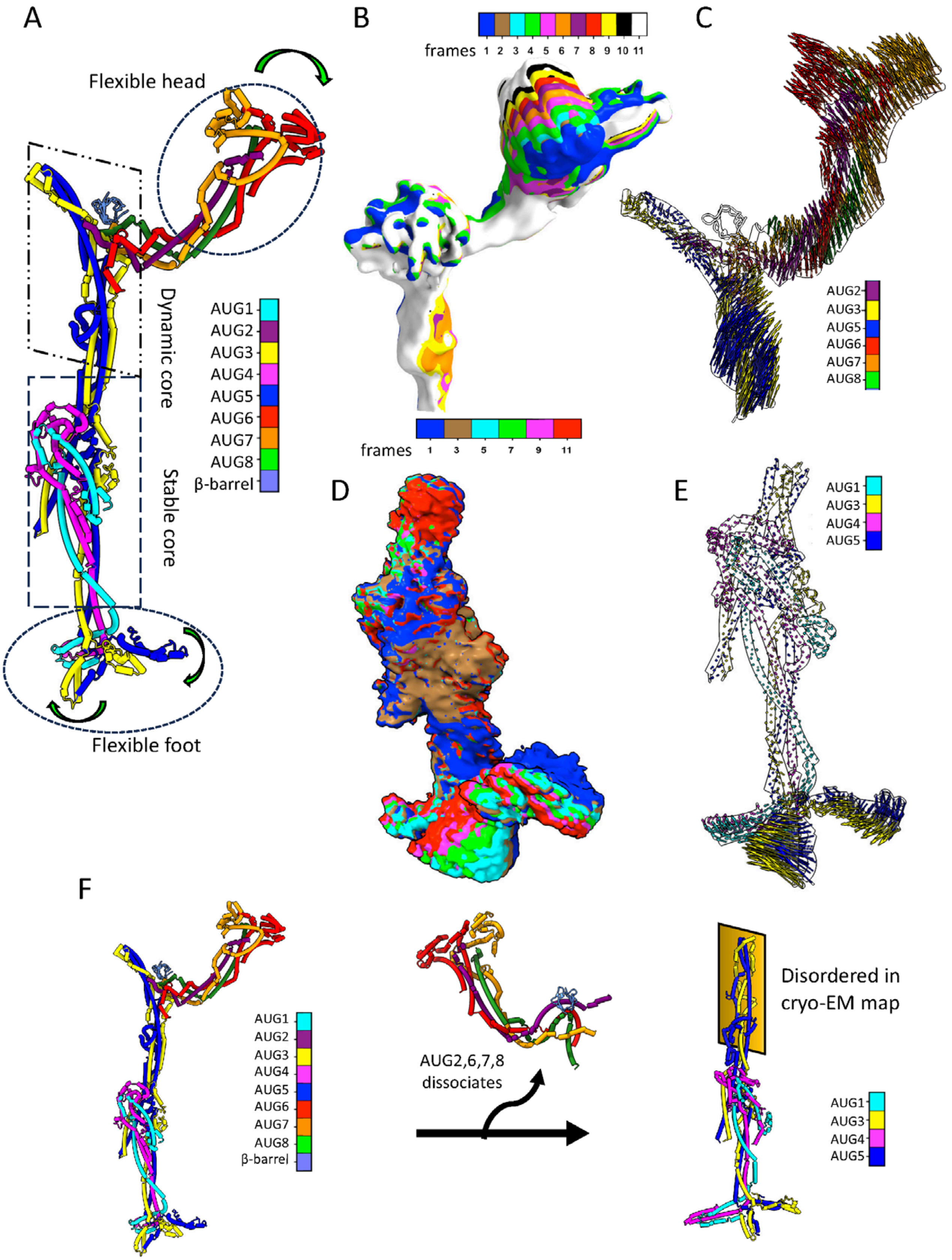
A
Flexible head
Dynamic core
Stable core
Flexible foot
AUG1
AUG2
AUG3
AUG4
AUG5
AUG6
AUG7
AUG8
β-barrel
B
frames 1 2 3 4 5 6 7 8 9 10 11
frames 1 3 5 7 9 11
C
AUG2
AUG3
AUG5
AUG6
AUG7
AUG8
D
E
AUG1
AUG3
AUG4
AUG5
F
AUG1
AUG2
AUG3
AUG4
AUG5
AUG6
AUG7
AUG8
β-barrel
AUG2,6,7,8 dissociates
Disordered in cryo-EM map
AUG1
AUG3
AUG4
AUG5

**Figure S7**: **Regional Flexibilities observed in Augmin particles based on frame-based 3D-flex**

A) Tube cartoon model for the Augmin hetero octamer showing the flexible regions in the assembly. The V-junction head domain and bow region show flexibility and the tripod region at the other end shows flexibility
B) Low resolution frame maps for the V-junction stem region, generated in the 3D-flex processing that marked with colors as shown in the guideline. These maps differ in the movement rotation of the V-junction bow and the head regions
C) Vectorial movement model showing the regions of the V-junction and stem that undergo movement. The magnitude of these movements is demonstrated by longer vector lines. The subunit color guide describes the subunits presented in the region.
D) Low resolution frame maps for the extended region, generated in the 3DFlex processing that marked with colors as shown in the guideline. These maps differ in the movement rotation of the leg region and the connected tripod region at the lowest part of the map.
E) Vectorial movement model showing the regions of the extended region map that undergo movement. The magnitude of these movements is demonstrated by size of the vector lines. The subunit color guide describes the subunits presented in the region.
F) A model describing the impact of the dissociation of AUG2,6,7,8 subcomplex leading to the destabilization of the structural stability of the AUG3,5 foldback zone in the extended region structure represented by AUG1,3,4,5

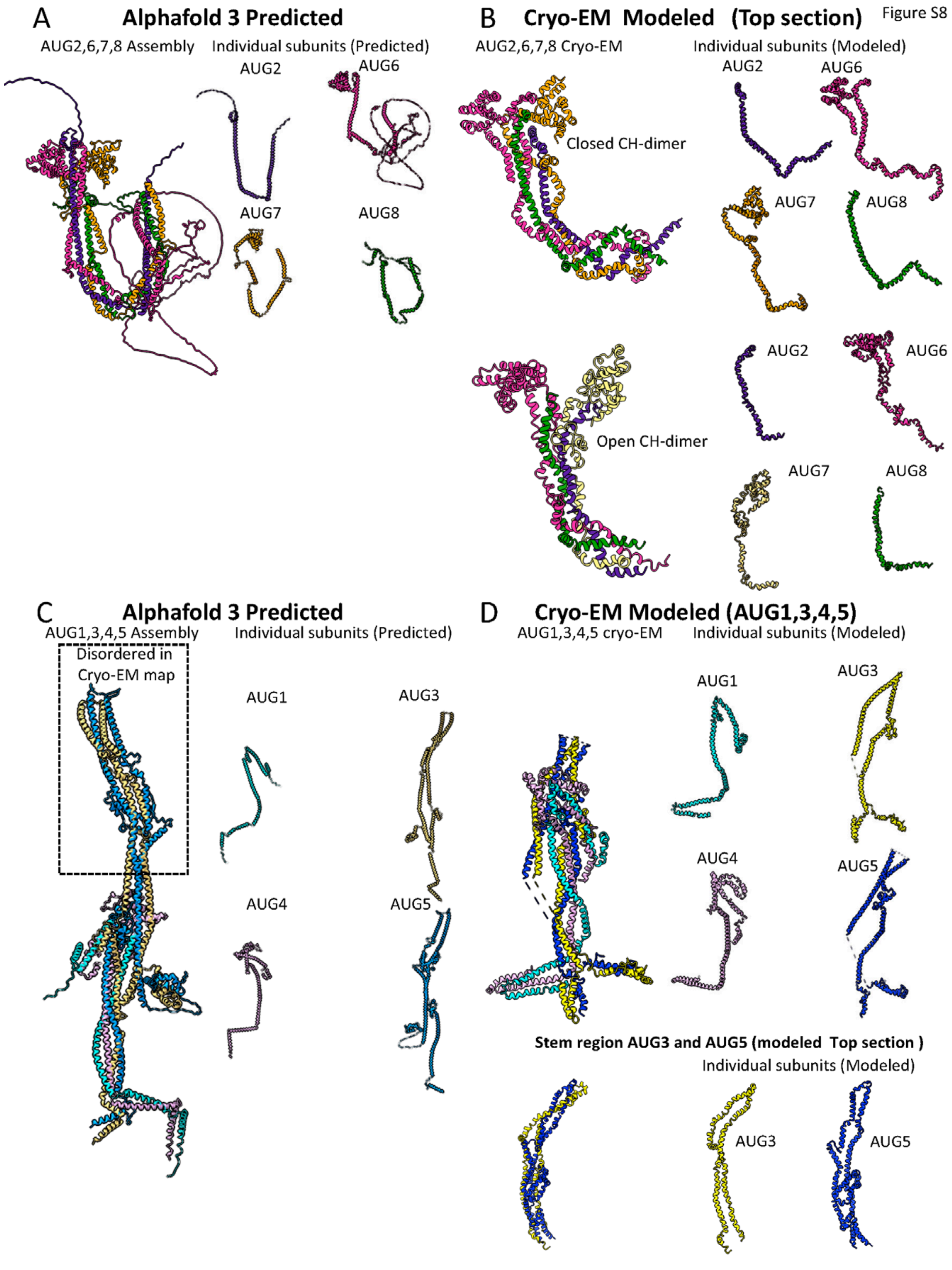
Figure S8
A
Alphafold 3 Predicted
AUG2,6,7,8 Assembly
Individual subunits (Predicted)
AUG2
AUG6
AUG7
AUG8
B
Cryo-EM Modeled (Top section)
AUG2,6,7,8 Cryo-EM
Individual subunits (Modeled)
AUG2
AUG6
Closed CH-dimer
AUG7
AUG8
AUG2
AUG6
Open CH-dimer
AUG7
AUG8
C
Alphafold 3 Predicted
AUG1,3,4,5 Assembly
Individual subunits (Predicted)
Disordered in Cryo-EM map
AUG1
AUG3
AUG4
AUG5
D
Cryo-EM Modeled (AUG1,3,4,5)
AUG1,3,4,5 cryo-EM
Individual subunits (Modeled)
AUG1
AUG3
AUG4
AUG5
Stem region AUG3 and AUG5 (modeled Top section )
Individual subunits (Modeled)
AUG3
AUG5

**Figure S8**: **Comparison of Alphafold3 and cryo-EM built Models of Augmin assemblies and isolated subunits**

A) Left, AlphaFold 2 ribbon model for the *At* AUG2,6,7,8 assembly: right model for isolated AUG2,6,7,8 subunits
B) Top left, Cryo-EM ribbon model for the *At* AUG2,6,7,8 assembly in the CH-dimer closed state; top right model for isolated AUG2,6,7,8 subunits. Bottom left, Cryo-EM ribbon model for the *At* AUG2,6,7,8 assembly in the CH-dimer open state; bottom right model for isolated AUG2,6,7,8 subunits
C) Left, AlphaFold 2 ribbon model for the *At* AUG1,3,4,5 assembly with a box marking the AUG3,5-foldback zone, which is not observed in the AUG1,3,4,5 cryo-EM map of the extended region: right, models for isolated AUG1,3,4,5 subunits. Note the open shape of the end of the extended domain.
D) Left, Cryo-EM ribbon model for the *At* AUG1,3,4,5 assembly of the extended region; Top right, models for isolated AUG1,3,4,5 subunits in the AUG,1,3,4,5 extended region map. Bottom right, models for the AUG3, AUG5 and their assembly in the foldback zone observed in the AUG1,3,4,5,6,7,8 map of the V-junction and stem.

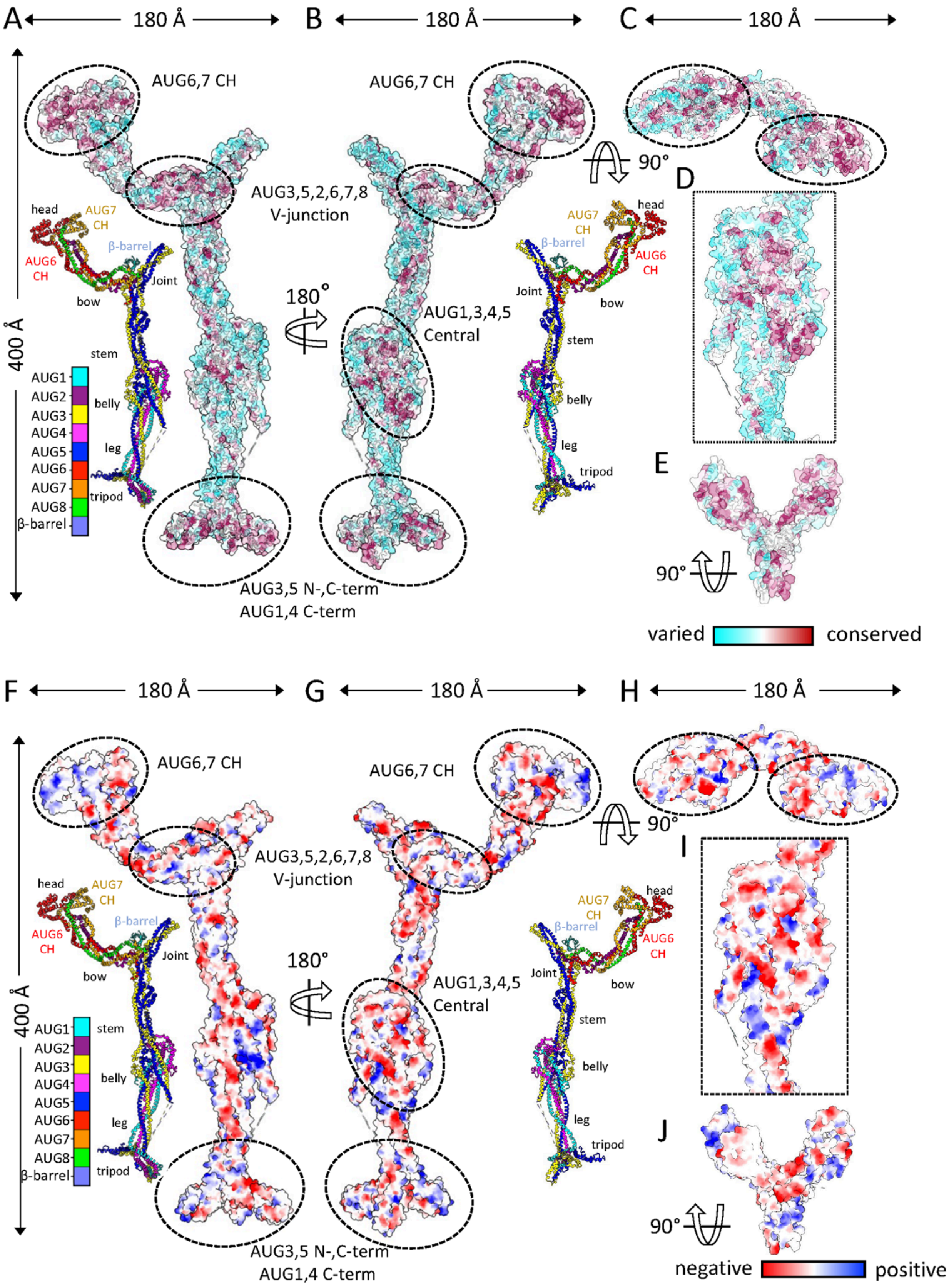

A
180 Å
B
180 Å
C
180 Å
400 Å
AUG6,7 CH
AUG6,7 CH
90°
AUG3,5,2,6,7,8
V-junction
D
head
AUG7
CH
AUG6
CH
β-barrel
Joint
bow
stem
180°
AUG1,3,4,5
Central
AUG1
AUG2
AUG3
AUG4
AUG5
AUG6
AUG7
AUG8
β-barrel
belly
leg
tripod
E
90°
AUG3,5 N-,C-term
AUG1,4 C-term
varied
conserved
F
180 Å
G
180 Å
H
180 Å
400 Å
AUG6,7 CH
AUG6,7 CH
90°
AUG3,5,2,6,7,8
V-junction
I
head
AUG7
CH
AUG6
CH
β-barrel
Joint
bow
180°
AUG1,3,4,5
Central
stem
belly
leg
tripod
J
90°
AUG3,5 N-,C-term
AUG1,4 C-term
negative
positive

**Figure S9**: **Charge distribution and Sequence Conservation mapped on Augmin hetero-octamer (AUG1,2,3,4,5,6,7,8) model**

A-C) A and B show two side views of the sequence conservation plotted by color on the Augmin surface model. The model is rendered using the color scheme shown on the top right. Inset models shown. A and B show the two inset side views of the Augmin ribbon model in the same views to help orient the view to the structure. C, Top shows the top end view of the V-junction. Middle shows close up view on the belly region shown in B. Bottom shows the view from the below the Tripod region. The surface conservation of the Augmin assembly structure suggests regions of critical functional importance in the Augmin structure which are labeled by ellipses.

D-F) D and D show two side views of the surface charge distribution plotted by color on the Augmin surface model. The model is rendered using the color scheme shown on the top right. Inset models shown. D and E show the inset two side views of the Augmin ribbon model in the same views to help orient the view to the structure. F, Top shows the top end view of the V-junction. Middle shows close up view on the belly region shown in E. Bottom shows the view from the below the Tripod region. The charge distribution marks the four critical regions of importance in the Augmin structure which are labeled by ellipses.

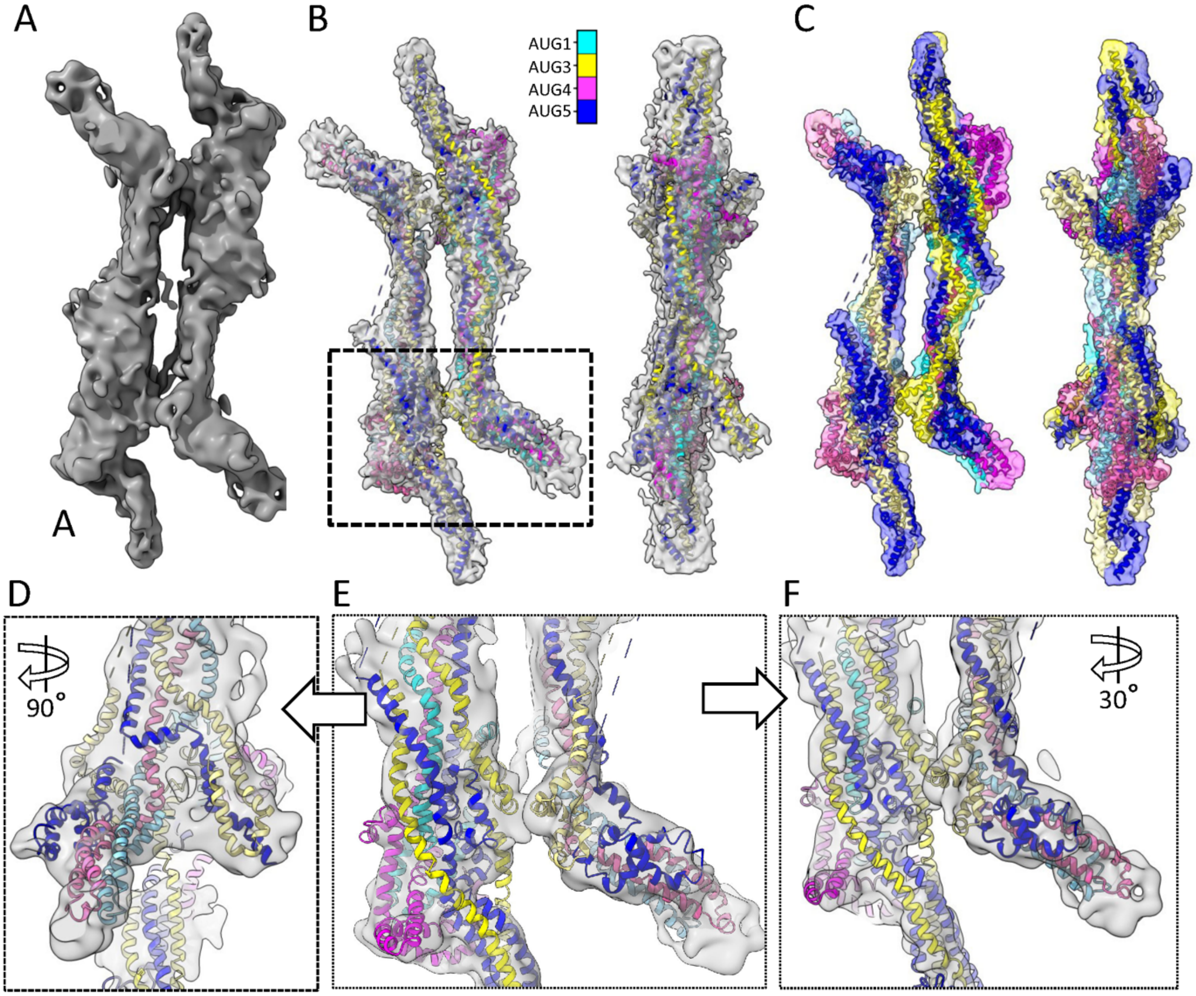


**Figure S10**: **Details of the Augmin dimer cryo-EM map to model**

A) The AUG1,2,3,4,5,6,7,8 Augmin C2 dimer map generated as described in **Figure S2**.

B) Two views of the assemblies of AUG1,3,4,5 built into the raw cryo-EM map (grey).

C) Two views of the assemblies of AUG1,3,4,5 built presented into the segmented cryo-EM map (multi-color). Subunits and segments are presented in the colors described in the palette above.

D-F) three rotated views of the Augmin Dimer map interfaces between the folded tripod region and the belly region. The map is shown in raw grey color while each subunit is presented in ribbon format following the color pallet

Figure S11

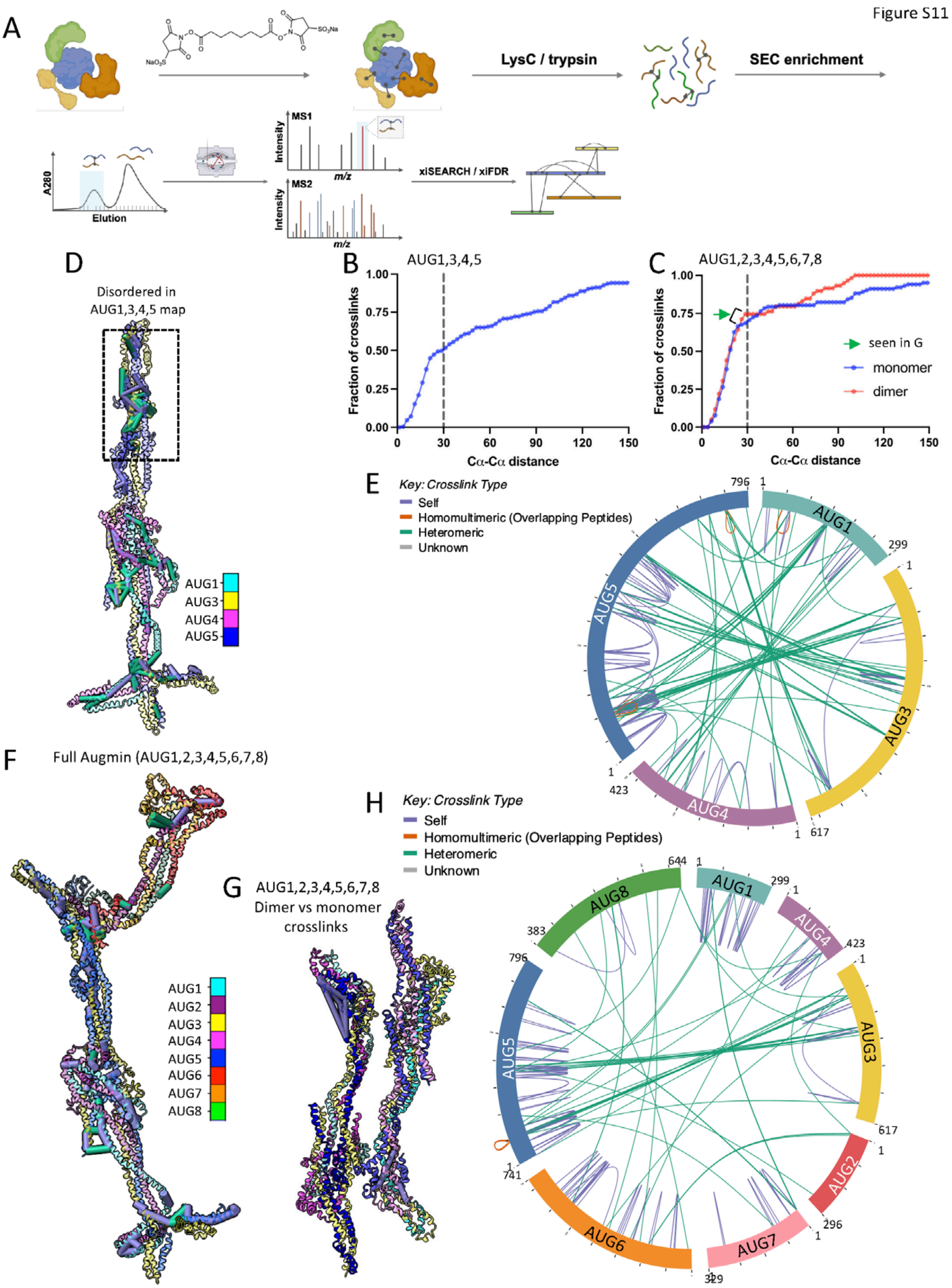
A
LysC / trypsin
SEC enrichment
A280
Elution
MS1
Intensity
m/z
MS2
xiSEARCH / xiFDR
B
AUG1,3,4,5
Fraction of crosslinks
Cα-Cα distance
C
AUG1,2,3,4,5,6,7,8
seen in G
monomer
dimer
D
Disordered in AUG1,3,4,5 map
AUG1
AUG3
AUG4
AUG5
E
Key: Crosslink Type
Self
Homomultimeric (Overlapping Peptides)
Heteromeric
Unknown
F
Full Augmin (AUG1,2,3,4,5,6,7,8)
AUG1
AUG2
AUG3
AUG4
AUG5
AUG6
AUG7
AUG8
G
AUG1,2,3,4,5,6,7,8
Dimer vs monomer crosslinks
H
Key: Crosslink Type
Self
Homomultimeric (Overlapping Peptides)
Heteromeric
Unknown

**Figure S11: Crosslinking Mass spectrometry (XLMS) of AUG1,3,4,5 and AUG1,2,3,4,5,6,7,8**

A) Scheme for XLMS: Assemblies are treated with crosslinker (BS3) and then the crosslinked assemblies are unfolded and proteolyzed. The proteolyzed collection of protein peptides are run on HPLC followed by mass spectrometry and the identity of paired region is determined
B) CDF plot for hetero-tetrameric (AUG1,3,4,5) Augmin showing the $C\alpha$-$C\alpha$ distance plotted as a function of percentage of crosslinks explained by the model. The line denotes 30Å distance indicating ~50% of the crosslinks can be explained by the model.
C) CDF plot for hetero-octameric (AUG1,2,3,4,5,6,7,8) Augmin showing the $C\alpha$-$C\alpha$ distance plotted as a function of percentage of crosslinks explained by the Augmin monomer (blue) or anti-parallel Augmin (red) model. The line denotes 30 Å indicating 70% and 75% of the crosslinks can be explained by Augmin monomer (Blue) and dimer (red) model. Differences between crosslinks are marked by green arrow and plotted in G.
D) Cryo-EM generated hetero-tetrameric (AUG1,3,4,5) Augmin model presented in ribbon format with the observed crosslinks generated in tube format. Note the region in AUG3,5 foldback zone are not observed in the cryo-EM structure but extensive crosslinks are observed. The tripod and the foldback zones are likely extensively flexible.
E) XLMS raw crosslinks showing the circular view of AUG1,3,4,5 revealing the large number inter and intramolecular crosslinks
F) Cryo-EM generated Hetero-octamer (AUG1,2,3,4,5,6,7,8) Augmin model presented in ribbon format with the observed crosslinks generated as tubes between crosslink sites
G) Left, difference crosslinks between hetero-octamer Augmin monomer and anti-parallel dimer model plotted on the ribbon model.
H) XLMS raw crosslinks showing the circular view of AUG1,3,4,5, revealing the large number inter and intramolecular crosslinks

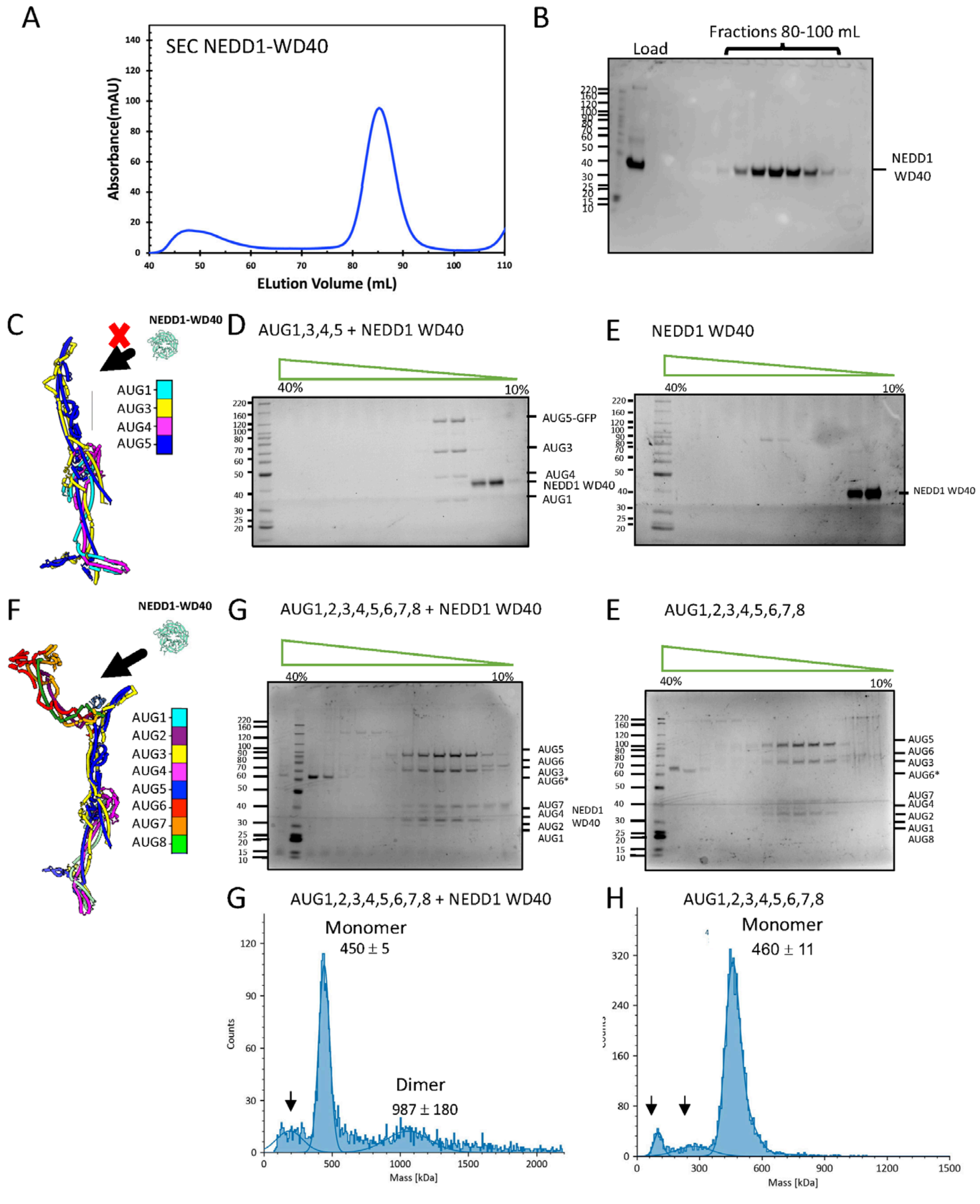
A
SEC NEDD1-WD40
Absorbance(mAU)
ELution Volume (mL)
B
Load
Fractions 80-100 mL
NEDD1
WD40
C
NEDD1-WD40
AUG1
AUG3
AUG4
AUG5
D
AUG1,3,4,5 + NEDD1 WD40
40%
10%
AUG5-GFP
AUG3
AUG4
NEDD1 WD40
AUG1
E
NEDD1 WD40
40%
10%
NEDD1 WD40
F
NEDD1-WD40
AUG1
AUG2
AUG3
AUG4
AUG5
AUG6
AUG7
AUG8
G
AUG1,2,3,4,5,6,7,8 + NEDD1 WD40
40%
10%
AUG5
AUG6
AUG3
AUG6*
AUG7
AUG4
AUG2
AUG1
NEDD1
WD40
E
AUG1,2,3,4,5,6,7,8
40%
10%
AUG5
AUG6
AUG3
AUG6*
AUG7
AUG4
AUG2
AUG1
AUG8
G
AUG1,2,3,4,5,6,7,8 + NEDD1 WD40
Monomer
450 ± 5
Dimer
987 ± 180
Counts
Mass [kDa]
H
AUG1,2,3,4,5,6,7,8
Monomer
460 ± 11
Mass [kDa]

**Figure S12: Purification of *At* NEDD1 WD40-β-propellor and biochemical reconstitution of NEDD1-WD40-β-propellor with AUG1,3,4,5 and AUG1,2,3,4,5,6,7,8.**

A) SEC purification chromatogram of insect cell expressed At NEDD1-WD40 β-propellor.
B) SDS-PAGE of SEC-purified NEDD1-WD40 fractions revealing the solubility and monodisperse behavior of the protein as globular entity.
C) 10-40% Sucrose density gradient of AUG1,3,4,5 with NEDD1-WD40 showing the isolated and non-overlapping and separate migration of these two entities
D) 10-40% Sucrose density gradient of NEDD1-WD40 showing its matching migration of pattern alone with panel C.
E) 10-40% Sucrose density gradient of AUG1,2,3,4,5,6,7,8 with NEDD1-WD40 showing they co-elute together as a collection of proteins with fractions showing their overlay or migration.
F) 10-40% Sucrose density gradient of AUG1,2,3,4,5,6,7,8 showing the subunits co-elute forming a single entity with co-migration of all relevant subunits.

Figure S13

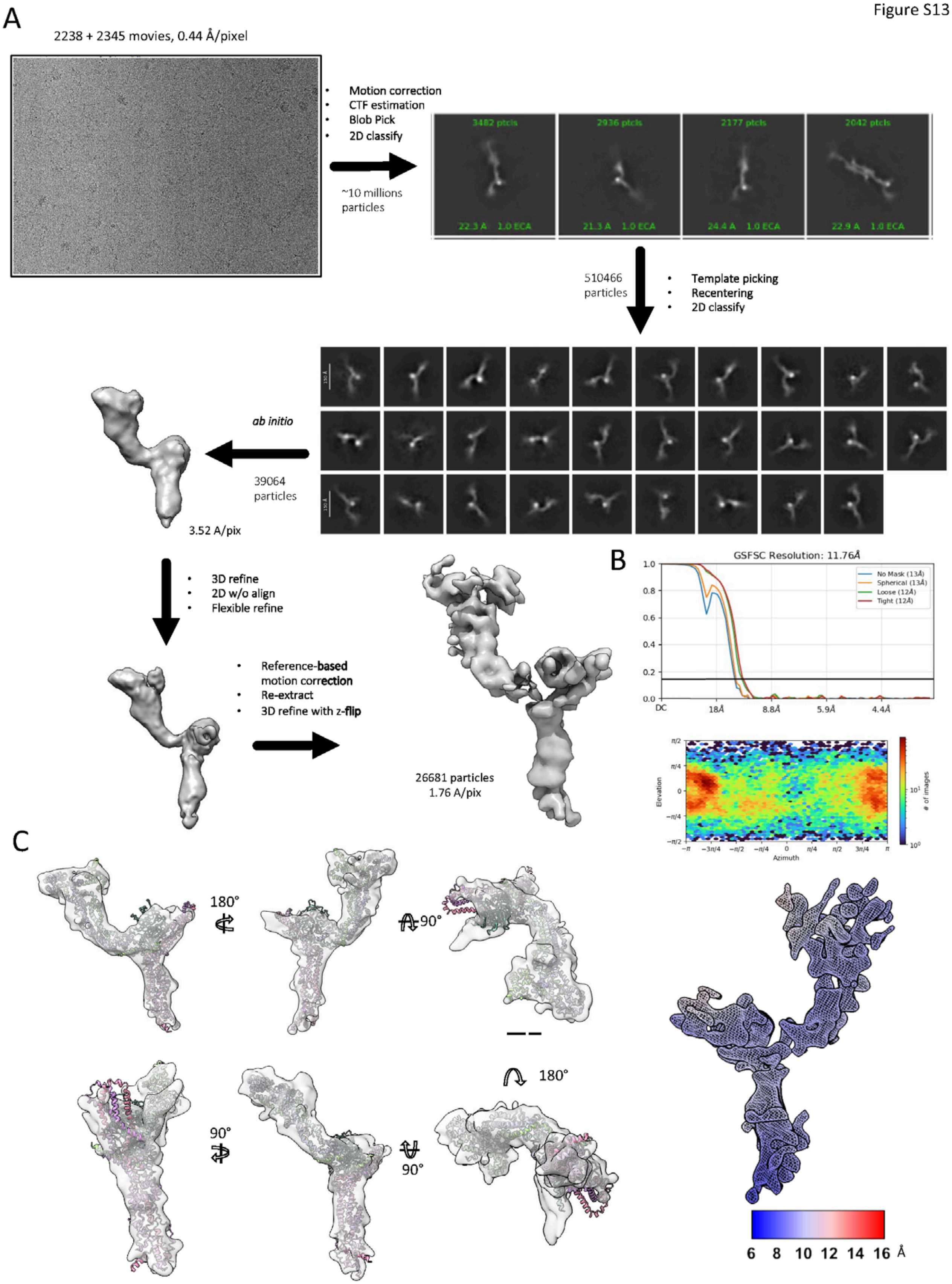
A
2238 + 2345 movies, 0.44 Å/pixel
Motion correction
CTF estimation
Blob Pick
2D classify
~10 millions particles
3482 ptcls
2936 ptcls
2177 ptcls
2042 ptcls
510466 particles
Template picking
Recentering
2D classify
ab initio
39064 particles
3.52 A/pix
3D refine
2D w/o align
Flexible refine
Reference-based motion correction
Re-extract
3D refine with z-flip
26681 particles
1.76 A/pix
B
GSFSC Resolution: 11.76Å
No Mask (13Å)
Spherical (13Å)
Loose (12Å)
Tight (12Å)
Elevation
Azimuth
### of images
C
180°
90°
90°
90°
180°
6 8 10 12 14 16 Å

**Figure S13: Cryo-EM structure determination of the NEDD1-WD40 β-propellor-Augmin-V-junction and stem region**.

A) From top to bottom: Cryo-EM data for AUG1,2,3,4,5,6,7,8 +NEDD1 WD40 assemblies (representative image shown on top left) were pre-processed with CTFfind3, motioncor2.1 then picked and 2D-classified with many rounds of 2D-classification leading a mixture of full monomeric or dimeric assemblies Template picking and recentering and 2D-classification led improved 2D-classes of V-junction-stem regions of Augmin. We note the density of the signal, likely representing the NEDD1-WD40 binding site, at the top of the V-junction is higher than Augmin -V-junction top section alone. These 2D-class average images were then to generate an *ab initio* model which was 3D-auto-refined. These data were subjected to reference-based refinement and 3DFlex refine leading to a 10 Å resolution structure.
B) Top, Fourier Shell correlation (FSC) for the final NEDD1-WD40 bound AUG1,2,3,4,5,6,7,8 V-junction stem. Middle, the angular distribution of the resulting map. Bottom,
C) Model comparison of the NEDD1-WD40-AUG1,3,4,5 V-junction-stem region map to a model generated for the complex showing the close overlay in many orientations, particularly the large size of the density representing NEDD1 compared density in AUG1,2,3,4,5,6,7,8 which is bound by the weaker β-barrel containing densities.

Figure S14

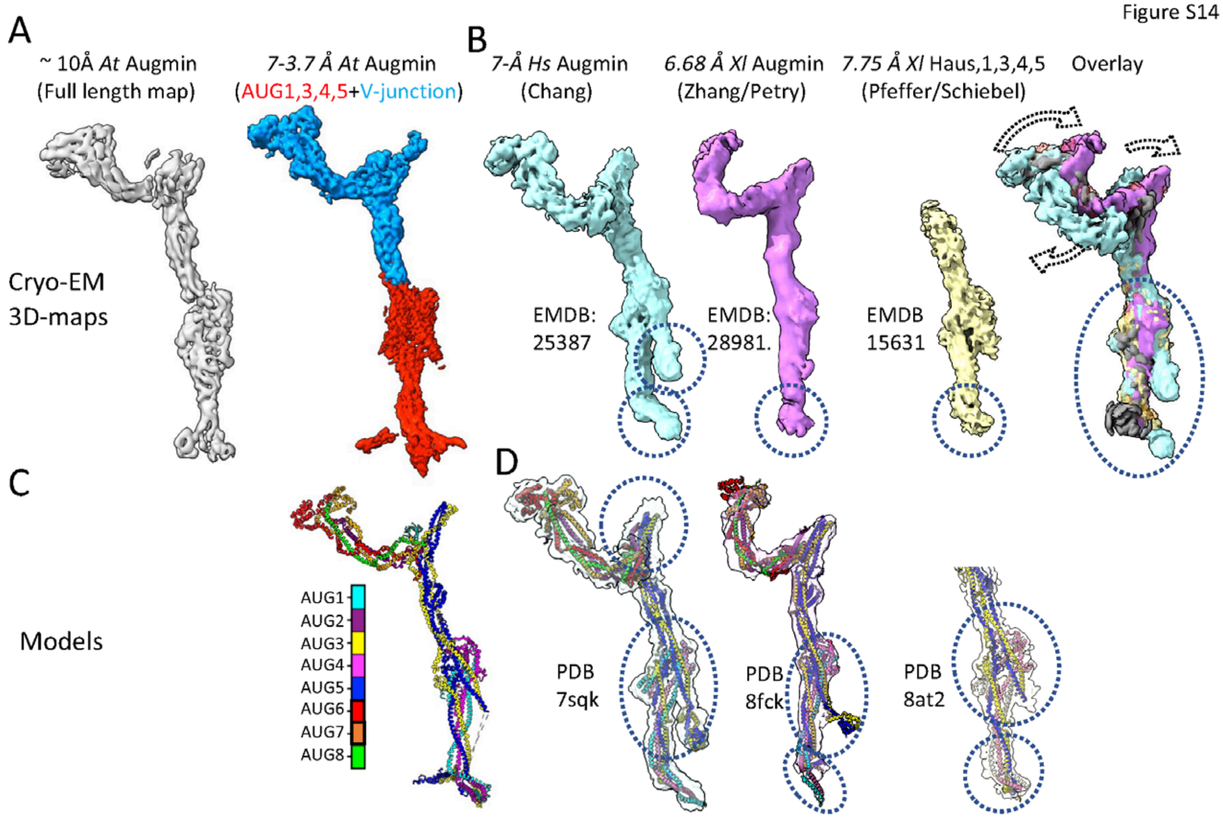


**Figure S14: comparison of Our Plant Augmin structures to previously published Augmin structures**

A) Left, full Augmin assembly map generated as described in **Figure 2**; right the overlay of the full Augmin assembly map onto the 7.3Å Augmin V-junction-stem map (blue) and 3.7-Å extended region domain map (red).

B) Left, the cryo-EM map for full Augmin generated Gabel *et al* (cyan). Second from left, the cryo-EM map for full Augmin generated by Travis *et al* (purple). Third left, cryo-EM map of the Haus1,3,4,5 region generated by Zuppa *et al* (purple)[29-31].

C) *De novo* At Augmin assembly presented in this manuscript

D) Left, Full model fit into the Cryo-EM density map placed into the *Gabel et al* (pink); second from left, Travis *et al*. Right, Zuppa *et al* and colleagues model fit into the cryo-EM of Haus 1,3,4,5 assemblies showing its nearly matching the shape and size of the AUG1,3,4,5 structure presented here[29-31].
