## Supplementary figures and images for "Cryo-EM structures of the Plant Augmin reveal its intertwined coiled-coil assembly, antiparallel dimerization and NEDD1 binding mechanisms"

### Figure S1

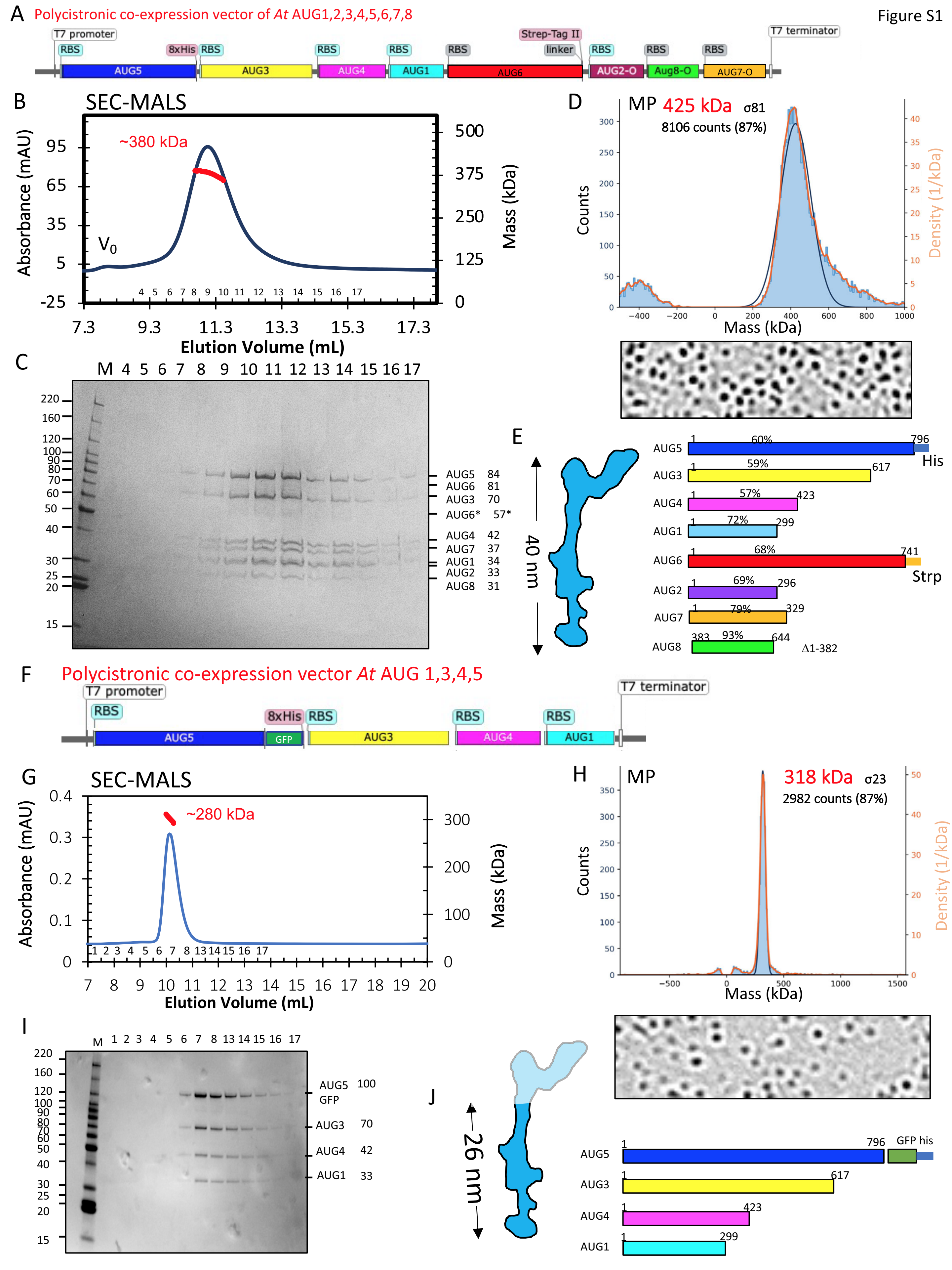

### Figure S2

Cryo-EM processing of the Aug1,2,3,4,5,6,7,8

Figure S2

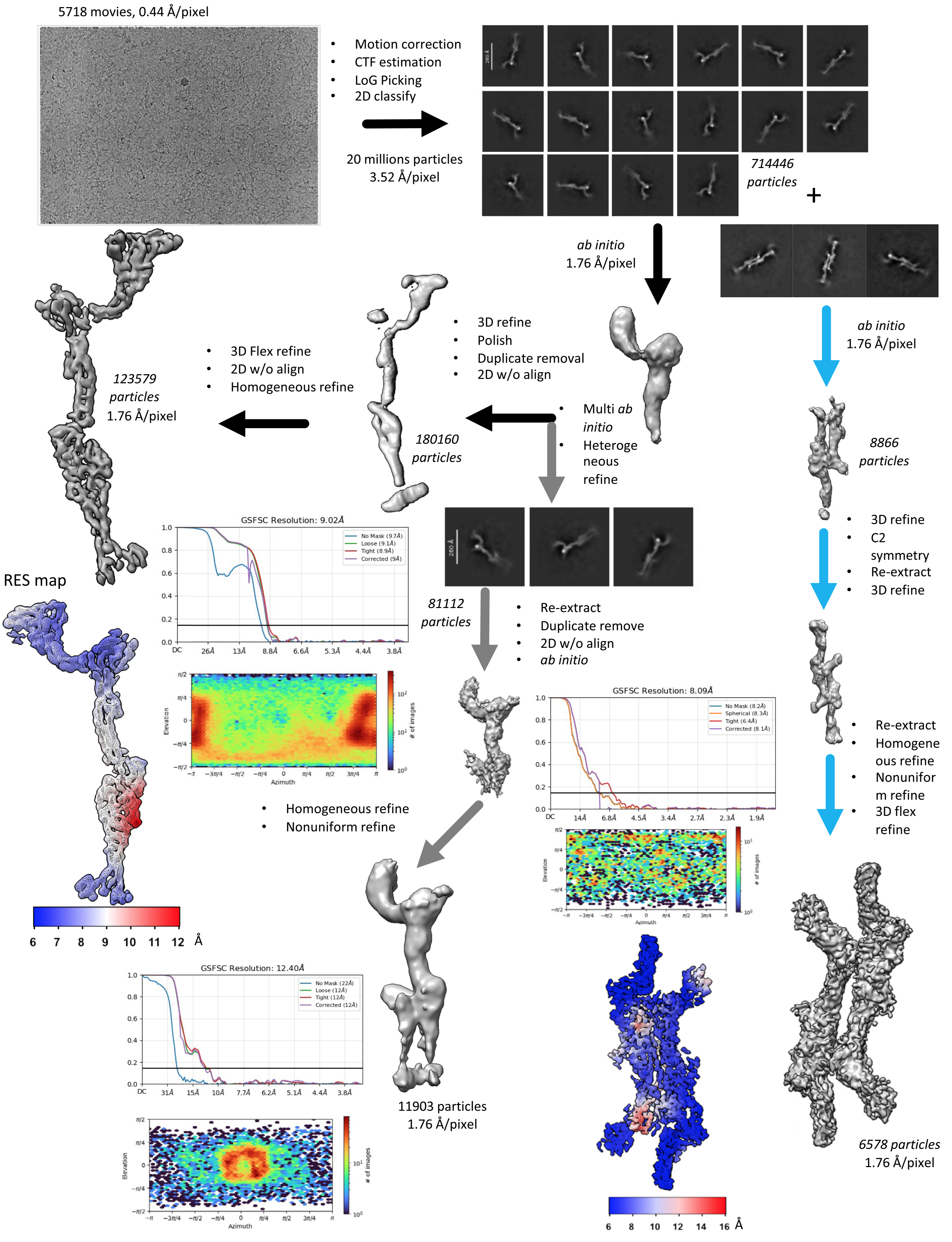

### Figure S3

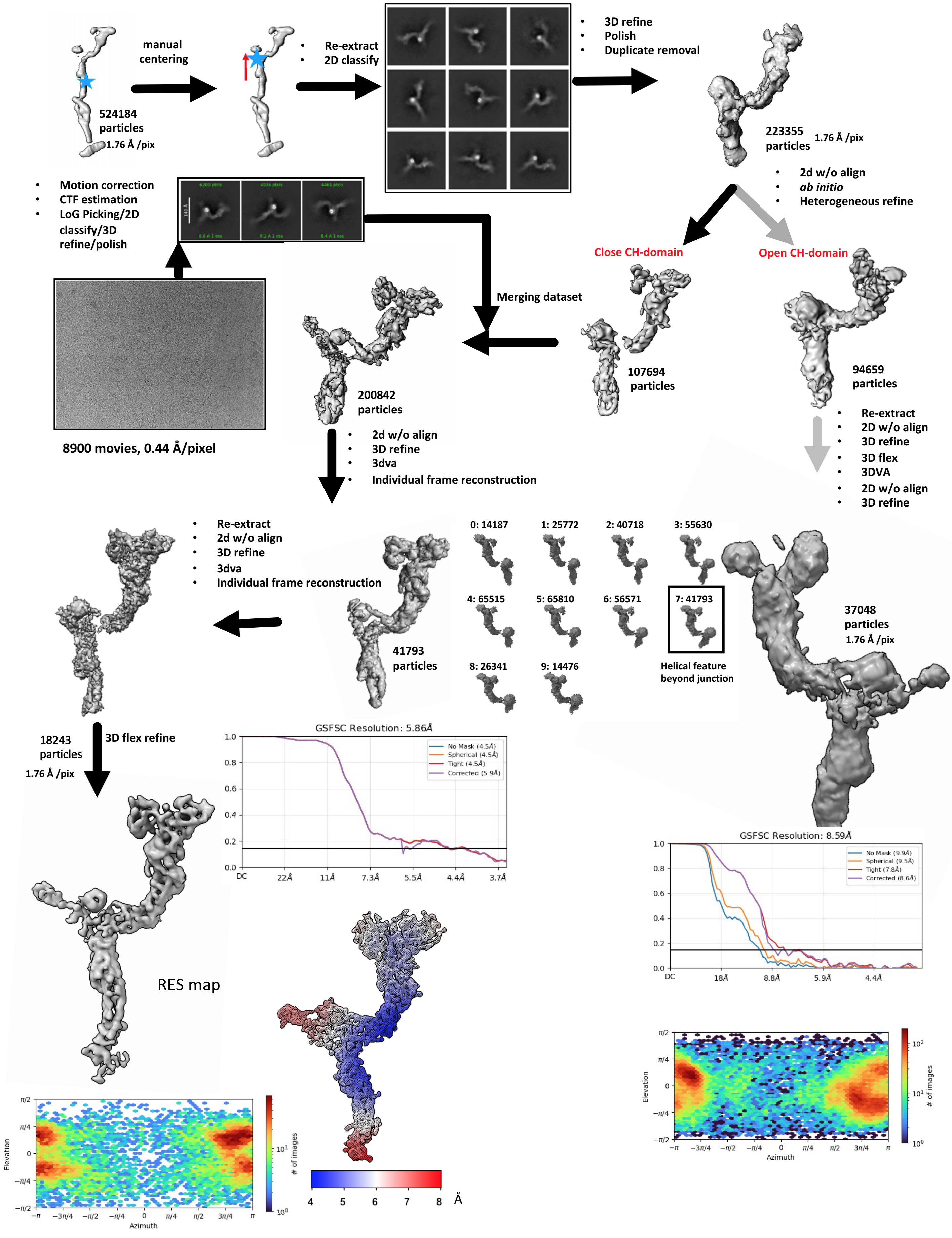

### Figure S4

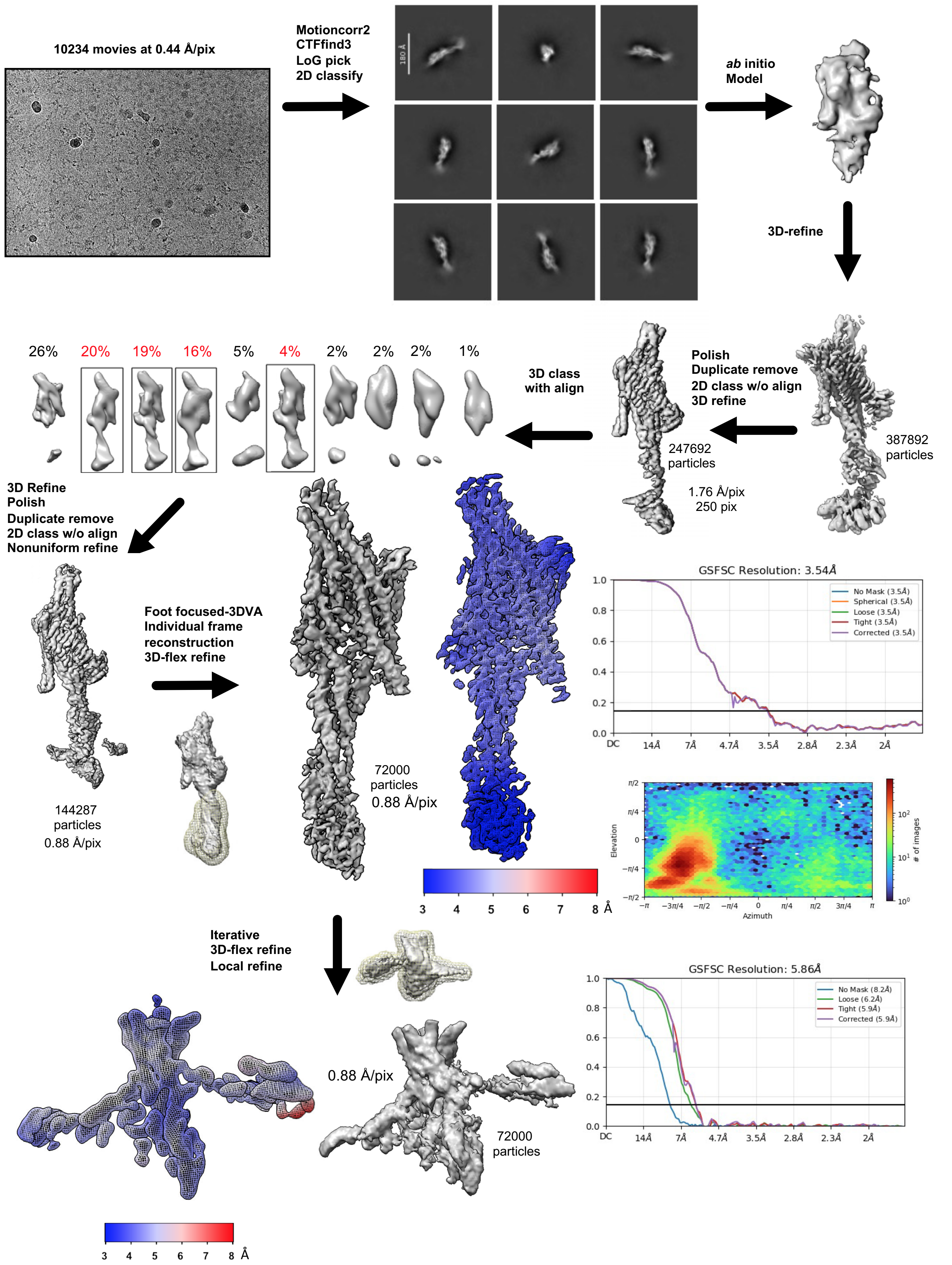

### Figure S5

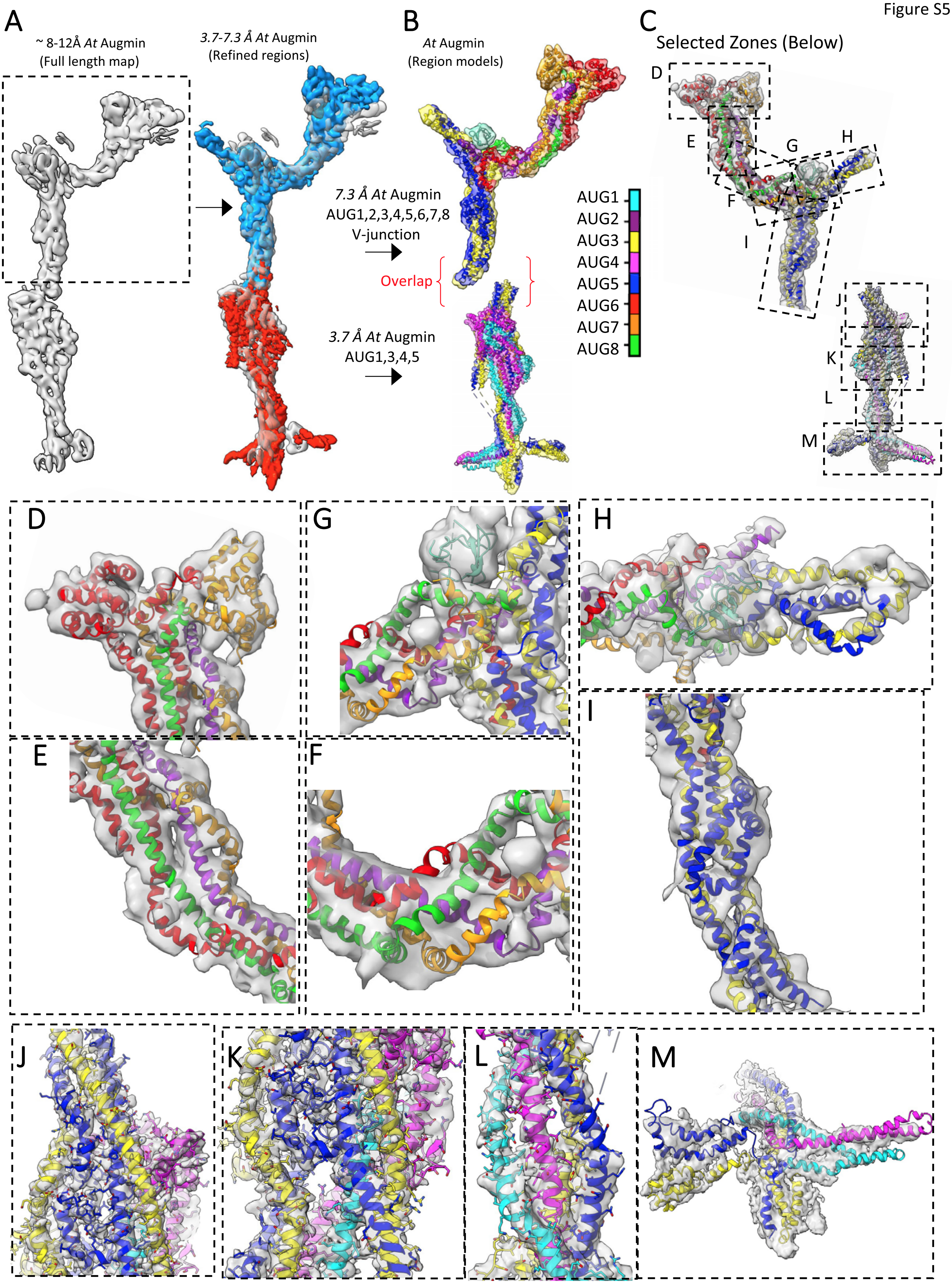

### Figure S6

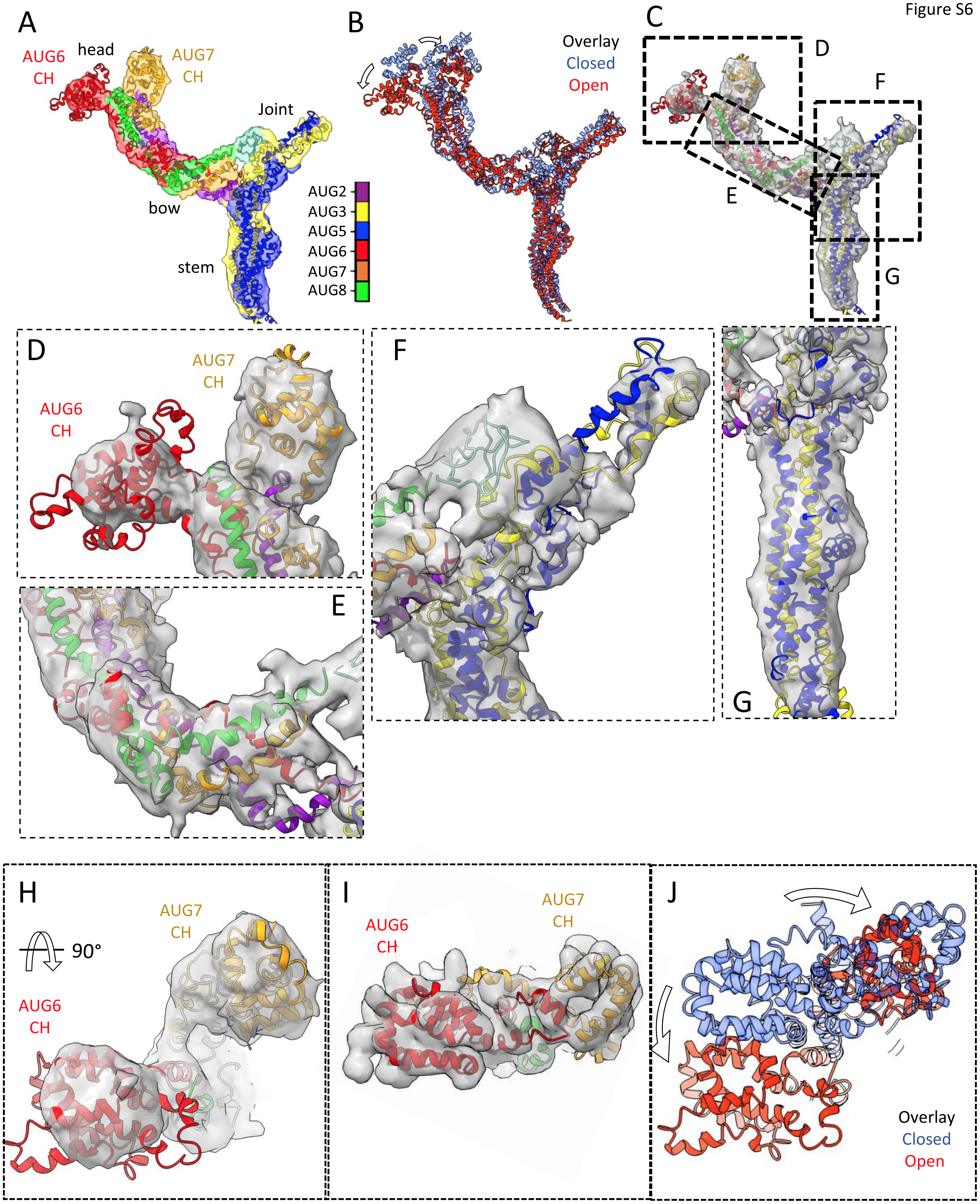

### Figure S7

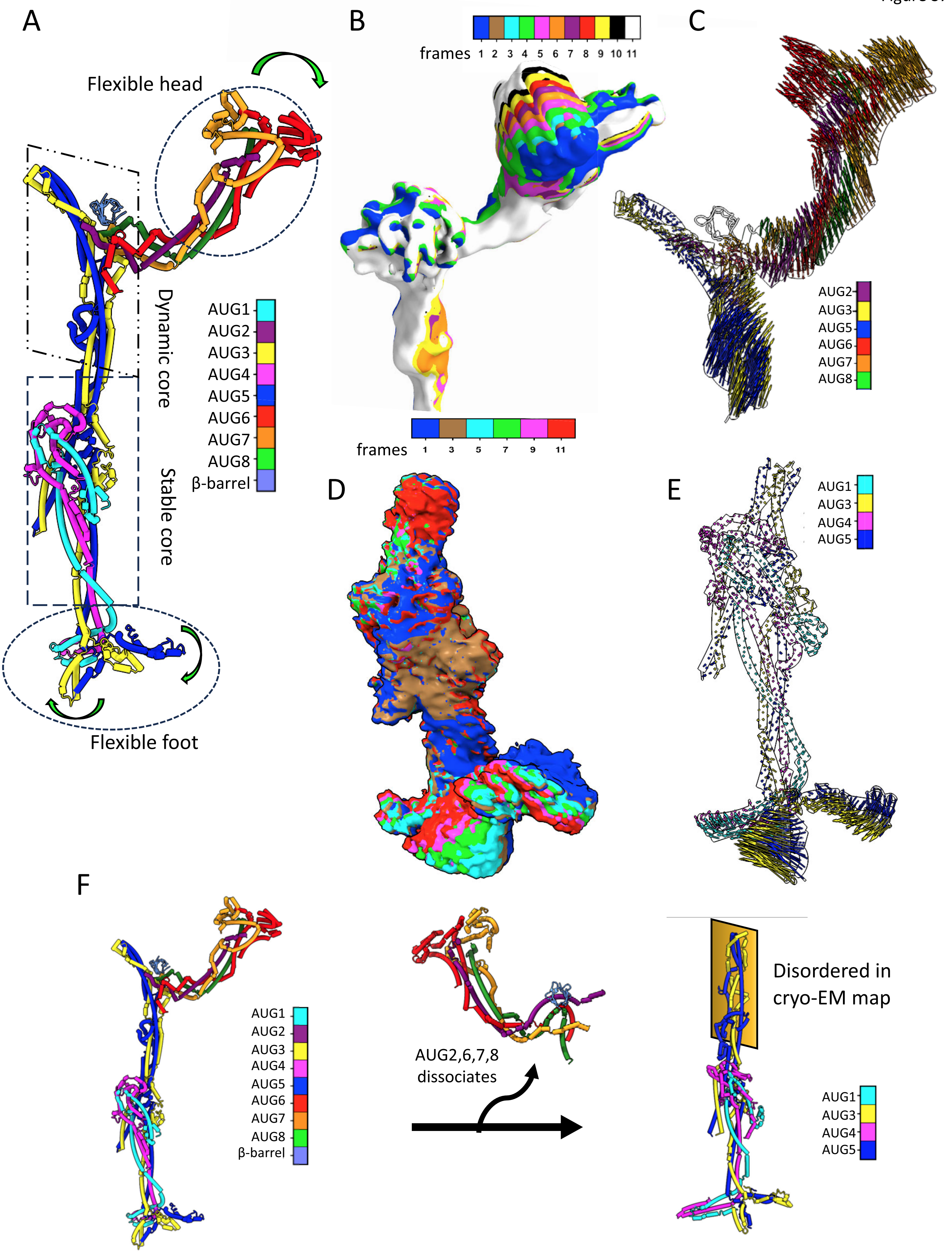

### Figure S8

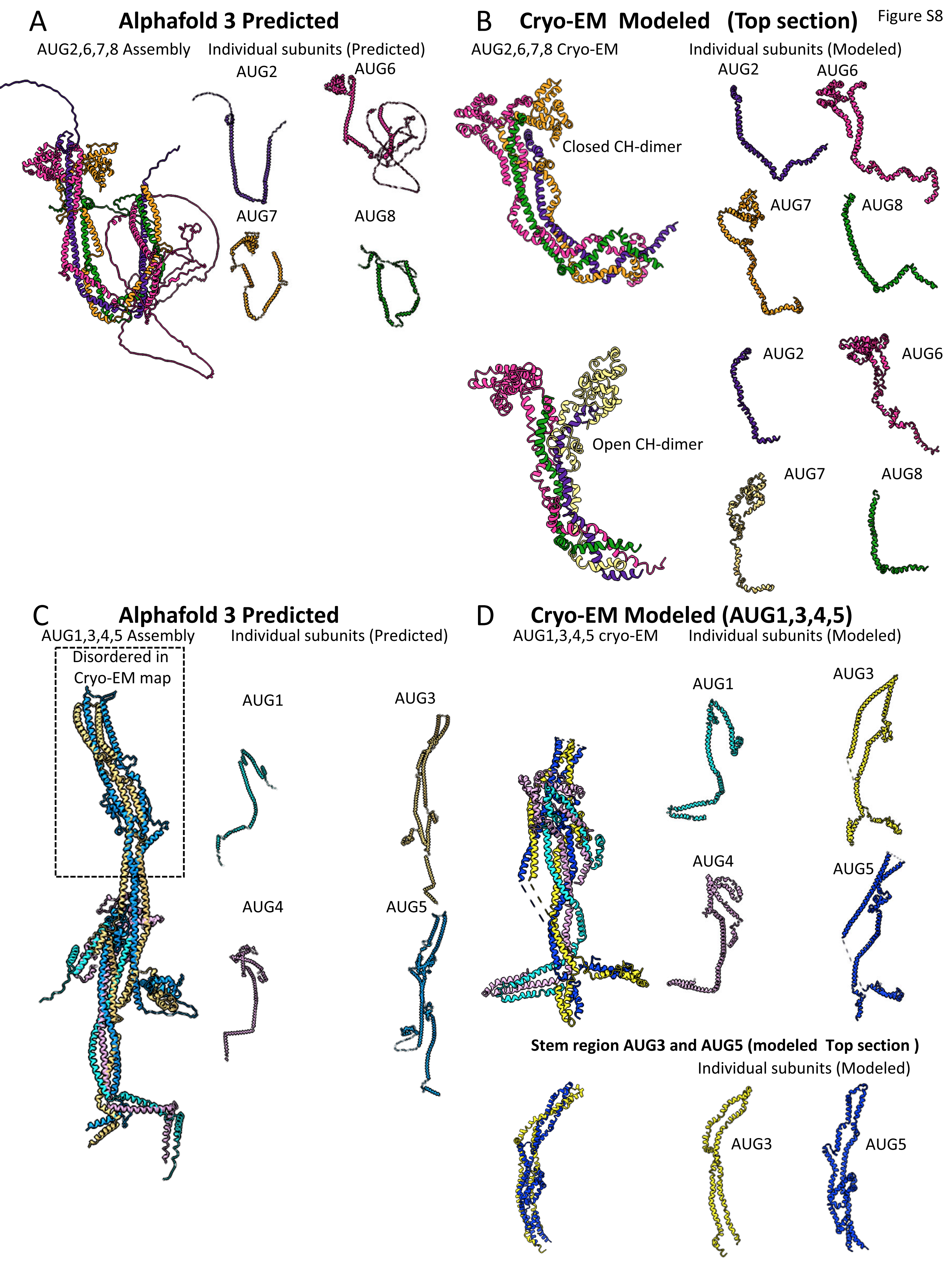

### Figure S9

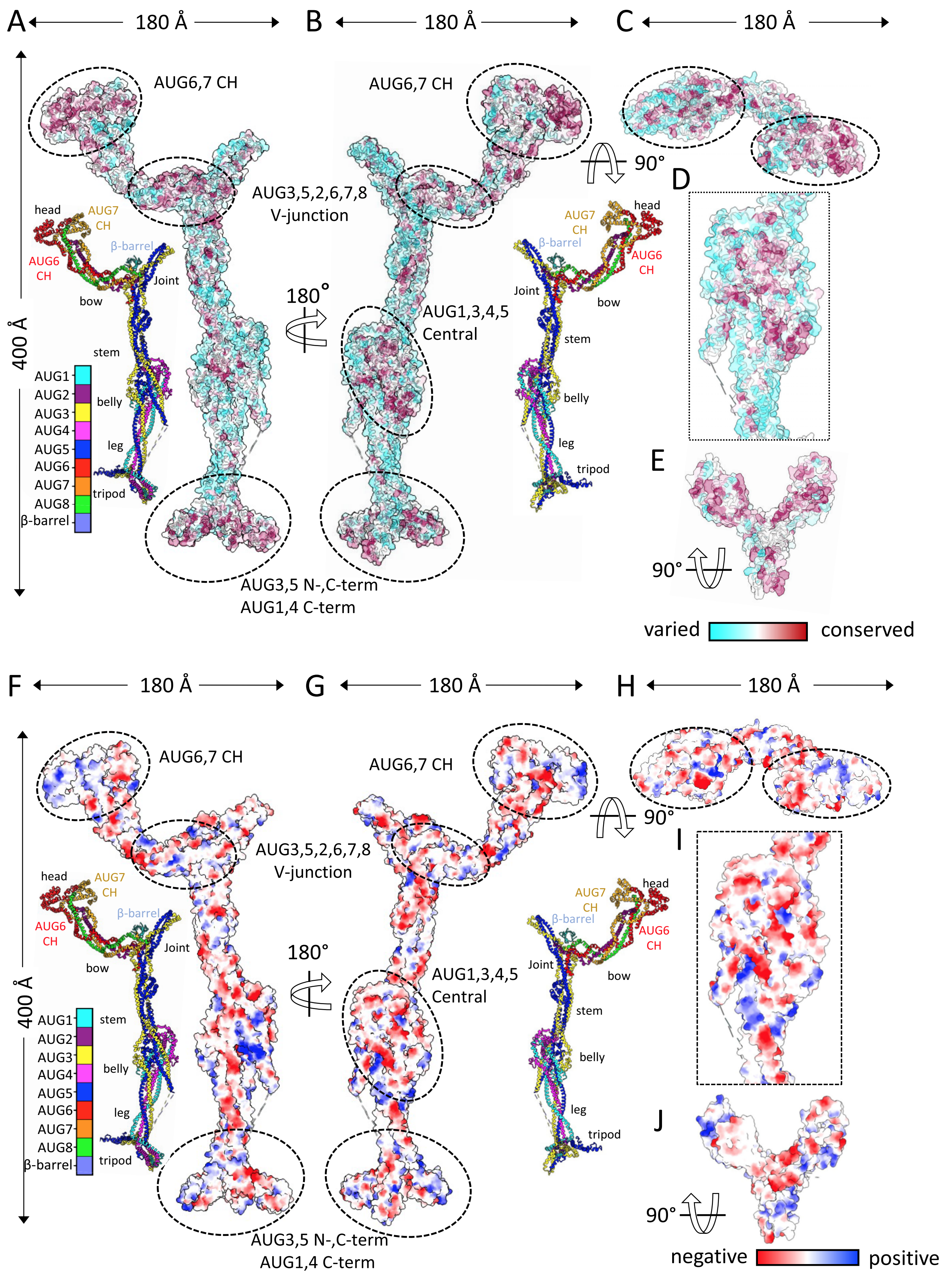

### Figure S10

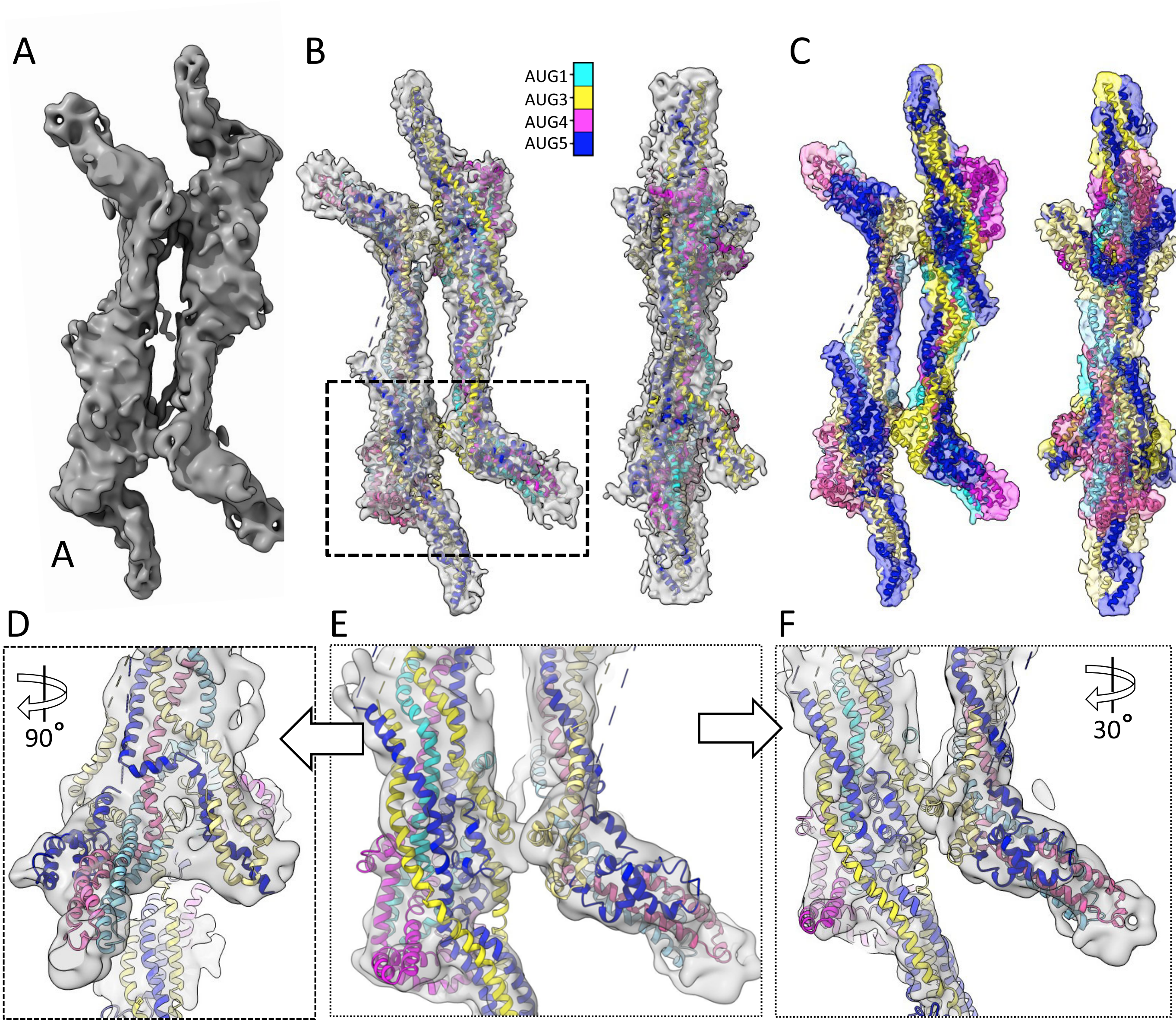

### Figure S11

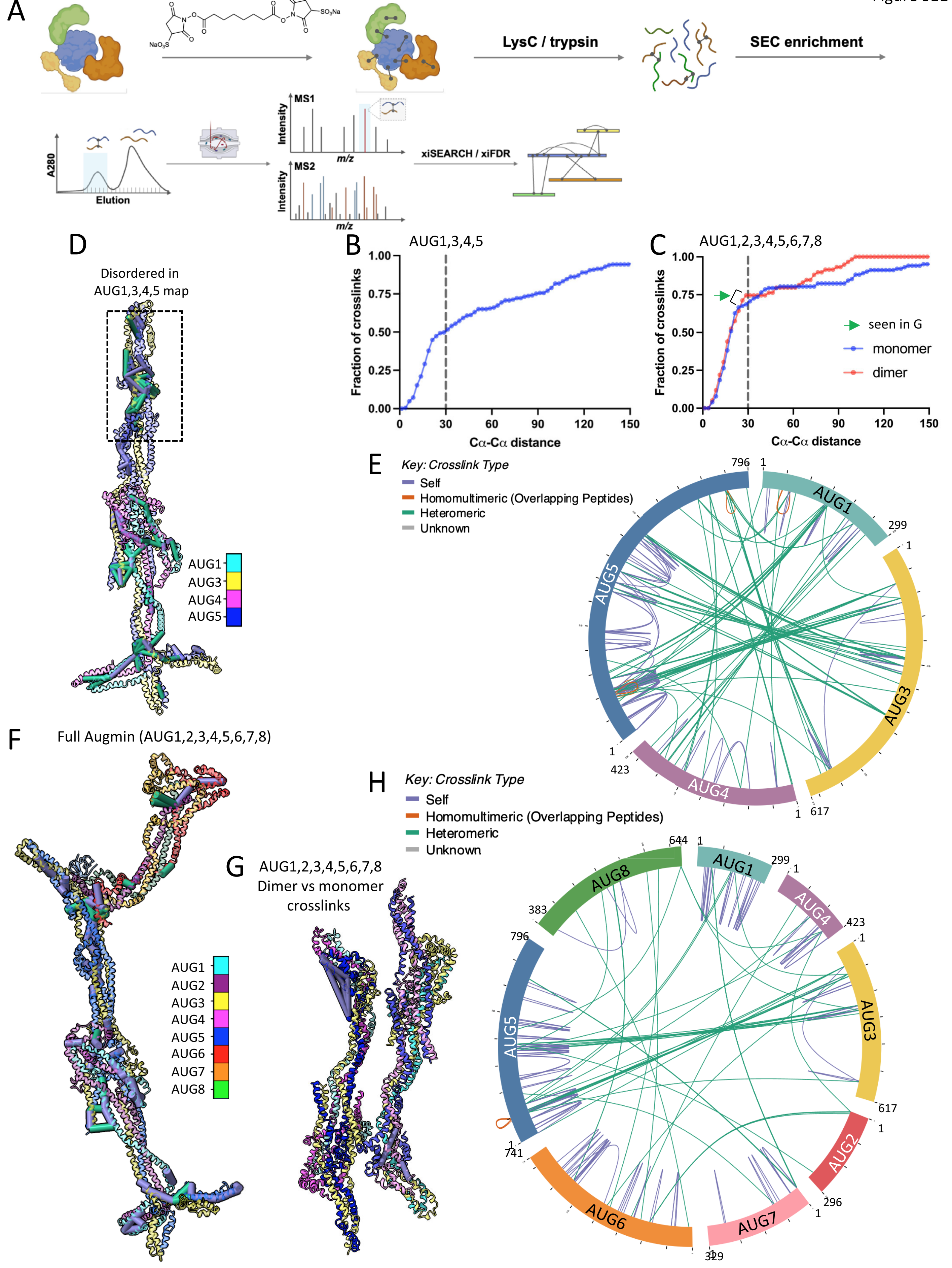

### Figure S12

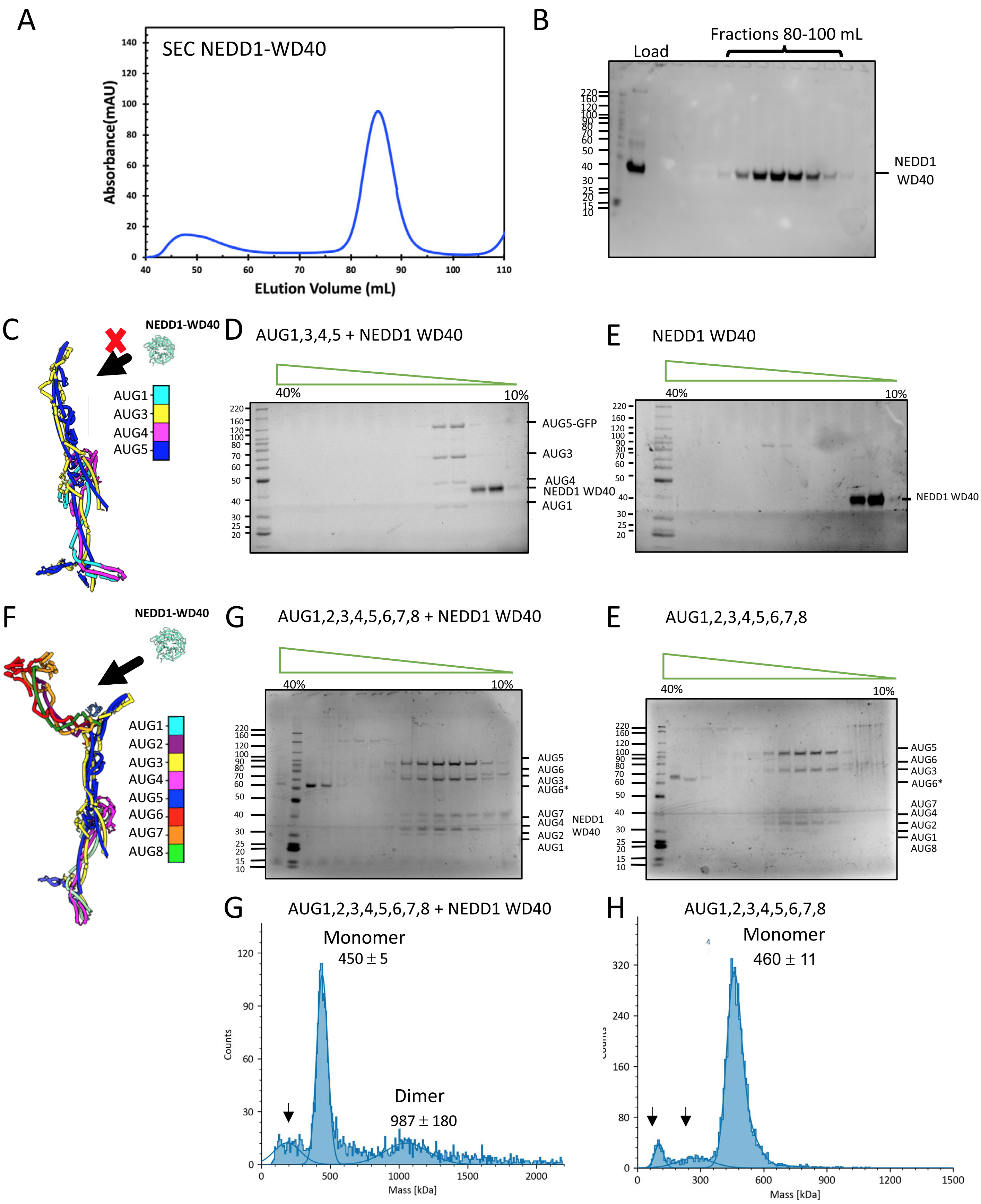

### Figure S13

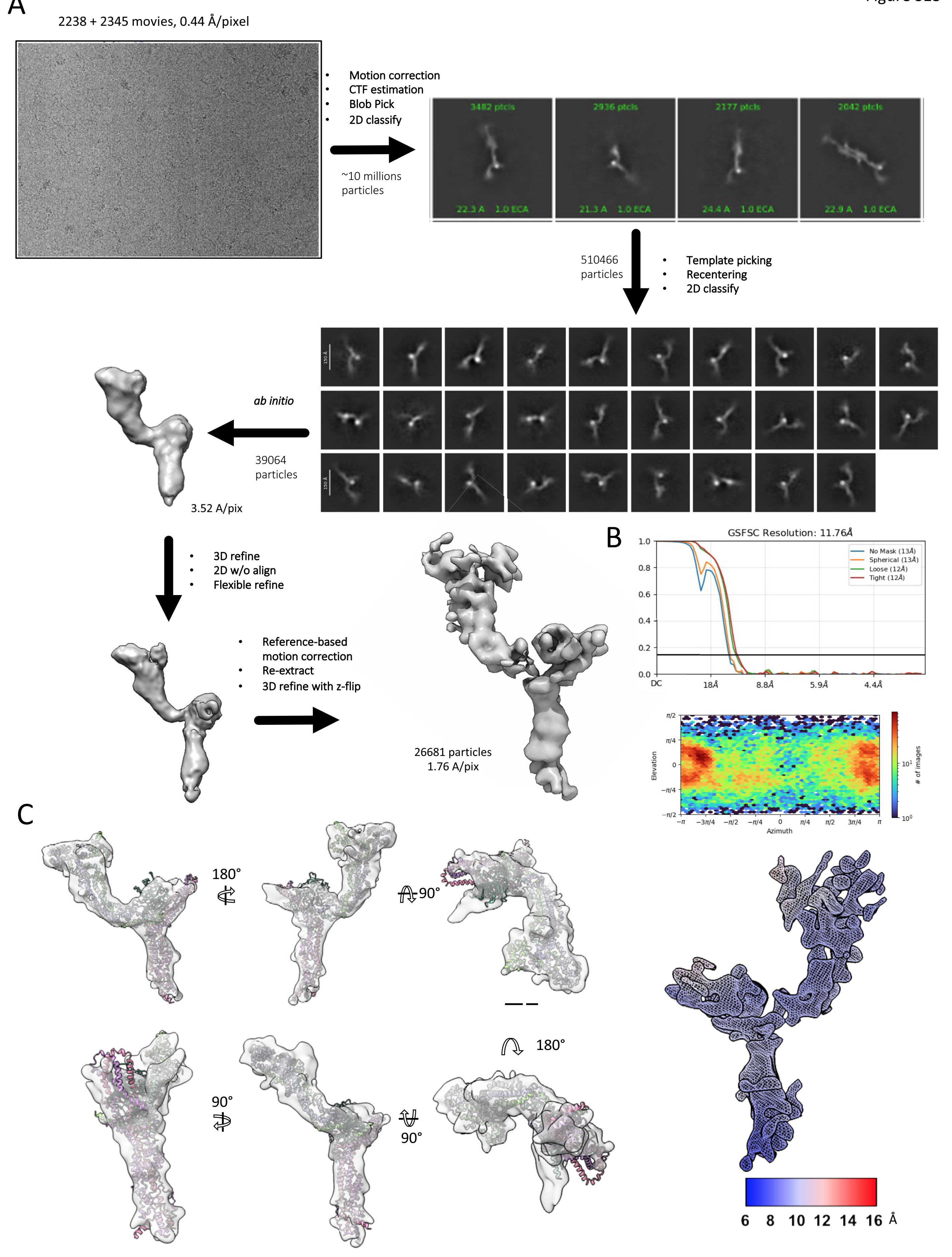

### Figure S14

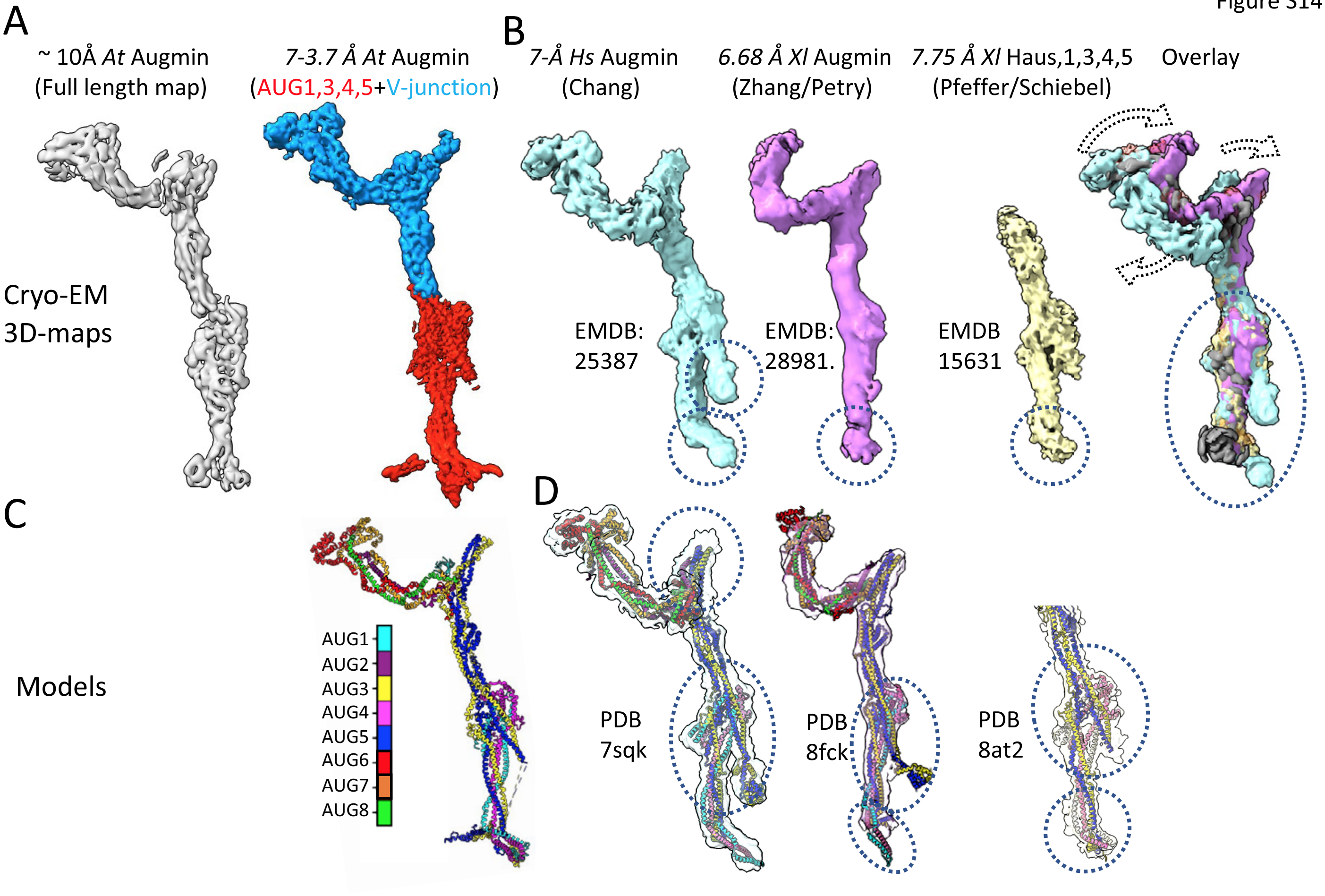
